# Exploring voltage-gated sodium channel conformations and protein-protein interactions using AlphaFold2

**DOI:** 10.1101/2024.10.15.618559

**Authors:** Diego Lopez-Mateos, Kush Narang, Vladimir Yarov-Yarovoy

## Abstract

Voltage-gated sodium (Na_V_) channels are vital regulators of electrical activity in excitable cells. Given their importance in physiology, Na_V_ channels are key therapeutic targets for treating numerous conditions, yet developing subtype-selective drugs remains challenging due to the high sequence and structural conservation among Na_V_ subtypes. Recent advances in cryo-electron microscopy have resolved most human Na_V_ channels, providing valuable insights into their structure and function. However, limitations persist in fully capturing the complex conformational states that underlie Na_V_ channel gating and modulation. This study explores the capability of AlphaFold2 to sample multiple Na_V_ channel conformations and assess AlphaFold Multimer’s accuracy in modeling interactions between the Na_V_ α-subunit and its protein partners, including auxiliary β-subunits and calmodulin. We enhance conformational sampling to explore Na_V_ channel conformations using a subsampled multiple sequence alignment approach and varying the number of recycles. Our results demonstrate that AlphaFold2 models multiple Na_V_ channel conformations, including those observed in experimental structures, states that have not been described experimentally, and potential intermediate states. Correlation and clustering analyses uncover coordinated domain behavior and recurrent state ensembles. Furthermore, AlphaFold Multimer models Na_V_ complexes with auxiliary β-subunits and calmodulin with high accuracy, and the presence of protein partners significantly alters both the modeled conformational landscape of the Na_V_ α-subunit and the coupling between its functional states. These findings highlight the potential of deep learning-based methods to expand our understanding of Na_V_ channel structure, gating, and modulation, while also underscoring the limitations of predicted models that remain hypotheses until validated by experimental data.

**Summary:** Lopez-Mateos et al.’s study demonstrates AlphaFold2’s potential to sample multiple states of human Na_V_ channels. Additionally, Na_V_ α-subunit interactions with β-subunits and calmodulin reshape Na_V_ α-subunit conformational landscape. This study reveals potential of deep learning methods to model structural diversity of ion channels.

## Introduction

Voltage-gated sodium (Na_V_) channels are essential regulators of electrical activity in excitable cells (Catterall, 2023). They open in response to membrane depolarization, allowing sodium ions to enter the cell and initiate transduction of sensory stimuli. This process is crucial for generating and propagating action potentials in neurons and muscle cells. Given their vital role in physiology, Na_V_ channels are targeted by numerous drugs for the treatment of various conditions such as epilepsy, pain or cardiac arrhythmia (Bagal et al., 2013; Noreng et al., 2021), and mutations in these channels have been linked to a wide range of diseases (Bennett and Woods, 2014; Meisler et al., 2021; Pan et al., 2021). The Na_V_ family in mammals contains nine subtypes, Na_V_1.1 through Na_V_1.9, with each exhibiting distinct tissue and cellular expression patterns that contribute to their specific physiological function (Catterall et al., 2020). Despite their pharmacological significance, achieving precise subtype selectivity—modulating a particular Na_V_ channel without affecting others— remains a significant challenge due to the high degree of overall sequence and structural conservation among the family members. Furthermore, due to the complex conformational states of Na_V_ channels, developing state-selective drugs that preferentially target channels in specific functional states is equally important for precisely regulating their electrical activity.

Recent advances in cryo-electron microscopy (CryoEM) have enabled the resolution of almost all human Na_V_ channels, except for Na_V_1.9 (Pan et al., 2021, 2019; Li et al., 2022; Pan et al., 2018; Li et al., 2021a; Fan et al., 2023; Huang et al., 2022a; b). All mammalian Na_V_ channels share a conserved architecture, with a core α-subunit and auxiliary β-subunits (Catterall, 2023). The Na_V_ α-subunit consists of four homologous domains, each comprising six transmembrane segments (S1-S6). The first four segments (S1-S4) form the voltage-sensing domains (VSDs), with the gating charges located in the S4 segment, while S5 and S6 form the pore domain (PD). The VSDs are positioned outwards from the pore, while the PD forms the central ion-conducting pore. Auxiliary Na_V_ β-subunits (β1-β4) are transmembrane proteins with extracellular immunoglobulin domains that interact with the Na_V_ α-subunit to modulate various channel properties (Namadurai et al., 2015). These include surface localization, kinetic behavior, and clustering, making them crucial regulators of cellular excitability. Additionally, Na_V_ channels are modulated by calcium through calmodulin (CaM) binding (Wu and Hong, 2021). Beyond endogenous protein partners, NaV channels are also targeted by exogenous modulators such as peptide toxins from animal venoms, for which structural data have provided additional insights into their mechanisms of action (Pan et al., 2019; Clairfeuille et al., 2019; Xu et al., 2019; Shen et al., 2019; Wisedchaisri et al., 2020). This growing body of structural and functional data has remarkably advanced our understanding of the Na_V_ conformational cycle and regulation (Catterall et al., 2020b). In the resting state, the channels are closed with deactivated VSDs, where S4 segments sit in the ‘down’ position. Depolarization drives the positively charged S4 segments upward, activating the VSDs and opening the activation gate (AG), a hydrophobic constriction on the intracellular pore, to allow sodium influx. Activation of VSDIV is coupled to channel fast inactivation: the Ile-Phe-Met (IFM) motif, stabilized beneath deactivated VSDIV, is released upon S4 movement and binds to the pore domain, closing the AG and stopping sodium flow (Capes et al., 2013; Clairfeuille et al., 2019). At the extracellular side, the selectivity filter (SF), defined by the Asp-Glu-Lys-Ala (DEKA) motif, ensures sodium selectivity, and its conformational changes have been implicated in the less understood process of slow inactivation (Chen et al., 2024).

Recent experimental structures have provided crucial insights into various states of the Na_V_ channel conformational cycle. Previously, the inability to control membrane voltage during structural determination presented a significant challenge in capturing channels in specific states. Researchers have employed strategies to overcome this limitation, including introducing mutations (Jiang et al., 2021), using peptide toxins (Clairfeuille et al., 2019), or applying chemical crosslinking (Lee and MacKinnon, 2019) to stabilize ion channels in particular conformational states. Recently, MacKinnon’s lab developed an approach using lipid membrane vesicles with a voltage difference across the membrane to solve cryoEM structures of Eag and KCNQ1 channels in different states (Mandala and MacKinnon, 2022, 2023). Despite this progress, fully resolving the conformational landscape of Na_V_ channel gating and modulation at high resolution remains challenging, hindering the development of subtype-selective, state-dependent Na_V_ channel modulators with therapeutic potential.

Computational methods have become powerful tools to advance our understanding of Na_V_ channel structure, gating, and modulation. The advent of deep learning-based methods for protein structure prediction has significantly enhanced our ability to accurately model protein structures (Pakhrin et al., 2021). Our group demonstrated that AlphaFold2 (Jumper et al., 2021), RoseTTAFold2 (Baek et al., 2021), and ESM Fold (Lin et al., 2023) achieved unprecedented and remarkable accuracy in modeling full Na_V_ channels, with AlphaFold2 outperforming the other methods (Nguyen et al., 2024). AlphaFold2 extends its functionality with AlphaFold Multimer (Evans et al., 2022), designed to predict protein complexes by accurately modeling protein-protein interactions. However, this methodology has yet to be explored for modeling interactions between Na_V_ channels and their protein partners, such as auxiliary subunits or CaM. Additionally, recent studies have shown that AlphaFold2 can model multiple conformational states of proteins by adjusting generation and use of multiple sequence alignment (MSA) to generate models (del Alamo et al., 2022; Wayment-Steele et al., 2023; Monteiro da Silva et al., 2024). Importantly, these studies demonstrate that multiple states can be modeled, although the resulting distribution of states does not necessarily reflect the true energetic landscape of the protein.

In this work, we asked five main research questions: (i) Does applying a subsampled AlphaFold2 approach and generating multiple models of a full Na_V_ channel allow sampling multiple distinct conformations beyond a single state? (ii) What is the range of conformational states that can be sampled with this approach, and how do the resulting state distributions correlate across functional regions? (iii) Which factors influence the states obtained? (iv) Can AlphaFold Multimer reproduce experimentally observed complexes of the Na_V_ α-subunit with its native partners? (v) Does the presence of these partners alter the modeled conformational landscape of the α-subunit? To address these questions, we modeled all human Na_V_ channel subtypes with AlphaFold2 using a subsampled approach (Monteiro da Silva et al., 2024) to promote conformational variability, generated hundreds of models per case, evaluated the diversity of the generated conformational states, and analyzed correlations between conformations of different Na_V_ channel regions. In parallel, we built models of the Na_V_ α-subunits with β-subunits and CaM using AlphaFold Multimer, and evaluated the accuracy of these predictions against available experimental structures of the complexes. Our findings demonstrate that AlphaFold2 can model multiple Na_V_ channel states observed in experimental structures, as well as those that have not been observed, and reveal potential intermediate states. Moreover, we show that AlphaFold Multimer accurately models the structure of Na_V_ α-subunit complexes with protein partners and demonstrate that the presence of these partners profoundly reshapes the modeled conformational ensemble of the Na_V_ α-subunit. While our results illustrate the potential of deep learning methods in studying Na_V_ channel conformations, it is essential to recognize that these predictions remain hypothetical, may include inaccuracies or hallucinated features, and must be rigorously validated by experimental data before firm mechanistic conclusions can be drawn.

## Materials and Methods

### General workflow

In this study, we followed a systematic pipeline for the modeling and analysis of NaV channels and their protein partners. A total of twenty cases were examined. These included individual α-subunits of NaV1.1 (Acc: P35498), NaV1.2 (Acc: Q99250), NaV1.3 (Acc: Q9NY46), NaV1.4 (Acc: P35499), NaV1.5 (Acc: Q14524), NaV1.6 (Acc: Q9UQD0), NaV1.7 (Acc: Q15858), NaV1.8 (Acc: Q9Y5Y9), and NaV1.9 (Acc: Q9UI33), as well as the non-voltage-gated NaX channel (Acc: Q01118). In addition, four cases corresponded to NaV1.7 modeled in complex with each of the four auxiliary β-subunits β1 (Acc: Q07699), β2 (Acc: O60939), β3 (Acc: Q9NY72), and β4 (Acc: Q8IWT1), and another four cases corresponded to NaV1.1 paired with the same set of β-subunits. Finally, we examined complexes of NaV1.2 and NaV1.5 with calmodulin (CaM; Acc: P0DP23). All the sequences used in this study can be found in Table S1. For each case, the general procedure involved sequence retrieval from UniProt, construction of multimeric complexes when required, structural modeling using the subsampled MSA approach described below, and subsequent analysis of the resulting ensembles. From these ensembles, we defined and calculated a series of distance-based coordinates that capture the conformational states of key structural elements, alongside both global and region-specific predicted local distance difference test (pLDDT) values to assess model confidence. This framework enabled us to characterize the conformational diversity and reliability of AlphaFold2 predictions across all study cases.

To ensure reproducibility, all input sequences, scripts, command lines, and analysis codes required to replicate this workflow step by step are provided in a dedicated GitHub repository with detailed instructions. Additionally, all the generated models have been deposited into the Dryad database (see Data Availability).

### Model generation with subsampled AlphaFold

All models in this study were generated using colabfold-batch 1.5.0 (localcolabfold) (Mirdita et al., 2022) with a subsampled MSA (Monteiro da Silva et al., 2024). The execution was performed with the following general options:

*colabfold_batch --num-models 5 --model-type auto --msa-mode mmseqs2_uniref_env \*

--num-seeds 20 \

--templates --max-seq 256 --max-extra-seq 512 \

--num-recycle 6 --save-recycles nav17alphafull.fasta outfiles-6r-256-512-sr

The key aspect that makes this colabfold execution a MSA subsampled implementation is the setting of two parameters: --max-seq and --max-extra-seq. The --max-seq parameter defines the maximum number of sequences randomly selected from the original generated master MSA. The target sequence is always included. Then, the remaining sequences are clustered around the selected sequences using Hamming distance. Finally, the cluster centers (the --max-seq sequences chosen originally) are used alongside a sample from each cluster (as defined by the --max-extra-seq parameter) for the structure prediction network. Previous studies have shown that adjusting these parameters, particularly by reducing their values, can increase the diversity of conformational sampling (del Alamo et al., 2022). By randomly selecting fewer sequences from the MSA, subsampling alters the co-evolutionary information provided to AlphaFold2. This, in turn, enables the prediction network to explore a broader range of physiologically relevant conformations rather than converging on a single ground state structure. In this study, we used values of 256 for --max-seq and 512 for --max-extra-seq, as these settings enhanced conformational diversity without compromising model accuracy in the original study (Monteiro da Silva et al., 2024). A total of 100 models were generated for each study case, using 6 recycles. Intermediate models were saved at each recycle step, resulting in 7 models per generated structure, ranging from recycle 0 to recycle 6, making a total of 700 models per test case (see Data Availability).

When modeling multimeric complexes (e.g., the Na_V_ α-subunit together with an auxiliary β-subunit or calmodulin), no changes in execution options were required. The only modification consisted of preparing the input FASTA file by placing the sequences of the two partners on the same line, separated by a colon “:”. This syntax instructs ColabFold to treat them as interacting partners in separate chains within the same modeling run.

In cases where we evaluated the effect of using state-specific experimental structures as templates, the subsampled AlphaFold2 execution was run in custom template mode. This only requires adding the option *--custom-template-path $PATH* to the execution command shown above, where *$PATH* specifies a directory containing one or more PDB files used as custom templates. All other parameters remained identical to the general execution described above.

### Model Analysis

All models were visually analyzed using UCSF ChimeraX (Goddard et al., 2018). ChimeraX was also used to calculate the root mean square deviation (RMSD) values after superimposing each model with the experimental structure of reference using the Matchmaker tool. Distance coordinates and subset pLDDT values were automatically calculated with custom Python scripts utilizing the PyRosetta package (Chaudhury et al., 2010). All other descriptive analyses, distribution calculations, and figure generation were conducted with custom Python scripts. Pore analysis of the channels was conducted using the MOLEonline web interface (Pravda et al., 2018). All scripts to conduct the analyses described in this study can be found in the dedicated GitHub repository (see Data Availability).

### Correlations and Clustering Analysis

Pairwise correlations between all state coordinates were calculated to assess the relationships among the conformational descriptors extracted from the modeled ensembles. For each pair of coordinates, we computed Spearman’s rank correlation to capture monotonic relationships independent of linearity, and the associated p-values to evaluate statistical significance. Additionally, mutual information was estimated using a non-parametric k-nearest-neighbor approach to detect potential non-monotonic dependencies.

To identify major conformational patterns shared across all models of the nine Na_V_ channel types, we constructed a joint feature matrix in which each row corresponds to a single model and each column represents a numerical descriptor of that model. These descriptors included both the distance-based state coordinates and the global and subset pLDDT values corresponding to specific functional regions. This combination allowed the clustering to account simultaneously for conformational geometry and model confidence across subdomains. The resulting multi-channel dataset was then subjected to Leiden community detection (Traag et al., 2019) on a 500-nearest-neighbor graph with a resolution parameter of 0.5, enabling the identification of densely connected groups of models that represent recurrent conformational states. The resulting cluster assignments were visualized using two-dimensional UMAP embeddings computed on the same feature set, allowing qualitative inspection of cluster separations.

## Results

### Defining distance coordinates to identify channel states

Na_V_ channels are complex molecular machines that exhibit dynamic transitions between distinct conformational states, influenced by voltage, regulatory protein partners, ion concentrations, lipids, and other factors like phosphorylation and glycosylation. To evaluate AlphaFold2’s ability to sample this conformational diversity, we focused on key regions of the Na_V_ α-subunit whose structural dynamics have been extensively characterized (Fig. 1). We defined specific distance coordinates for each of these Na_V_ α-subunit regions that could be calculated across all generated models to assess different conformational states (Table S2).

**Figure 1.**
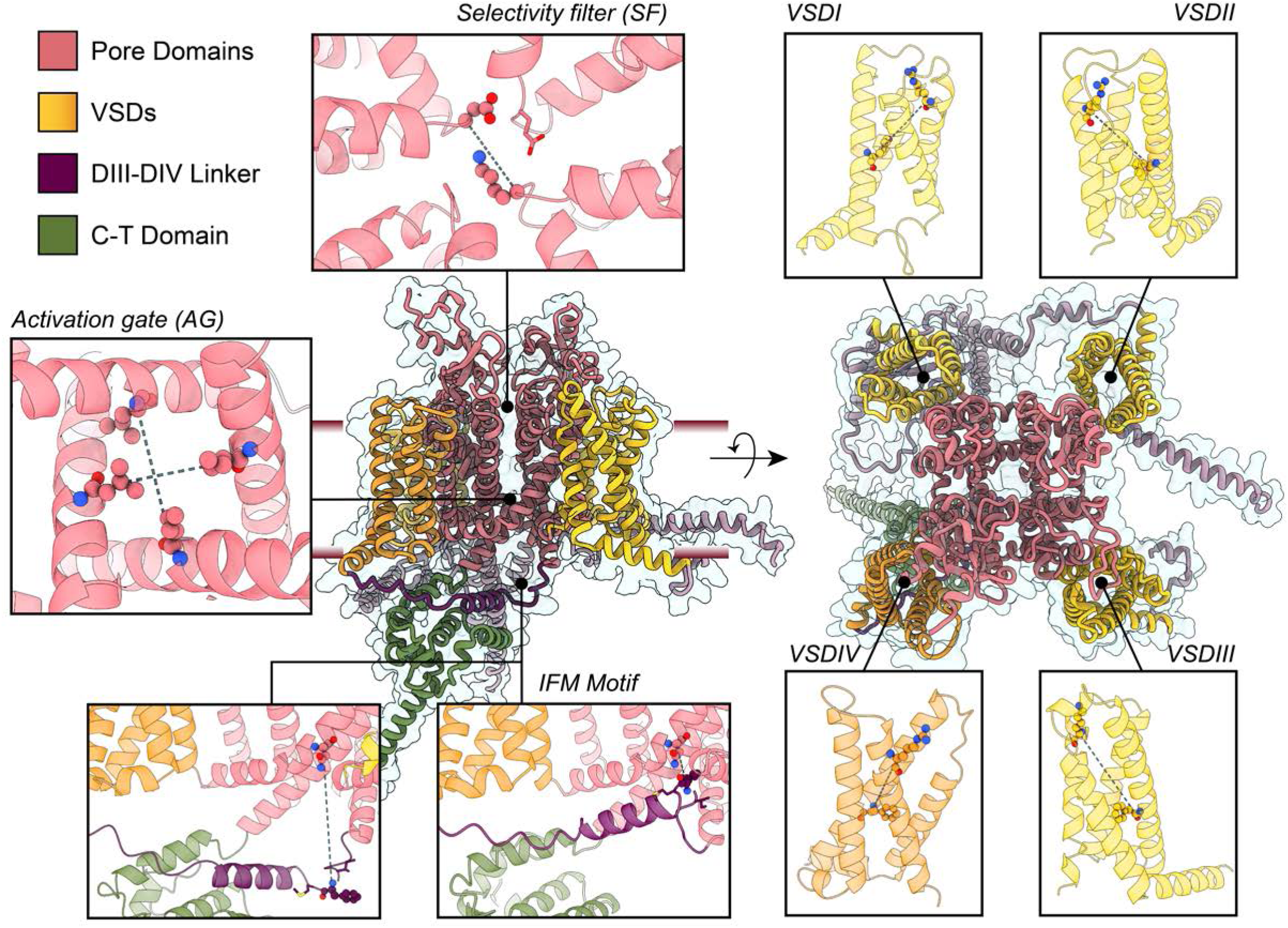
Selected distances to use as coordinates to identify channel states. We focused on seven different regions of Na_V_ channels with known conformational dynamics: the four VSDs, the SF, the AG and the IFM motif. The calculated distances are illustrated in the figure with dashed lines.

For the VSDs, we measured the distance between the α-carbon of the first gating charge in the S4 segment (GC1-S4) and the α-carbon of the residue forming the hydrophobic constriction site in the S2 segment (HC-S2). Larger values of this distance (GC1-S4 – HC-S2) indicate an activated or “up” VSD state, while smaller values would represent a deactivated or “down” VSD state. In the activation gate (AG), we calculated the distances between the α-carbons of opposing residues that form the hydrophobic intracellular gate of the channel (named AG1 and AG2, for S6_I_-S6_III_ and S6_II_-S6_IV,_ respectively). This allows us to distinguish whether the activation gate is open or closed. We also measured the distance between the α-carbon of the phenylalanine in the IFM motif and the α-carbon of the aspartic acid within the IFM binding site in the pore domain (IFM – PD distance). A low value for this distance indicates that the IFM motif is bound, placing the channel in a fast-inactivated state. Additionally, we assessed the distance between the α-carbons of the lysine and aspartic acid residues in the DEKA motif of the selectivity filter (SF), naming this distance SF-D – K. Variations in this distance indicate changes in the dilation of the SF, which may represent different conformational states associated with slow inactivation (Vilin and Ruben, 2001). To assess whether the generated models encompass the range of experimentally observed conformations, we calculated these same distances in available structures of human Na_V_ channels, as well as specific non-human Na_V_ channels representing distinct states, for comparison (Table S3).

After defining these distance coordinates across all human Na_V_ channels, we proceeded with the modeling using ColabFold (Mirdita et al., 2022). We generated 100 models per full channel using a subsampled MSA (Monteiro da Silva et al., 2024) with six recycles, saving intermediate recycles for analysis (see Methods). Note that “recycles” in AlphaFold modeling refers to iterative steps of feeding the predicted structure back into the neural network to refine and improve the model’s accuracy. Although we initially analyzed each region of interest separately, the models included the full channel sequence, enabling us to subsequently evaluate the coupling between the states of different regions. We modeled all human voltage-gated Na_V_ channels (hNa_V_1.1 to hNa_V_1.9) and the non-voltage-gated hNa_X_ channel with a Na_V_-like architecture (Noland et al., 2022). Modeling hNa_X_ allowed us to evaluate whether the state distributions of AlphaFold models of hNa_V_ channels versus the hNa_X_ channel reflected the distinct functional features of the Na_X_ channel. In addition to calculating the distance coordinates described above, we evaluated the predicted local distance difference test (pLDDT) values for all models as the prediction confidence metric. Generally, pLDDT values above 90 indicate very high confidence, 70-90 indicate good confidence, 50-70 indicate low confidence, and below 50 indicate very low confidence. While AlphaFold provides a global pLDDT score as a measure of overall model confidence, since this is a metric reported at the residue backbone level, we focused on subset pLDDT values for specific regions of interest: the VSDs, the four residues comprising the AG, the DEKA motif in the SF, and the three residues forming the IFM motif. Furthermore, we calculated an adjusted-global pLDDT score that excluded the large unstructured intracellular loops. These regions typically receive low and variable pLDDT values, which can distort the overall score and lead to misleading variations in the global pLDDT. By focusing only on well-structured regions of the channel, the adjusted score provides a more accurate reflection of the overall model’s confidence.

### AlphaFold samples multiple VSD states with varying degrees of gating charge translocations

We first focused on the states of the VSDs (Fig. 2). Overall, we observed that various conformational states were sampled, with the extent of this sampling varying across VSDs. Notably, within each VSD, the state distribution was consistent across hNa_V_ subtypes, meaning that for a given VSD, the sampled states were similar among the nine different hNa_V_ subtypes. The remarkable exception was hNa_X_, which is not voltage-dependent, and we expected to observe a distinct state distribution. Indeed, hNa_X_’s state distributions stand out across all four VSDs, aligning with the fact that the VSD-like domains of Na_X_ are not able to sense voltage and therefore adopt different conformational states compared to Na_V_ channels. Below, we analyze each VSD in detail.

**Figure 2.**
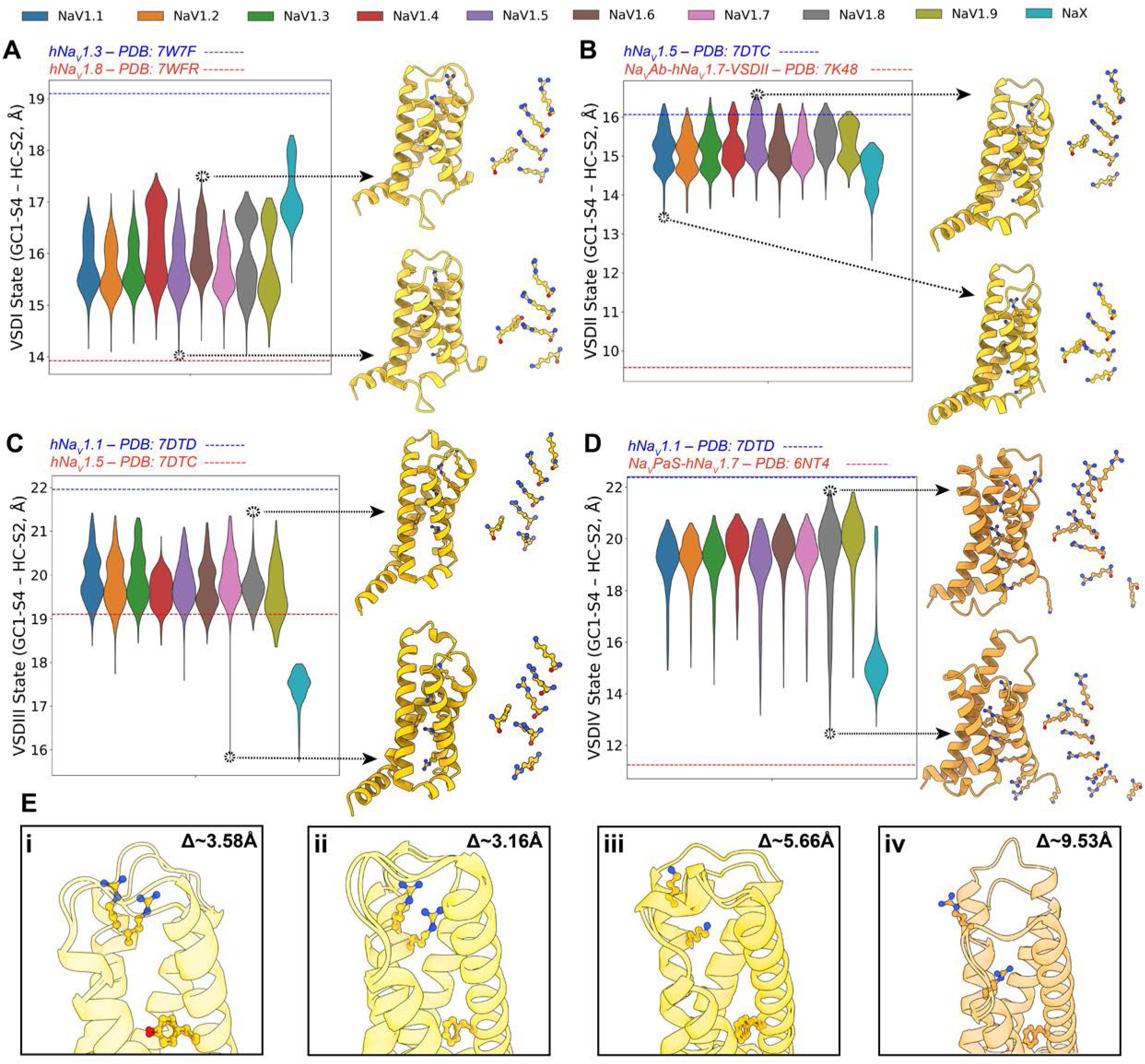
Distribution of VSD states in generated AlphaFold models of hNa_V_ channels. (A-D) Distribution of the GC1-S4 – HC-S2 distances for the four VSDs across the nine hNa_V_ channel subtypes plus the hNa_X_ channel. Blue dashed lines mark the largest distance observed in experimental structures for that coordinate, while the red dashed lines represent the lowest; this provides an idea of the observed range of states in experimental structures. The VSD models with the largest and smallest distance coordinates for each VSD are shown, highlighting the relative positioning of the gating charges in relation to the hydrophobic constriction site. (E) Superimposition of the most activated and deactivated models for each of the VSDs shown in panels A-D, highlighting the difference in the distance coordinates between the most activated and deactivated models.

In the VSDI, we observed bimodal distributions of the GC1-S4 – HC-S2 distances, which fall within the range of conformational states seen in experimental structures (Fig. 2 A). The most activated state was observed in the experimental structure of hNa_V_1.3 (PDB: 7W7F) (Li et al., 2022) with the GC1-S4 – HC-S2 distance of 19.1 Å, while the most deactivated state was seen in hNa_V_1.8 (PDB: 7WFR) (Huang et al., 2022b) with the GC1-S4 – HC-S2 distance of 14.3 Å. In our models, the shortest GC1-S4 – HC-S2 distance was found in a model of hNa_V_1.5 at 14.0 Å, and the longest in a model of hNa_V_1.6 with a distance of 17.6 Å. Visual comparison of these models, specifically the relative positions of the gating charges in the S4 segment, indicates that these structures represent two distinct VSDI states: an activated state has three of the four gating charges in the S4 segment positioned above the hydrophobic constriction site in the S2 segment, while a partially deactivated state shows only two gating charges above this site, indicating a shift of one “click” downward (Fig. 2 E, panel i; Video S1). Interestingly, hNa_X_ VSDI samples larger GC1-S4 – HC-S2 distances, reaching up to 18.3 Å, and reflecting further activated states of VSDI S4. Notably, Na_X_ VSDI contains only three positively charged residues in S4 compared to four positively charged residues in Na_V_ VSDIV (Noland et al., 2022).

For VSDII GC1-S4 – HC-S2 distances, we observed activated and partially deactivated states, differing by one “click” of gating charge movement, with three versus two gating charges in the S4 segment positioned above the hydrophobic constriction site in the S2 segment (Fig. 2 B). Experimental structures exist for deactivated conformations of VSDII, such as chimeric Na_V_Ab-hNa_V_1.7-VSDII stabilized by the peptide toxin Huwentoxin-IV (PDB: 7K48) (Wisedchaisri et al., 2020), which shows a distance of 9.6 Å and only one gating charge above the hydrophobic constriction site. However, none of our human Na_V_ models sampled these fully deactivated conformations of the VSDII, with the lowest distance observed in a model of hNa_V_1.1 at 13.4 Å. hNa_X_ VSDII stands out, sampling lower distances up to 12.3 Å. Notably, hNa_X_ VSDII contains the same number of positively charged residues in S4 (five) as in Na_V_ VSDII S4 (Noland et al., 2022).

We observed larger GC1-S4 – HC-S2 distances for VSDIII, as this VSD contains an additional gating charge in the S4 segment (Fig. 2 C). Our models sampled the observed conformational space from experimental structures, with hNa_V_1.1 (PDB: 7DTD) (Pan et al., 2021) marking the upper limit at 21.8 Å and hNa_V_1.5 (PDB: 7DTC) (Li et al., 2021b) marking the lower limit at 20.9 Å. This highlights the limited observable states for Na_V_ VSDIII in experimental structures, where in all of them, four gating charges are positioned above the hydrophobic constriction site. In contrast, our models showed a broader range of states. The most activated model generated was for hNa_V_1.8 with a distance of 21.5 Å and four gating charges positioned above the hydrophobic constriction site. The most deactivated VSDIII was found in a model of hNa_V_1.7 with a distance of 15.8 Å, below the limit observed in experimental structures, with three gating charges positioned above the constriction site, corresponding to a shift of one “click” downward (Fig. 2 E, panel iii; Video S1). This state has not yet been observed in any experimental structure of hNa_V_ VSDIII. Additionally, hNa_X_ VSDIII stood out with significantly lower GC1-S4 – HC-S2 distances, consistent with its experimental structure, which also displayed a lower distance of ∼18 Å and reflected further deactivated states of S4. Notably, hNa_X_ VSDIII contains only four positively charged residues in S4 compared to five positively charged residues in Na_V_ VSDIII (Noland et al., 2022).

VSDIV exhibited the broadest range of sampled GC1-S4 – HC-S2 distances (Fig. 2 D). We observed two “clicks” of difference in the translocation of gating charges across the hydrophobic constriction site with different frequencies. The largest and smallest distances were observed in hNa_V_1.8 models, where the most activated state had four gating charges above the constriction site and the most deactivated state had two, resulting in a distance difference of 9.5 Å (Fig. 2 E, panel iv; Video S1). These results encompass the range of states observed in experimental structures, where the upper limit is marked by hNa_V_1.1 (PDB: 7DTD) (Pan et al., 2021) with 22.4 Å, and the lower limit by the Na_V_PaS-hNa_V_1.7 chimera, which has VSDIV trapped in a deactivated state by an α-scorpion toxin (PDB: 6NT4) (Clairfeuille et al., 2019) with a distance of 11.2 Å. The deactivated VSDIV in this chimera shows two gating charges above the hydrophobic constriction site, similar to what we observed in the hNa_V_1.8 model. hNa_X_ VSDIV models also sampled the GC1-S4 – HC-S2 states resulting from the two ’click’ translocations. However, for hNa_X_, the most frequently sampled state was the more deactivated state, in contrast to Na_V_ channels, where the most frequently sampled state was the activated one. Notably, hNa_X_ VSDIV contains only three positively charged residues in S4 compared to six positively charged residues in Na_V_ VSDIV (Noland et al., 2022).

Overall, these results demonstrate AlphaFold2 subsampling method’s ability to partially explore the conformational diversity of Na_V_ channel VSDs. By analyzing the distribution of GC1-S4 – HC-S2 distance coordinates across VSDs and comparing them to experimental structures, we have identified different potential conformational states in the VSDs of all hNa_V_ channels. Notably, AlphaFold2 is able to generate models that sample intermediate states between fully activated and deactivated conformations. These intermediate states may represent important steps in the activation and deactivation pathways. However, our results also reveal certain limitations. For VSDI, VSDII, and VSDIII, we observed only one “click” of activation across the sampled models, potentially missing more deactivated states, particularly for VSDII, where more deactivated conformations have been observed in experimental structures in the presence of natural peptide toxins (Xu et al., 2019; Wisedchaisri et al., 2020).

### AlphaFold modeling reveals frequent sampling of IFM motif in bound and unbound states, along with potential slow inactivation conformations of the selectivity filter

When analyzing the distribution of IFM states using the IFM – PD distance, we observe a distinct bimodal pattern: one state where the IFM is bound to the pore domain, corresponding to a fast-inactivated channel with an IFM – PD distance of ∼8 Å, and another state where the C-terminal region traps the IFM, unbound from the pore domain, with the IFM positioned nearly 30 Å away from its binding site in the pore. Interestingly, all nine hNa_V_ channels sampled both states (Fig. 3 A), although with varying frequencies. Notably, hNa_V_1.9 samples the unbound state more frequently (19.4% of models with IFM – PD Distance < 15 Å) (Fig. S1), in agreement with the slow kinetics of Na_V_1.9 channel activation and inactivation (Dib-Hajj et al., 1999). Multiple other Na_V_ channel subtypes have unequal distribution of the IFM bound and unbound states (Fig. S1). Once again, hNa_X_ stands as an outlier, with only the bound state being sampled. When we analyzed the global distribution of IFM states across the nine hNa_V_ channels (Fig. 3 B), we found that, despite the IFM motif predominantly being observed in the bound state in experimental structures (Table S3), both the bound and unbound states were frequently sampled in our models. A notable result is the appearance of intermediate IFM states between the bound and unbound conformations (Fig. 3 A), states that have not been captured in experimental structures.

**Figure 3.**
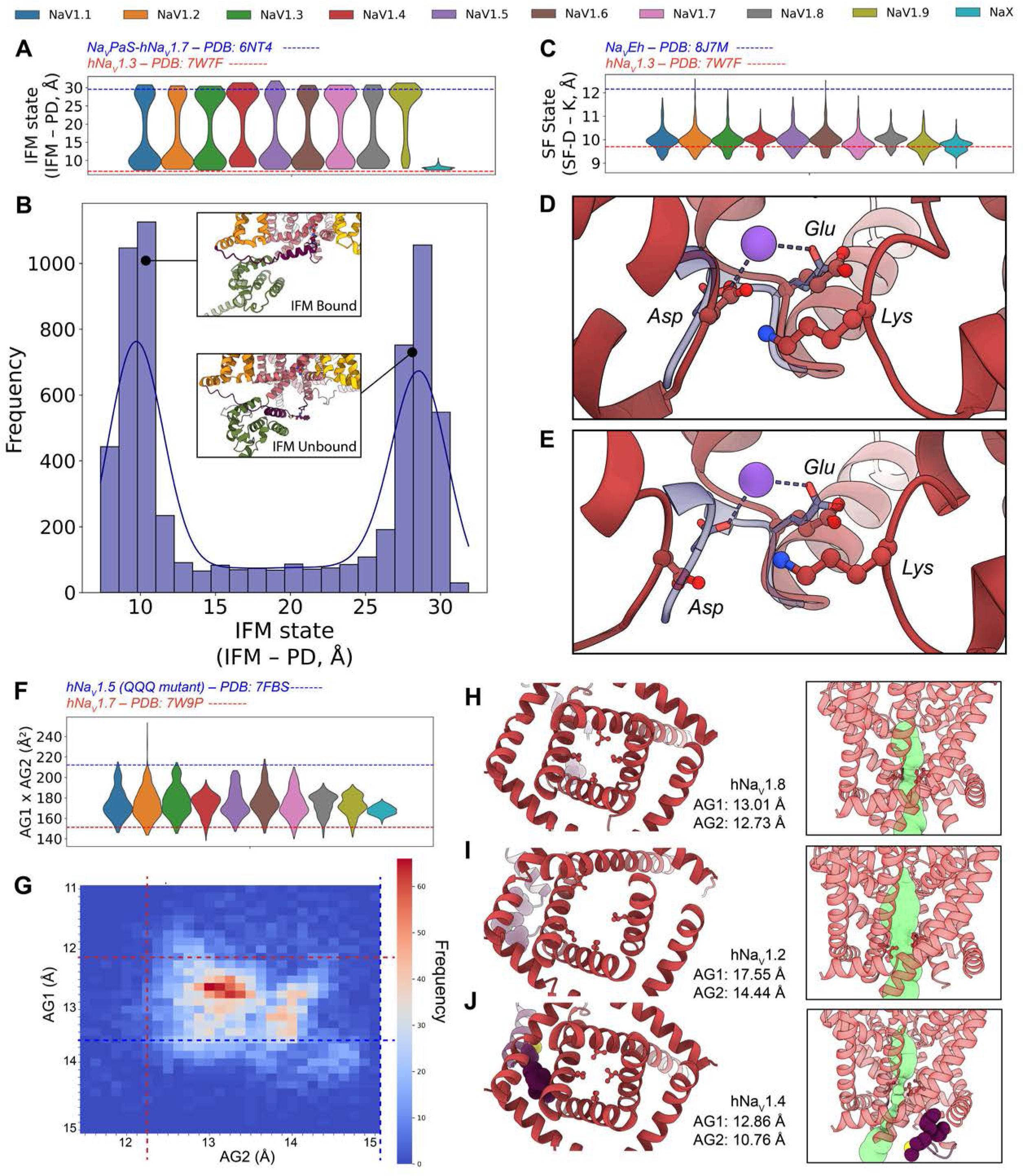
Distribution of IFM, SF and AG states in generated AlphaFold models of hNa_V_ channels. (A) Distribution of the IFM – PD distance for the nine hNa_V_ channels plus hNa_X_. (B) Overall distribution of IFM-PD distances in all generated models. (C) Distribution of the SF-D – K distance for the nine hNa_V_ channels plus hNa_X_. (D, E) Models representing the generated state with lowest (D, 9 Å) and largest (E, 12.6 Å) SF-D – K distances. The side chains of the aspartate, glutamate and lysine of the DEKA motif are shown. The superimposed SF of the experimental structure of hNa_V_1.4 (PDB: 6AGF) (Pan et al., 2018) is shown in purple to illustrate the structural arrangement of the SF that allows Na^+^ coordination. For A and C, the blue dashed lines mark the largest distance observed in experimental structures for that coordinate, while the red dashed lines represent the lowest. (F) Distribution of the AG1xAG2 area for the nine hNa_V_ channels plus hNa_X_. (G) Frequency heatmap showing the combined frequency of the AG1 and AG2 coordinate distances in all hNa_V_ models; the blue dashed lines mark the largest distance observed in experimental structures and the red dashed lines represent the lowest. (H-J) Models representing the most frequent (H), largest (I) and lowest (J) AG area sampled in our models; the IFM motif is highlighted in purple, and the calculated volume of channel pore is represented with the green surface.

In the SF region, we focused on identifying potential SF conformations that might represent slow-inactivated states. Most experimental structures show the SF-D – K distance ranging from 9.7 to 11.2 Å, corresponding to conductive states of the SF (Table S3). However, a recent experimental structure of Na_V_Eh with a putative slow-inactivated state (PDB: 8J7M) showed a significantly larger distance of 12.2 Å, accompanied by clear structural changes that suggest a non-conductive, slow inactivated conformation (Chen et al., 2024). Interestingly, the state distribution across the nine hNa_V_ channels indicates that, while the state ∼10 Å is the most frequently sampled (Fig. 3 C), some channels show low-frequency sampled models with SF-D – K distances similar to the putative slow-inactivated state ∼12 Å. Visually comparing models with the smallest and largest SF-D – K distances, we observe that the model with the lowest distance (9 Å), corresponding to hNa_V_1.9, shows the aspartate and glutamate residues of the DEKA motif positioned to allow sodium coordination (Fig. 3 D). More intriguingly, in the model with the largest distance (12.6 Å) corresponding to hNa_V_1.2, the aspartate is further from the glutamate, disrupting the spatial arrangement necessary for sodium coordination, suggesting that this model may represent a hypothetical slow-inactivated, non-conductive state (Fig. 3 E; Video S2). This is particularly significant, as such a state has not been observed in any experimental structure of a SF containing the DEKA motif. The previously mentioned experimental structure of a putative slow-inactivated Na_V_ channel corresponds to an ancient eukaryotic symmetric channel, Na_V_Eh, that contains the EEEE motif in the SF. Notably, the frequency of sampling of this potential slow-inactivated state varied across the hNa_V_ family models. Channels hNa_V_1.2 and hNa_V_1.6 had the highest number of models in this state, with 8 and 5 models respectively showing SF-D – K distances greater than 11.5 Å. In contrast, hNa_V_1.4, hNa_V_1.8, and hNa_V_1.9 did not produce models with SF-D – K distances corresponding to this state, with the maximum sampled distance being around 11.3 Å (Fig. 3 C). Future studies using more advanced methods, such as AlphaFold3 (Abramson et al., 2024), that support modeling with sodium ions explicitly, will be necessary for a more accurate assessment of SF conformational dynamics under physiological conditions.

### AlphaFold predicts known and novel activation gate states

To investigate the conformational states of the AG across the hNa_V_ channels, we focused on the area formed by multiplying the two calculated distances, AG1 and AG2, which provide insight into the backbone-determined state of the gate (Fig. 1). While we acknowledge that the actual space available for ion permeation depends on the positioning of side chains within the gate, our focus here is on the overall structural state dictated by the backbone. When we calculated the equivalent area in available experimental structures of human channels, we observed that the lower limit was 151 Å², found in hNa_V_1.7 in complex with the peptide toxin Huwentoxin-IV (PDB: 7W9P) (Huang et al., 2022a), a structure hypothesized to represent a closed channel state. The experimental structure of the cockroach Na_V_PaS (PDB: 5X0M) (Shen et al., 2017) exhibits an even more tightly closed pore, with an area of 129.6 Å². On the other hand, experimental structures hypothesized to represent open states showed areas ∼200 Å², such as the rat Na_V_1.5/QQQ mutant (PDB: 7FBS) (Jiang et al., 2021). Notably, hNa_V_ channel experimental structures in fast-inactivated state, in which the IFM motif is bound to its receptor site (such as hNa_V_1.1, PDB: 7DTD, or hNa_V_1.6, PDB: 8FHD), also display AG areas near ∼200 Å². This indicates that while VSD motions can drive backbone opening of the gate, ion conduction is ultimately controlled by small side-chain rearrangements within the AG.

Our models of the nine hNa_V_ channels revealed a distribution of activation gate areas that encompasses this experimental range with different frequences (Fig. 3 F). The smallest area was observed in a model of hNa_V_1.4 at 138.3 Å², while the largest, an outlier, was a model of hNa_V_1.2 with an area of 253 Å² (Fig. 3 F). This outlier model of hNa_V_1.2 was notable, as no other hNa_V_ channel sampled such a large area. Most channels, however, sampled states with areas around 200 Å²except for hNa_V_1.4, hNa_V_1.8, and hNa_V_1.9, which showed smaller areas. hNa_X_ exhibited a narrower distribution compared to the other channels (Fig. 3 F).

When we analyzed the AG1 and AG2 distances together (Fig. 3 G), the most frequently sampled state had AG1 distances between 12.6–12.8 Å and AG2 distances between 13–13.1 Å, resulting in an area range of 163.8–167.3 Å² which represents either a closed or inactivated state. To gain more detailed insights, we visually analyzed models presenting the smallest and largest activation gate areas, as well as a model corresponding to the most frequently sampled state (Video S3). In the model with the most frequently sampled area (Fig. 3 H), we observed a narrow activation gate and an unbound IFM motif, which we identified as a possible closed state. The model with the largest area, the outlier hNa_V_1.2 mentioned earlier (Fig. 3 I), showed a dramatic displacement of the DI-S6 segment, creating a clear open pathway. This model also exhibited the IFM motif in an intermediate state between bound and unbound, suggesting that it may represent a hypothetical low-frequency or short-lived open state. Finally, the model with the smallest area, found in hNa_V_1.4 (Fig. 3 J), displayed a fully bound IFM motif and a closed pathway, which we identified as an inactivated state.

Although this may be an oversimplification, given the likely existence of multiple distinct open, closed, and inactivated states, the key finding is that AlphaFold2 successfully sampled a wide range of activation gate conformations across all hNa_V_ channels, from putative closed and inactivated to fully open states.

### AlphaFold models reveal region-specific relationships between conformational states and model confidence

After demonstrating that AlphaFold2 can model multiple states of Na_V_ channels, we asked two key questions: (1) Do specific states have distinct pLDDT values, indicating varying model confidence? (2) How does the number of recycles affect the distribution of these states? This section addresses these questions by analyzing model confidence, represented by pLDDT values, and state distributions across recycles. For the pLDDT values, we used a subsetting pLDDT calculation, reporting the average pLDDT of the residues forming the region of the channel of interest. All models in this study were generated using six recycles and saving intermediate recycle models.

We first examine the correlation between the distinct conformational states of key regions in the Na_V_ channels and their respective pLDDT values. For the VSDs, we observe distinct distributions for each of the four domains. In VSDI, both deactivated and activated states achieve high confidence; however, a trend emerges where more activated states tend to receive higher pLDDT scores (Fig. S2 A). In VSDII, the most confident models are generally those in intermediate states with pLDDT values up to 86, while the more activated conformations have pLDDT values ∼82 (Fig. S2 B). A similar pattern is seen in VSDIII (Fig. S2 C), whereas in VSDIV (Fig. S2 D), we note that more deactivated states have slightly lower confidence scores (pLDDT ∼75), while fully activated states exceed a pLDDT of 80.

For the IFM Motif (Fig. S2 E), a clear pattern emerges: the bound state consistently achieves the highest pLDDT values (approaching 70), while the unbound state—where the IFM motif is trapped by the C-T domain—shows moderate values (∼55). Intermediate states between the bound and unbound conformations exhibit the lowest pLDDT values (∼30-40). Analyzing the AG (Fig. S2 F), the highest pLDDT scores (∼80) are associated with areas between 150-190 Å^2^. Possible open states with activation gate areas around ∼190-220 Å^2^ cluster around pLDDT values of 65-70, whereas outlier models with activation gate areas larger than 220 receive lower pLDDT values, ∼60. The selectivity filter (Fig. S2 G) also exhibits a trend: less dilated conformations have higher pLDDT scores (reaching 90), whereas states showing greater dilation have reduced model confidence (∼70-75).

These observations reveal that the relationship between conformational states and pLDDT generates distinct distributions for each Na_V_ region, providing insight into how model confidence varies across states. When further analyzing the relationship between states and the number of recycles, a pattern emerges in which outlier states, such as extreme conformations, predominantly appear at Recycle 0, with models clustering more tightly as the number of Recycles increases (Fig. S2 A-G). This trend leads us to investigate the evolution of state distributions as a function of the number of recycles.

We first examined how recycling affects the overall confidence of the models. Fig. S2H shows that the adjusted-global pLDDT score for the full Na_V_ channel increases from recycle 0 to recycle 1 on average, after which it stabilizes. Then, in examining the distribution of region-specific states over an increasing number of recycles, we observed that intermediate IFM states are primarily sampled at recycle 0, while increasing recycles tend to favor either the bound or unbound IFM state (Fig. S2 I). A similar trend is observed for the SF (Fig. S2 J), where more dilated conformations are sampled at recycle 0. For the AG (Fig. S2 K), we observe that super-open states are sampled exclusively at recycle 0, while recycles 1 through 6 converge on states consistent with experimental structures (between 150 and 200 Å^2^ in gate area). The VSDs exhibit a notable trend: deactivated states are more commonly sampled at recycle 0, shifting toward more activated states in subsequent recycles, particularly in VSDIV (Fig. S2 L). Across recycles, VSD pLDDT values rise slightly between recycle 0 and 1 before reaching a plateau (Fig. S2 M).

Our results reveal a relationship between pLDDT and the conformational states that depends on the specific region of the channel, with distinct patterns emerging. While some models with outlier states may have slightly lower pLDDT scores, the lowest values remain around a pLDDT of 60-70, which suggests moderate confidence, meaning the reliability of these models cannot be completely dismissed. The number of recycles significantly impacts the generated conformational distribution, with recycle 0 showing the greatest diversity of states, while increased recycling tends to bias the modeling towards particular states in each region.

Finally, we compared our models to available experimental structures and examined how structural agreement relates to model confidence. For each hNa_V_ subtype with a reference structure (all except hNa_V_1.9), we computed backbone RMSDs between every generated model and a corresponding subtype experimental structure (Fig. S3, i). Across channels, top models typically lie within ∼1.0–1.3 Å RMSD of the reference, and, except for hNa_V_1.5 and hNa_V_1.8, the lowest-RMSD models also exhibit the highest global pLDDT scores (∼78–80). For hNa_V_1.5 and hNa_V_1.8, the best-aligning models fall in an intermediate global pLDDT range (∼75–76), though the RMSD advantage over higher-pLDDT models is minimal. Notably, the largest RMSDs are consistently observed at recycle 0 (Fig. S3, ii), reinforcing that recycle 0 samples the broadest, and often most divergent, portion of conformational space, whereas additional recycles drive convergence toward experimentally observed structures.

### Minimal impact of state-specific custom templates on VSD state distributions

We then tested whether using state-specific custom templates could bias AlphaFold towards states represented by those in the experimental structure templates. As a practical case, we focused on the hNa_V_1.7 channel, which has several structures of the VSDII in the deactivated conformation. To conduct this analysis, we used two specific templates: PDB 6N4R (Xu et al., 2019) and PDB 7K48 (Wisedchaisri et al., 2020). These structures are chimeric constructs in which the human Na_V_1.7 VSDII sequence is grafted onto a bacterial channel Na_V_Ab. Both structures are four-fold symmetric, with all four VSDs in the deactivated conformation, (stabilized by peptide toxins) providing four copies of the deactivated VSD per structure.

We generated the same number of models under identical conditions for hNa_V_1.7, but instead of using the default template mode (where AlphaFold automatically selects relevant templates from a predefined refined version of the PDB to guide predictions), we used a custom template mode (See Methods), providing only the two aforementioned structures as templates. The results indicate that the overall effect of using these templates is minimal (Fig. S2 N-P). When observing the distribution of states across the four VSDs, we found no significant differences between models generated with the default template mode and those generated with the custom template mode, particularly in VSDII, where an effect would be most expected (Fig. S2 N).We then examined the VSD state distribution from recycle 0 to 6 in the default and custom template model sets. In VSDII, we observed that at recycle 0 and 1, more deactivated states were sampled with the custom templates compared to the default mode (Fig. S2 O). However, as the number of recycles increased, the differences between default and custom template modes diminished. Interestingly, a more pronounced effect was observed in VSDIV (Fig. S2 P), as using the custom template mode resulted in deactivated states being sampled in recycles 4, 5, and 6, while the default mode only sampled deactivated states up to recycle 3..

### Correlation analysis uncovers coherent structural coupling across Na_V_ functional regions

We next asked whether state coordinates from different regions correlate in a way that reflects known Na_V_ gating mechanisms. For each human subtype (Na_V_1.1– Na_V_1.9) and Na_X_, we quantified dependence between every pair of coordinates in all generated models using two complementary metrics: Spearman’s ρ (monotonic, rank-based) (Fig. 4 A, i), and Mutual Information (MI; non-parametric sensitivity to non-monotonic structure) (Fig. 4 A, ii).

**Figure 4.**
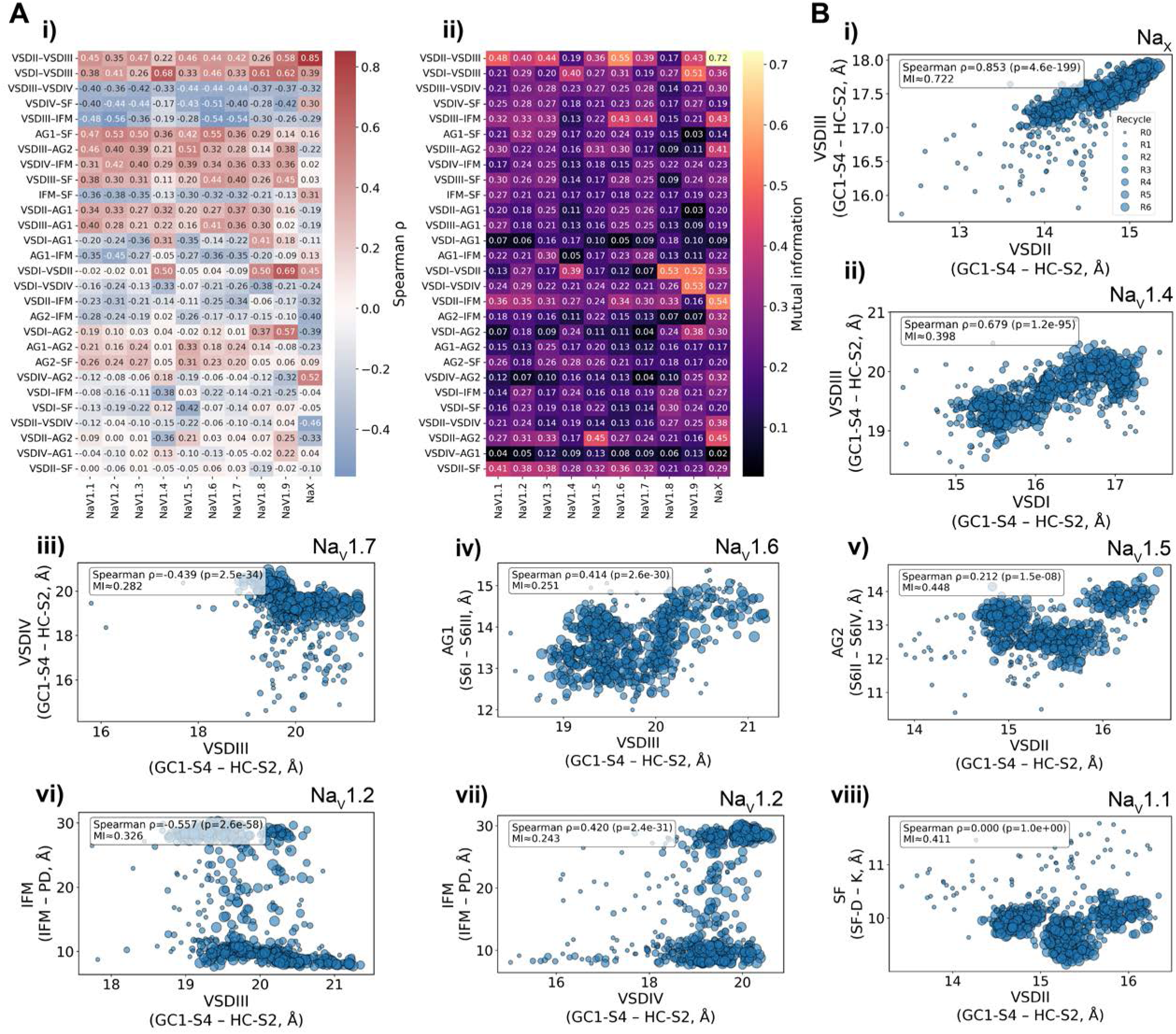
Correlations between state coordinates of different hNa_V_ channel regions. (A) (A) Pairwise dependence heatmaps summarizing correlations between all state coordinates across nine hNa_V_ subtypes. (i) Spearman’s ρ (monotonic dependence) and (ii) Mutual Information (MI; non-parametric sensitivity to non-monotonic dependence). (B) Representative subtype-specific examples of state-state relationships illustrated as scatter plots, showing monotonic (ρ) and non-monotonic (MI) dependencies; dot sizes represent recycle number.

Across VSD–VSD pairs, we observed robust, subtype-dependent correlations. The strongest and most consistent coupling was between VSDII and VSDIII: Spearman’s ρ ranged from 0.22 (hNa_V_1.4) to 0.58 (hNa_V_1.9), with MI between 0.20 and 0.55. hNa_X_ stood out with markedly higher coordination (VSDII–VSDIII ρ=0.85; MI=0.72) (Fig. 4 B, i). VSDI also correlated VSDIII positively in all subtypes (ρ up to ∼0.68 in hNa_V_1.4, hNa_V_1.8, and hNa_V_1.9; large MI up to ∼0.4-0.5) (Fig. 4 B, ii). By contrast, VSDI–VSDII was generally weak, except in those same three subtypes (hNa_V_1.4, hNa_V_1.8, and hNa_V_1.9), which showed a clear positive association (ρ≈0.5-0.6) and higher MI (∼0.4-0.5). VSDIV behaved differently from the other sensors: VSDI and VSDIII both tended to negatively correlate with VSDIV activation (VSDI–VSDIV ρ≈−0.1 to −0.3; VSDIII–VSDIV ρ≈−0.3 to −0.4, Fig. 4 B, iii), whereas VSDII showed only weak dependences (ρ≈−0.1 and MI up to ∼0.2). These patterns align with VSDIV’s distinct mechanistic role in Na_V_ channel function.

Coupling between VSD activation and AG coordinates further reflected the expected known molecular mechanisms. AG1 (S6_I_–S6_III_) increased with VSDII and VSDIII activation (typical ρ≈0.3), consistent with these sensors driving pore opening (Fig. 4 B, iv); hNa_V_1.9 showed a weaker effect, and hNa_X_ inverted the trend (negative ρ). VSDI generally negatively correlated with AG1 (ρ≈−0.2 to −0.3), again with hNa_V_1.4/1.8/1.9 as exceptions showing positive coupling (ρ≈0.2 to 0.4). For AG2 (S6_II_–S6_IV_), VSDI was largely uncorrelated except in hNa_V_1.8 (ρ=0.37; MI=0.24) and hNa_V_1.9 (ρ=0.57; MI=0.38). VSDII Spearman dependences with AG2 were weak overall, except for hNa_V_1.4 (negative, ρ=−0.36), hNa_V_1.5 (positive, ρ=0.21) (Fig. 4 B, v) and hNa_V_1.9 (positive, ρ=0.25), although large MI values for all subtypes (∼0.2-0.4) suggest a more complex, non-monotonic dependence. VSDIII correlated positively with AG2 (ρ≈0.3–0.5) in most subtypes, attenuated in hNa_V_1.4 and hNa_V_1.8 (ρ≈0.1–0.2). VSDIV showed little correlation to AG in all subtypes, but hNa_X_ again deviated with a strong positive VSDIV–AG2 correlation (ρ=0.52). As expected, AG1 and AG2 were positively coupled (ρ≈0.2– 0.3) in most channels, with weaker dependences in hNa_V_1.4 and hNa_V_1.9.

Relations with the IFM motif were consistent for VSDI to VSDIII and unexpectedly mixed for VSDIV. VSDI–III activation is associated with a more “bound” IFM (lower IFM–PD distance), yielding negative ρ and elevated MI, meaning that the most activated VSDI–III models coincide with IFM bound state, with VSDIII displaying the strongest correlations (ρ≈0.3-0.6) (Fig. 4 B, vi). In contrast, VSDIV showed a positive ρ with IFM–PD and, importantly, the scatter plots (Fig. 4 B, vii) reveal that highly activated VSDIV states co-occur with both bound and unbound IFM configurations. This observation is compatible with more nuanced recent models of fast inactivation in which VSDIV activation is necessary but not always sufficient for IFM engagement, allowing VSDIV-activated yet IFM-unbound states to exist (Liu et al., 2023).

Finally, dependencies between VSD activation and the SF were generally diffuse and heterogeneous across subtypes. VSDI showed low correlations with SF dilation, except in hNa_V_1.5, where a moderate negative association was detected (ρ=−0.42). VSDII–SF relationships were also weak in terms of monotonic trends (ρ ≈ 0), but elevated MI values (up to 0.41) indicated non-monotonic, likely U-shaped, dependencies (Fig. 4 B, viii). VSDIII displayed a clearer positive association with SF dilation (ρ up to 0.45 in hNa_V_1.9). In contrast, VSDIV showed the opposite pattern, with consistently negative correlations (as low as ρ =−0.56 in hNa_V_1.2) and MI values up to ∼0.4.

Together, these results indicate coherent, statistically supported correlations among Na_V_ state coordinates. The clearest signals are (i) coordinated activation among VSDs, especially VSDII with VSDIII, and (ii) positive VSDII/VSDIII–AG coupling consistent with pore opening, contrasted by VSDIV’s distinct, often negatively correlated behavior. Notably, hNa_X_ consistently separates from the Na_V_ pattern, as expected from its non-voltage-gated nature. Additionally, we observed that correlation magnitudes generally increase with recycle number, with the strongest associations (particularly for VSD-VSD pairs) emerging at higher recycles, and weak correlations when only considering models at recycle 0 (Fig. S4).

### Clustering analysis of modeled conformational ensembles

Given the heterogeneous dependencies among state coordinates, we next asked whether the models cluster into recurrent patterns of hNa_V_ channel state distributions. We built a joint feature matrix (distance-based coordinates plus region-specific and global pLDDT) across all nine hNa_V_ subtypes generated models, excluding recycle-0 models (which consistently sampled broader and less converged conformations in our previous analyses), and performed Leiden community detection (Traag et al., 2019) on a 500-NN graph (see Methods). The UMAP projection shows five well-separated clusters (0–4) (Fig. 5 A), which are relatively balanced in size, comprising approximately 20%, 26%, 20%, 20%, and 13% of all models, respectively. Distributions of state coordinates and pLDDT by cluster are summarized in Fig. 5B (i–vii) and Fig. S5, respectively.

**Figure 5.**
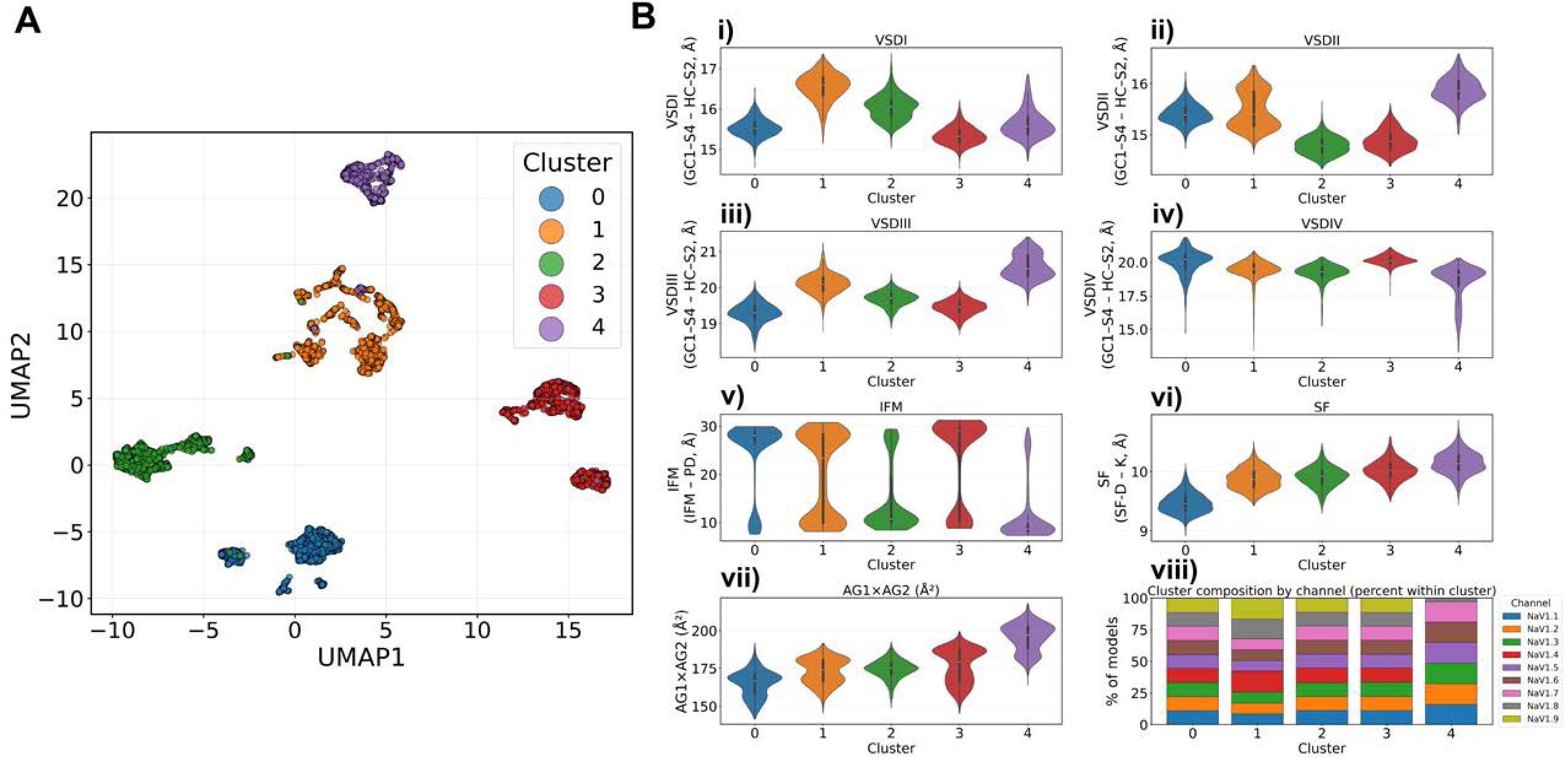
Clustering analysis of modeled conformational ensembles. (A) UMAP projection of all hNa_V_ models (recycles 1–6) built from distance-based state coordinates and pLDDT features. Five clusters (0–4) were identified using Leiden community detection on a 500-nearest-neighbor graph. (B) Violin plots showing the distribution of state coordinates across clusters: (i–iv) state distributions corresponding to the four VSDs (GC1-S4 – HC-S2 distances), (v) IFM (IFM – PD distance), (vi) SF (SF D–K distance), and (vii) AG area (AG1 x AG2). (viii) Cluster composition by hNa_V_ subtype, shown as the percentage of models within each cluster.

Cluster-level patterns suggest interpretable putative channel states. Cluster 0 shows the smallest AG areas (Fig. 5 B, vii) with IFM predominantly unbound (Fig. 5 B, v) and VSDI/VSDIII shifted down (Fig. 5 C, i & iii), consistent with a closed state, although VSDIV appears predominantly in the “up” state (Fig. 5 B, iv) and VSDII in an intermediate state (Fig. 5 B, ii), complicating definitive state assignment. Cluster 1 shows intermediate AG areas (∼160-180 Å^2^), a mixed IFM distribution, and broad VSD activation, resembling a basal/intermediate ensemble. In Cluster 2, the IFM is mostly bound, the AG areas are intermediate (∼175 Å^2^), and the VSDI/VSDIII are more activated, consistent with a fast-inactivated-like state. Cluster 3 exhibits larger AG areas, reaching values close to 200 Å², with the IFM still unbound, consistent with a more activated/open-like ensemble. Cluster 4 combines the largest AG areas, a dilated SF, IFM bound, and more deactivated VSDIV (Fig. 5B, iv–vii), which might represent a different type of inactivated state.

Confidence trends were consistent with these cluster-level distinctions (Fig. S5). Cluster 0 showed the highest global and region-specific pLDDT values followed by Cluster 2, indicating that these ensembles correspond to more confident structural predictions, whereas Clusters 1, 3, and 4 displayed lower pLDDT values, with Cluster 4 exhibiting the lowest confidence values overall (Fig. S5 A). Interestingly, pLDDT values for the IFM motif were relatively uniform across clusters (Fig. S5 G), suggesting comparable local confidence for this region regardless of state. In contrast, Clusters 1, 3 and 4 showed noticeably lower pLDDT in the other structural regions.

Subtype composition provided additional insight into the organization of these clusters (Fig. 5 B, viii). Clusters 0, 2, and 3 were evenly populated by all nine hNa_V_ subtypes, despite the clustering algorithm being agnostic to subtype identity, supporting that these three ensembles reflect broadly sampled conformational regimes. By contrast, Cluster 1 was enriched in hNa_V_1.8 and hNa_V_1.9 models, while Cluster 4 lacked hNa_V_1.9 entirely and contained fewer hNa_V_1.8 models (Fig. 5 B, viii). This channel-dependent occupancy of Clusters 1 and 4 for hNa_V_1.8 and hNa_V_1.9 is notable, as NaV1.8 and NaV1.9 are known to exhibit distinct inactivation behavior compared to other Na_V_ subtypes (Dib-Hajj et al., 1999).

We stress that the state labels used here should be regarded as preliminary, hypothesis-driven interpretations rather than definitive functional assignments. They are motivated by coherent patterns emerging from the clustering analysis and by established mechanistic principles of Na_V_ gating, but they do not by themselves constitute proof of state identity. A more formal state classification would require broader conformational sampling (particularly of the VSDs), deriving specific structural hypotheses from these ensembles and testing them experimentally or evaluating their energetic plausibility using physics-based simulations. Nevertheless, this analysis demonstrates that subsampled AlphaFold2 ensembles do not distribute randomly across conformational space but instead resolve into a small number of recurrent, interpretable patterns of state distributions, suggesting that this workflow captures functionally meaningful structural variation.

### High accuracy of α and β-subunit complex models generated by AlphaFold Multimer agree with experimental structures

Up until now, we have focused on modeling the Na_V_ α-subunit alone, which forms the core of the channel itself. However, in physiological conditions these channels are regulated by auxiliary protein partners, the most significant of which are the auxiliary β-subunits: β1, β2, β3, and β4 (Namadurai et al., 2015). These β-subunits have two domains: a transmembrane domain and an extracellular immunoglobulin domain. We modeled hNa_V_1.7 paired with each of the four β-subunits, generating 100 models with 6 recycles per case. Additionally, we used hNa_V_1.1 as a control to validate the replicability of our results. For each of these cases, we wanted to assess whether the generated models align with the experimental structures available for the different complexes.

We compared the top-ranked model of hNa_V_1.7 paired with each of the β-subunits to reference experimental structures (Fig. 6 A-D, i). Overall, we observed an almost perfect alignment between the experimental structures and our models. For β1 (Fig. 6 A), the β-subunit is positioned next to VSDIII, consistent with the reference structure (PDB: 7W9K, hNa_V_1.7-β1-β2 complex) (Huang et al., 2022a), with an RMSD value of 0.7 Å when superimposing the auxiliary subunits. For β2 (Fig. 6 B), the immunoglobulin domain aligns perfectly with the experimental structure (PDB: 7W9K, hNa_V_1.7-β1-β2 complex) (Huang et al., 2022a) and is situated above VSDI, with an RMSD value of 0.4 Å. For β3 (Fig. 6 C), both the immunoglobulin domain and transmembrane domain align well with the reference experimental structure (PDB: 7TJ8, hNa_X_-β3 complex) (Noland et al., 2022) and are positioned next to VSDIII, with an RMSD value of 2.2 Å. For β4 (Fig. 6 D), while we observe some shift relative to the reference experimental structure (PDB: 7DTD, hNa_V_1.1-β4 complex) (Pan et al., 2021) when superimposing the full channel, it is also accurately positioned next to VSDI, with an RMSD value of 1.2 Å when superimposing the β4 subunit alone. The highest RMSD value is for β3, which is expected because the comparison is with the reference experimental structure of the Na_X_ channel, not a Na_V_ channel. Therefore, the observed difference in RMSD could reflect conformational changes specific to the β-subunit that depend on the type of α-subunit involved. For both β2 and β4, while the transmembrane domain is known to exist, it is not resolved in the available experimental structures, likely due to conformational flexibility. Despite this, the immunoglobulin domain aligns well, and the transmembrane domain appears slightly tilted relative to the membrane axis in our models. Top models of hNa_V_1.1 in complex with the β-subunits showed similar patterns and also aligned well with the reference experimental structures (Fig. S6).

**Figure 6.**
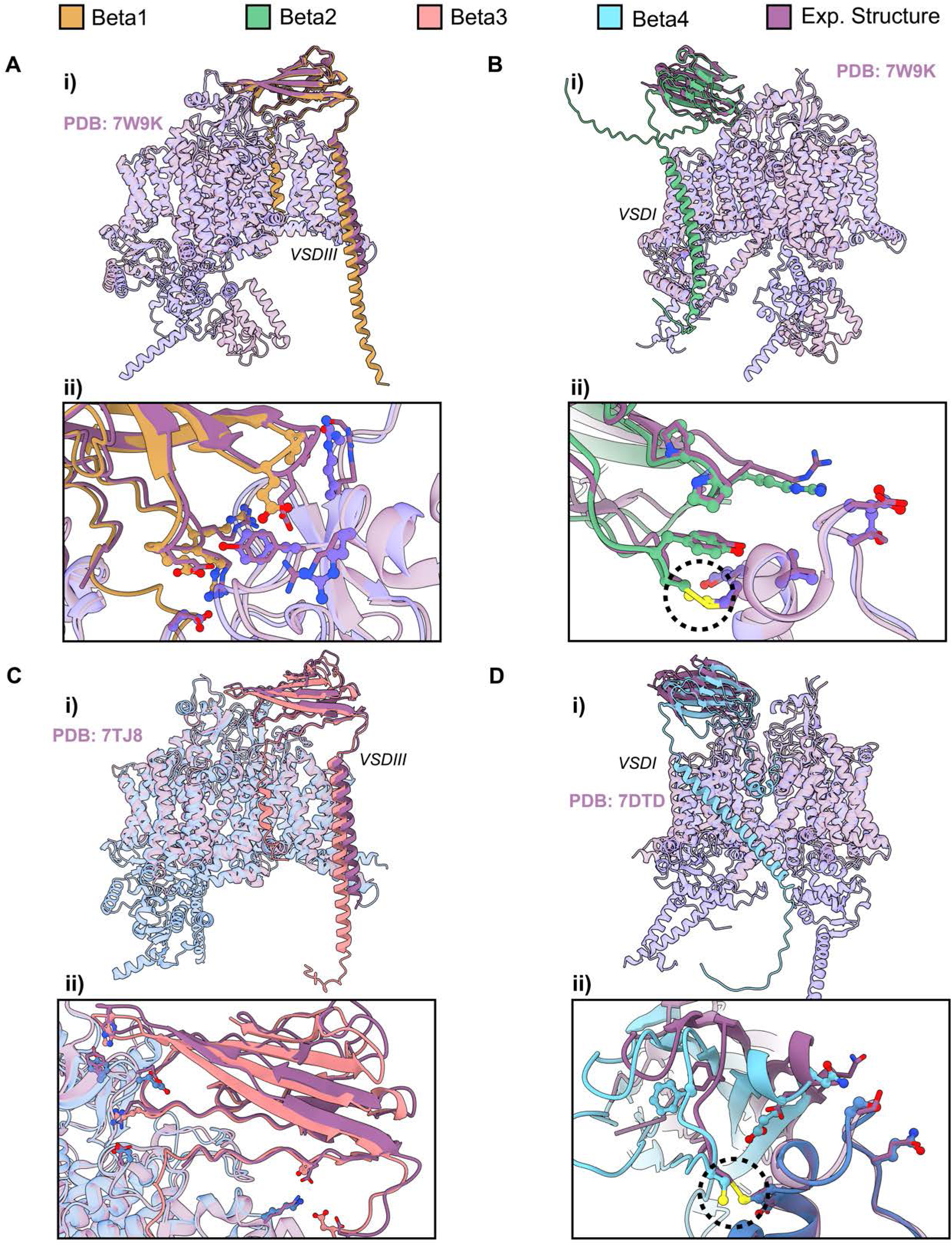
Comparison of top models of hNa_V_1.7 in complex with auxiliary β-subunits with reference experimental structures. (A-D, i) Superimposition of top ranked AlphaFold models of hNa_V_1.7 in complex with the four auxiliary β-subunits with the corresponding experimental structure of reference: (A,B) hNa_V_1.7-β1-β2 complex, PDB: 7W9K (Huang et al., 2022a); (C) hNa_X_-β3 complex, PDB: 7TJ8 (Noland et al., 2022); (D) hNa_V_1.1-β4 complex, PDB: 7DTD (Pan et al., 2021). (A-D,ii) Detailed view of the α-β interface comparing top models (side chains represented as balls and sticks) with experimental structures (side chains shown as sticks).

An important distinction among the auxiliary subunits is that β2 and β4 interact with the Na_V_ α-subunit by forming disulfide bridges, whereas β1 and β3 interact with the Na_V_ α-subunit non-covalently. When examining the atomic details of the interface for all four β-subunits (Fig. 6 A-D, ii), we found nearly perfect alignment between the experimental structures and our models, both in the interactions present and in the conformations of the side chains of amino acids at the interface. Notably, for β2 and β4, the cysteines responsible for forming disulfide bridges are positioned accurately to establish these bonds, with the sulfur atoms of the cysteines nearly perfectly aligned with the experimental structures (Fig. 6 B and D, ii, dashed circles). Although AlphaFold2 does not model covalent bonds between different protein partners, the positioning of the sulfur atoms suggests a high likelihood of disulfide bridge formation.

When analyzing the distribution of pLDDT values across the four β-subunits, we observe that β1 and β3 have a similar distribution (Fig. S7 A and C). The immunoglobulin domain has very high pLDDT values, ∼90, while the transmembrane domain maintains relatively high accuracy but exhibits more variability in pLDDT values. We also observe that the N-terminus of the protein, the signal peptide, is inserted into the membrane with low confidence, but we know this part is likely cleaved and doesn’t form part of the final structure. The end of the transmembrane domain in the intracellular region, which is not resolved in experimental structures, also has low confidence. For β2 and β4, we see a similar pattern between the two (Fig. S7 B and D). The immunoglobulin domain has higher pLDDT values, again ∼80-90. For β4, we observe regions of the immunoglobulin domain with lower pLDDT values, suggesting that β4 immunoglobulin domain may exhibit more conformational flexibility. The difference we observe in the transmembrane domain (keeping in mind that, for β2 and β4, this domain is unresolved in experimental structures) is not only that it is tilted with respect to the membrane axis but also that it shows significantly lower confidence, with pLDDT values dropping below 50.

The other question we addressed was whether the presence of the β-subunits affects the pLDDT distribution of the α-subunit (Fig. S7 E). For most of the structured regions of the α-subunit’s four domains, the presence of the β-subunits increases pLDDT compared to when the β-subunits are absent. This applies to the structured regions. However, in the unstructured intracellular loops, we observe that when the α-subunit is alone, it exhibits a higher pLDDT. Although these values remain relatively low, below 30 in all cases, this indicates high uncertainty in the conformation of this entire region. Therefore, the presence of β-subunits appears to increase the pLDDT of the structured regions of the α-subunit, which may indicate conformational stabilization due to the β-subunits.

### Effect of recycles and ipTM values on Na_V_ α-β multimer complex modeling

The next aspect we wanted to investigate, knowing that high-accuracy models can be generated, was how these multimer models evolve over the course of six recycles, and to analyze a confidence parameter beyond pLDDT—namely, ipTM. ipTM is a metric reported by AlphaFold Multimer that relates to the confidence in the interface between the different monomers that form the multimer complex and takes values from 0 (low confidence) to 1 (high confidence).

To conduct this analysis, we examined the top model for each of the four β-subunits in complex with hNa_V_1.7, observing how the structures evolved from recycle 0 to recycle 6 (Fig. S8 A-D). At recycle 0, the β-subunit is positioned in an improbable location with numerous steric clashes with the α-subunit. As the number of recycles progresses to recycles 1 and 2, the β-subunit gradually relocates to the appropriate region, either adjacent to VSDI or VSDIII, depending on the specific β-subunit. From Recycle 3 onwards, minimal structural changes occur, with no significant alterations observed up to Recycle 6 (Fig. S8 A-D; Video S4).

We next analyzed the ipTM values and their evolution over the number of recycles. Consistent with our visual observations, ipTM values increase significantly from Recycle 0 to 1, 2, and 3, after which they stabilize across all four β-subunits (Fig. S8 E). We also observed distinct differences between the four subunits. β1 and β3 achieve higher ipTM values, reaching up to 0.8 in some instances, whereas β2 and β4 have lower ipTM values ∼0.5. Notably, β4, which exhibited lower confidence, also displayed the lowest ipTM values.

Finally, we investigated the relationship between ipTM values and the pLDDT of the β-subunit alone. As expected, a positive correlation was observed, with models that had higher ipTM values also demonstrating higher pLDDT values for the β-subunit (Fig. S8 F). For β1 and β3, there was a clearer trend towards higher ipTM and pLDDT values, while β2 and β4 showed more models within the intermediate ipTM range.

### Auxiliary β-subunits profoundly influence the conformational landscape of Na_V_ α-subunit, especially for VSDIV

After analyzing the results presented so far, we sought to determine if the presence of the different β-subunits affects the distribution of conformations in the different regions of the α-subunit. We first examined the impact of the β-subunits on the distribution of states of the VSDs (Fig. 7 A). Overall, we found that the effect on VSDII and VSDIII was minimal, with very small differences in GC1-S4 – HC-S2 distance distributions. However, larger effects were observed in VSDI, where the presence of auxiliary subunits, particularly β1 and β3, led to more deactivated states being sampled. The most pronounced effect was observed in VSDIV, where the presence of β-subunits significantly altered the state distribution, with a much greater frequency of deactivated states with lower GC1-S4 – HC-S2 distances (10-12 Å).

**Figure 7.**
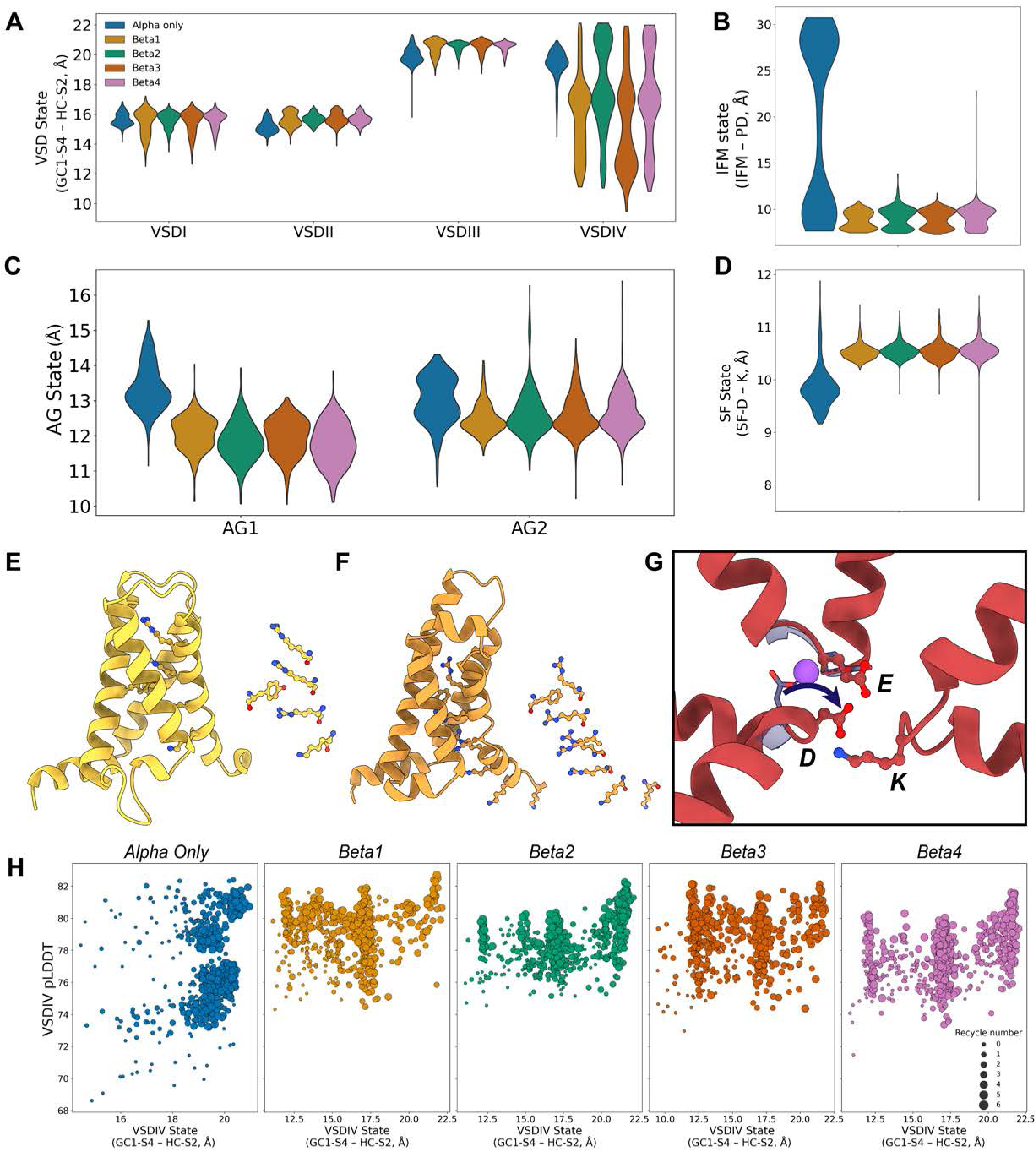
Effect of the presence of auxiliary β-subunits on α-subunit conformational distribution. (A-D) Comparison of the distribution of states of the VSDs (A), IFM motif (B), AG (C), and SF (D) in the presence of the four different β-subunits and when the α-subunit is modeled alone. (E) Model of hNa_V_1.7-β-1 that shows a VSDI in a more deactivated state than what was observed with the α-subunit alone. (F) Model of hNa_V_1.7-β-3 that shows the VSDIV in a state representing one additional “click” of translocation of gating charges across the hydrophobic constriction site from what was observed for the α-subunit alone. (G) Outlier model of hNa_V_1.7-β-4 that shows a contracted SF where the aspartate interacts with the lysine from the DEKA motif. (H) Relationship between VSDIV state distributions for each modeling case (4 β-subunits plus α-subunit alone) and the VSDIV pLDDT; dot sizes represent recycle number.

We then visually inspected models that might represent novel states introduced by the presence of the β-subunits in the modeling. In the model with VSDI in the most deactivated state, which corresponds to a model with β1, we observed a conformation similar to the most deactivated VSDI state seen with the α-subunit alone: only two gating charges were positioned above the hydrophobic constriction site (Fig. 7 E). However, compared to the most deactivated VSDI α-only model (Fig. 2 A), the third gating charge was positioned significantly lower relative to the hydrophobic constriction site in the model with β1. Interestingly, for the model with β3, where VSDIV was in its most deactivated state (GC1-S4 – HC-S2 ≈ 10 Å), we noted that this state was more deactivated than what was observed in the α-only model. In the α-only model, the most deactivated state reached had two gating charges above the hydrophobic constriction site and GC1-S4 – HC-S2 ≈ 13 Å (Fig. 2 D). In contrast, in the β3 model, only one gating charge was positioned above the hydrophobic constriction site, which results in an additional “click” down to what we have previously observed (Fig. 7 F, Video S5). This represents a state that has not been observed experimentally.

We observed a significant effect of the β-subunits on the IFM motif conformations distribution (Fig. 7 B). With the α-subunit alone, a bimodal distribution of the IFM motif, being either bound or unbound, was present (Fig. 3 AB). However, in the presence of the β-subunits, the IFM motif was more frequently modeled bound to the pore domain (Fig. 7 B), suggesting that the presence of β-subunits biases the models towards an IFM-bound state.

Analysis of the AG (Fig. 7 C) revealed that the presence of β-subunits generally reduced the area of the activation gate, although certain outliers with larger areas appeared. For the SF (Fig. 7 D), we observed significant changes in the SF-D–K distance distribution, with the average distances being more dilated (∼10.5 Å). Interestingly, for β4, we identified outlier models with a highly contracted selectivity filter, with distance coordinate values below 8 Å. Visual analysis of this outlier (Fig. 7 G; Video S6) suggests that the contraction is due to an interaction between the aspartic acid and lysine residues of the DEKA motif, disrupting sodium coordination. This interaction may represent another possible slow inactivated state.

Returning to VSDIV, we observed that the presence of β-subunits led to models spanning a broad range of states, from very deactivated to fully activated. This provided an opportunity to examine the relationship between pLDDT values and a broad range of VSDIV states (Fig. 7 H). The results revealed that the presence of β-subunits led to a complete redistribution of pLDDT/state relationships in VSDIV compared to the α-only models. Notably, even the most deactivated states of VSDIV (GC1-S4 – HC-S2 distances ∼10-12 Å), when β-subunits were present, maintained high pLDDT values ∼80, indicating that these deactivated models were generated with high confidence.

Our results show that the presence of β-subunits during AlphaFold modeling profoundly affects the modeled conformational landscape of the α-subunit, resulting in novel states not observed when modeling the α-subunit alone. Notably, we observed a similar reshaping of state distributions for hNa_V_1.1 (Fig. S6 E-H).

### Modeling Na_V_-CaM interactions resulted in sampling of both Ca²⁺-bound and Ca²⁺-free conformations

In the final part of this study, we explored the interaction between Na_V_ channel α-subunit and CaM, a Ca²⁺ sensor protein known to modulate Na_V_ channel activity. Particularly, CaM modulates channel inactivation by interacting with the α-subunit C-T region, specifically with an alpha-helical segment known as the IQ motif (Wu and Hong, 2021). Unlike the auxiliary β-subunits, CaM is a protein with greater conformational diversity that depends on Ca²⁺ concentration, making model analysis more complex. Moreover, there is no experimental structure of a full Na_V_ channel bound to CaM, although partial structures with CaM bound to the IQ segment in the C-T domain of Na_V_ channels have been resolved (Gabelli et al., 2014; Wang et al., 2014; Hovey et al., 2017). This analysis is also particularly interesting because AlphaFold2 does not support modeling of non-protein atoms, like cations. Therefore, Ca²⁺ was absent from our modeling; in the future, with newer versions of AlphaFold that support modeling of cations, more rigorous analysis could be conducted. In this case, we focused on two test cases, hNa_V_1.2 and hNa_V_1.5, and modeled both with CaM using the same setup as before, generating 100 models with six recycles.

The first analysis involved visually analyzing the evolution of the top-ranked models over the six recycles for each case. The behavior of the IFM motif and CaM conformation revealed notable differences between hNa_V_1.2 and hNa_V_1.5 (Fig. 8 A and C). Interestingly, in both channels, the IFM motif initially binds to the pore domain at recycle 0 but becomes unbound as recycles progress. Additionally, when analyzing how ipTM values evolve over the number of recycles, we observe that for hNa_V_1.2, ipTM increases more gradually and doesn’t stabilize until recycle 5 (Fig. 8 B). In contrast, for hNa_V_1.5, ipTM quickly stabilizes by recycle 2, reaching values ∼0.8 (Fig. 8 D). For hNa_V_1.2, the ipTM values are slightly lower, averaging ∼0.7.

**Figure 8.**
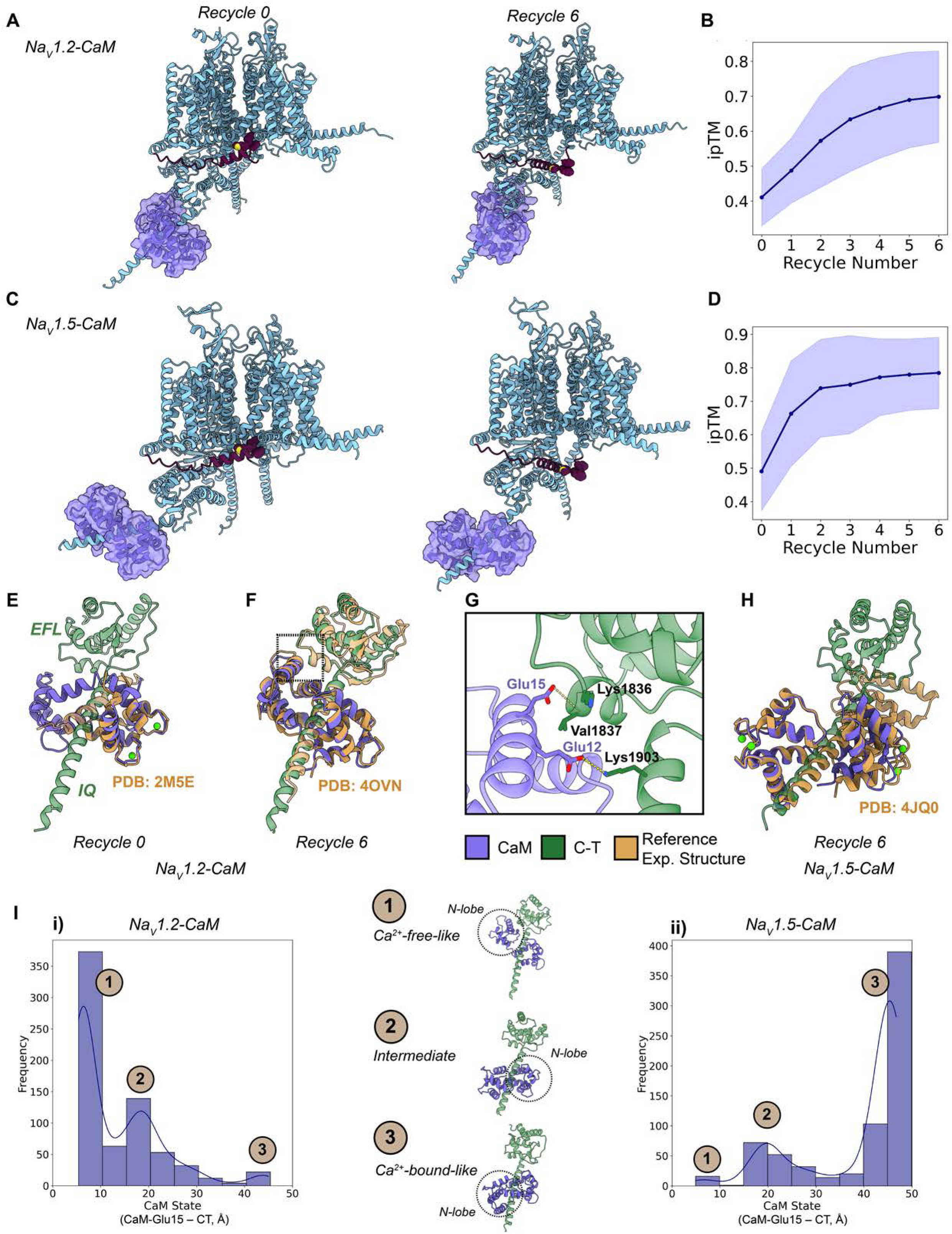
Analysis of models generated with AlphaFold Multimer of CaM bound to hNa_V_1.2 and hNa_V_1.5. (A, C) Overview of CaM-α-subunit complex top models for hNa_V_1.2 (A) and hNa_V_1.5 (B) at recycle 0 and recycle 6. The IFM motif is highlighted in purple. Unstructured intracellular loops are hidden for clarity. (B, D) Evolution across recycles of the average iPTM values of all generated models of CaM bound to hNa_V_1.2 (B) and hNa_V_1.5 (D); shaded areas indicate the standard deviation. (E) Detailed view of top CaM-hNa_V_1.2 model (purple, green) at recycle 0 superimposed into the experimental structure (orange) of Ca^2+^-saturated-CaM bound to the IQ motif of Na_V_1.2 (PDB: 2M5E, (Hovey et al., 2017)). (F) Detailed view of top CaM-hNa_V_1.2 model (purple, green) at recycle 6 superimposed into the experimental structure (orange) of Ca^2+^-free-CaM bound to the IQ motif of Na_V_1.5 (PDB: 4OVN, (Gabelli et al., 2014)). (G) Interactions between CaM and the C-T of the α-subunit in the area highlighted with a dashed square in (F). (H) Detailed view of top CaM-hNa_V_1.5 model (purple, green) at recycle 6 superimposed into the experimental structure (orange) of Ca^2+^-saturated-CaM bound to the IQ motif of Na_V_1.5 (PDB: 4JQ0, (Wang et al., 2014)). (I) CaM state distributions defined by the CaM-Glu15 – CT distance for hNa_V_1.2 (i) and hNa_V_1.5 (ii); representative models of the three identified states are shown.

Notably, CaM displayed conformational changes across recycles for hNa_V_1.2. We observed two distinct states between recycle 0 and recycle 6 (Fig. 8 E and F; Video S7). In recycle 0, the N-lobe of CaM adopts a conformation similar to the Ca²⁺-bound experimental structure (PDB: 2M5E, Ca^2+^-saturated-CaM bound to the IQ motif of Na_V_1.2) (Hovey et al., 2017), while the C-lobe does not. By recycle 6, CaM exhibits a conformation corresponding to the Ca²⁺-free experimental structure (PDB: 4OVN, Ca^2+^-free-CaM bound to the IQ motif of Na_V_1.5) (Gabelli et al., 2014), where the N-lobe interacts with the EFL domain of the C-T region of hNa_V_1.2 (Fig. 8 G). For hNa_V_1.5, CaM remains in the same conformation across recycles in the top model. Compared with the reference experimental structure (PDB: 4JQ0, Ca^2+^-saturated-CaM bound to the IQ motif of Na_V_1.5) (Wang et al., 2014), this state resembles the fully Ca²⁺-bound conformation of CaM bound to the IQ motif (Fig. 8 H).

To further understand these differences, we defined a coordinate distance between Glu15 of CaM and a specific residue (Val1837 in hNa_V_1.2 and the equivalent Ile1833 in hNa_V_1.5) in the EFL domain of the α-subunit C-T domain (Fig. 8 G). This distance (CaM-Glu15 – CT) helps to determine the CaM state: it is shorter in the Ca²⁺-free-like state when the N-lobe interacts with the C-terminal, and larger when CaM is in the Ca²⁺-bound-like state, where the N-lobe no longer interacts. Using this metric, we observed distributions of CaM states for hNa_V_1.2 and hNa_V_1.5 (Fig. 8 I). Three distinct states emerged: (1) a Ca²⁺-free-like CaM state with low CaM-Glu15 – CT distances, where the N-lobe interacts with the C-terminal; (2) an intermediate CaM state, with one lobe Ca²⁺-bound-like and the other not; and (3) a fully Ca²⁺-bound-like CaM state in both lobes. AlphaFold Multimer models of hNa_V_1.2-CaM and hNa_V_1.5-CaM sampled all three CaM states. Notably, the Ca²⁺-free-like CaM state was predominant for hNa_V_1.2 in agreement with experimental data demonstrating that the Ca²⁺-free CaM has a higher affinity for hNa_V_1.2 compared to the Ca²⁺-bound CaM (Feldkamp et al., 2011; Hovey et al., 2017; Mahling et al., 2021). In contrast, the Ca²⁺-bound-like CaM state was more frequently sampled for hNa_V_1.5, in agreement with experimental structure of CaM-Na_V_1.5 IQ domain complex at a high Ca^2+^concentration (PDB: 4JQ0), showing that the N-lobe of CaM was induced to interact with the distal IQ domain of Na_V_1.5, and experimental structure of CaM-Na_V_1.5 IQ domain complex without Ca^2+^ (PDB: 4DCK), showing that the N-lobe of CaM did not make contact with the EFL domain (Wu and Hong, 2021). We also analyzed how these CaM states varied with the recycle number (Fig. S9 AB, i). Consistent with previous observations, recycle 0 showed the greatest conformational diversity. As the number of recycles increased, CaM converged to a predominant state: the Ca²⁺-free state for hNa_V_1.2, and the Ca²⁺-bound state for hNa_V_1.5.

We also assessed the relationship between CaM states and ipTM values (Fig. S9 AB, ii). Extreme states (either Ca²⁺-free or Ca²⁺-bound) showed higher ipTM values. For hNa_V_1.2, the Ca²⁺-free state reached ipTM values above 0.8, while the Ca²⁺-bound state was slightly lower. Conversely, hNa_V_1.5’s Ca²⁺-bound state achieved ipTM values up to 0.9. These differences in maximum ipTM values may partly explain the preferential sampling of specific states for each Na_V_ channel. Lastly, we examined how the presence of CaM influences the distribution of IFM motif states (Fig. S9 AB, iii). For both hNa_V_1.2 and hNa_V_1.5, the presence of CaM increased the sampling frequency of the unbound IFM motif state, as we expected from the visual analysis of the models. Notably, the presence of CaM also affected the state distributions of other channel regions (Fig. S9 AB, iv-vi).

Remarkably, AlphaFold2 predicts CaM’s interaction with Na_V_ channels in the absence of full experimental reference structures of Na_V_ channel – CaM complexes and the absence of Ca²⁺ ions in the modeling. Our models of Na_V_ channel – CaM complexes agree well with known structures of CaM in different states (Feldkamp et al., 2011; Hovey et al., 2017; Mahling et al., 2021; Wu and Hong, 2021), but for a more definitive understanding, future modeling efforts incorporating Ca²⁺ ions will be essential.

### Protein partners reshape correlations among α-subunit state coordinates

Motivated by our observation that the presence of protein partners in the modeling reshapes the conformational landscape sampled by the α-subunit, we next asked whether their presence also alters the relationships between state coordinates themselves (i.e., the state– state correlations within the α-subunit regions). To address this question, we recomputed the full pairwise dependence metrics (Spearman’s ρ and Mutual Information) for the four partner settings: hNa_V_1.1±β1–β4, hNa_V_1.7±β1–β4, hNa_V_1.2±CaM, and hNa_V_1.5±CaM (Fig. S10).

The presence of β-subunits systematically strengthened VSD–VSD correlations (Fig. S10 A). For VSDII–VSDIII and VSDI-VSDIII, ρ increased from ∼0.30–0.45 in α-only models to as high as ∼0.70 with βs (strongest for hNa_V_1.7) (Fig. S10 B, i). The most striking change was the emergence of a robust VSDI–VSDII coupling, negligible in α-only ensembles but rising to ρ≈0.5–0.7 (MI up to ∼0.6) with βs (Fig. S10 B, ii). VSD–AG correlations also became stronger, especially VSDII–AG2, which shifted from ∼0 to ρ≈0.4. The VSDIV–IFM relation became more negative, consistent with increased prevalence of IFM-bound models when βs are present. However, this trend largely reflects state occupancy rather than a strictly monotonic dependence (Fig. S10 B, iii). Presence of CaM produced analogous reshaping of VSD-VSD coupling although to different extents for each pair (Fig. S10 C). In hNa_V_1.2, VSDIII–VSDIV flipped from a negative association (ρ=−0.36) in α-only models to a positive one with CaM (ρ=0.57) (Fig. S10 D, i). Likewise, VSDI–VSDIV switched from ρ=−0.24 to ρ=0.83 with CaM (Fig. S10 D, ii). Similar increases were also observed across other VSD–VSD pairs and in VSD– AG correlations (Fig. S10 D, iii).

Together, these analyses show that protein partners not only shift state occupancies in the modeled ensemble, but they also reshape correlations among α-subunit state coordinates, most notably strengthening correlations among the VSDs and VSD-AG. While the ultimate real implications of these observations will require targeted experiments or physics-based simulations, these results highlight the importance of explicitly considering protein partners when modeling Na_V_ channels with deep learning methods like AlphaFold2, as their presence substantially reshapes the conformational ensemble and state relationships within the α-subunit.

## Discussion

With the emergence of deep learning methods, the field of structural biology has undergone a profound transformation (Baek and Baker, 2022). This evolution is primarily driven by increased computational capabilities, particularly involving the development of advanced neural network algorithms, the substantial growth of protein structure and sequence databases that enable effective training of these models, and GPU-based computation. With the release of AlphaFold2 in 2020 (Jumper et al., 2021), the protein structure prediction accuracy significantly improved, although modeling protein folding remains challenging (Chen et al., 2023). Therefore, the next frontier in computational structural biology lies in predicting the full range of protein structure conformational ensembles, as proteins are inherently dynamic and undergo complex structural changes that govern their diverse functions (Lane, 2023). This challenge of capturing different protein conformations inspired the addition of a new section for predicting protein conformations in the Critical Assessment of Techniques for Protein Structure Prediction (CASP) in 2022 (Kryshtafovych et al., 2023). If a protein sequence determines structure and structure represents an ensemble of conformations or states, then understanding the conformational ensemble, both structurally and energetically, represents the next complex step to fully comprehend the structural changes a complex protein, like an ion channel, undergoes during its functional cycle. This structural modeling challenge is directly relevant to Na_V_ channels, whose conformational landscape is also dependent on the membrane potential, adding another layer of complexity.

We know that the energetic distributions of protein states are influenced by interactions with other partners, such as proteins, lipids, small molecules, and cations, and are also modulated by posttranslational modifications. This adds another dimension to the challenge, as understanding how these interactions influence the equilibrium of the different protein states is required for fully defining the protein’s function and regulation in its physiological context. Additionally, establishing how these interactions reshape the conformational landscape of a protein is particularly important for drug design. The most effective approach for achieving a targeted effect in drug development often involves focusing on a specific conformational state and determining how drug binding influences the protein’s function by altering its state distributions.

Several approaches are currently being explored to apply deep learning methods to solve the problem of predicting protein conformational distributions. First, some methods use deep learning to extract multiple conformations from experimental data (Ed et al., 2021; Punjani and Fleet, 2023; Wankowicz et al., 2024). Second, other deep learning methods aim to predict protein conformational ensembles using algorithms predominantly derived from AlphaFold, where modifications to the MSA construction are made to promote more diverse conformational sampling (Stein and Mchaourab, 2022; Wayment-Steele et al., 2023; Sala et al., 2023; Monteiro da Silva et al., 2024).

In this study, we explored the capabilities of AlphaFold2 to model conformational diversity in different structural regions of Na_V_ channels by employing a subsampled MSA approach (Monteiro da Silva et al., 2024). Structural data has already provided a diverse view of Na_V_ channels conformational dynamics, and we aimed to assess how well the conformations generated by AlphaFold2 capture the diversity of states observed in experimental structures. Additionally, we used AlphaFold Multimer (Evans et al., 2022) to evaluate the ability of these methods to accurately model the interactions between the Na_V_ α-subunit and protein partners, such as auxiliary Na_V_ β-subunits and CaM. We also examined how these interactions influence the conformational landscape of the Na_V_ α-subunit.

We first examined whether AlphaFold2 could generate multiple conformational states for key functional regions of the Na_V_ α-subunit. Focusing on the four VSDs, the IFM motif, the SF, and the AG (Fig. 1), we observed substantial conformational diversity across all nine human Na_V_ subtypes. For each VSD, AlphaFold2 sampled at least two discrete states consistent with one S4 gating charge translocation, in agreement with established activation mechanisms (Fig. 2). VSDIV displayed the broadest conformational range, up to two gating charge translocations (Fig. 2 D), likely reflecting its representation in experimental structures, particularly toxin-stabilized deactivated states (Clairfeuille et al., 2019). In contrast, VSDII did not reach deeply deactivated conformations observed experimentally (Xu et al., 2019; Wisedchaisri et al., 2020), even when a custom template bias was applied (Fig. 2 B; Fig. S2 N-P). These observations suggest an intrinsic limitation in AlphaFold2’s ability to sample deactivated VSD states, potentially reflecting biases in training data. Distributions were largely consistent across subtypes, with the expected exception of Na_X_, which lacks canonical voltage sensitivity. For the IFM motif, AlphaFold2 modeled both experimentally observed bound and unbound states (Fig. 3 AB). The relative frequency of these states varied by subtype; notably, Na_V_1.9 favored the unbound configuration, in line with its slow inactivation kinetics (Fig. S1). In the SF (Fig. 3 C-E), most sampled conformations matched conductive pore states from cryo-EM structures (Fig. 3 D), though small dilations of the DEKA motif resulted in a small number of outlier models with a potentially slow-inactivated-like conformation not experimentally observed (Fig. 3 E). Future modeling incorporating explicit ions may clarify the relevance of these states. Finally, a continuum of AG conformations was sampled (Fig. 3F-J), including rare wide-open states. These would be difficult to observe experimentally due to rapid channel inactivation, underscoring the value of computational modeling in proposing potential transient states that will serve as structural hypothesis to be validated in experiments.

The pLDDT is a metric reported by AlphaFold2 that indicates the confidence of the generated models at the residue backbone level. In some cases, it has been linked to conformational flexibility (Wilson et al., 2022; Vander Meersche et al., 2025). However, some studies show there is not a clear correlation between pLDDT and experimental B-factors, which represent atomic mobility or uncertainty in experimental structures (Carugo, 2023). Although the actual physical implications of the pLDDT values are still unclear and should be interpreted cautiously, it is one of the primary approaches to evaluate models generated by AlphaFold2. We observed specific distributions of pLDDT for each of the state coordinates of these regions (Fig. S2 A-G). Understanding the relationship between pLDDT, state distributions, and the actual energetic landscape of protein states remains a major challenge in the field, and it is still unclear whether this information can be directly obtained from AlphaFold2-generated data. However, one clear conclusion from our results is that the greatest conformational diversity occurs at recycle 0 (Fig. S2 I-L). As the recycle number increases, the structures tend toward specific states with high pLDDT values. This raises the question of whether AlphaFold2 sampling of specific states with increased recycles occurs because these states are more commonly seen in the training set or if these states are more energetically stable. However, this is a complex issue because the states we often observe in experimental structures are typically the most energetically stable.

The Na_V_ α-subunit is finely regulated under physiological conditions (Namadurai et al., 2015) by auxiliary Na_V_ β-subunits and intracellular proteins, such as CaM, a Ca^2+^sensor (Wu and Hong, 2021). In our modeling of hNa_V_1.1 and hNa_V_1.7 channels with the four β-subunits, we successfully generated high-accuracy models that agreed well with experimental structures (Fig. 6). This demonstrates that AlphaFold Multimer is capable of accurately sampling the complexes between the α-subunit and β-subunits.. While the transmembrane domains of β-2 and β-4 have not been resolved experimentally, AlphaFold Multimer models predict that they span the membrane, although with a slight tilt relative to the membrane’s normal axis (Fig. 6; Fig. S7). This could be attributed to increased flexibility of these regions, although other factors, such as other protein partners and membrane lipid composition, might also contribute. Our results also demonstrate that the presence of the β-subunits reshapes the modeled conformational landscape of the α-subunit (Fig. 7). In particular, we were able to generate models of VSDIV with more deactivated states than observed experimentally and when modeling the α-subunit alone. These data reveal a possible redistribution of the Na_V_ α-subunit conformational landscape due to the presence of the auxiliary β-subunits, which may modulate channel activity. We see a similar pattern in the case of Na_V_ α-subunit complexes with CaM, where the IFM motif state shows a clear redistribution in the presence of CaM (Fig. S9 AB, iii). For CaM, our models agree well with reference experimental structures of Na_V_ channel regions in complex with CaM (Fig. 8 E-H). However, currently, there are no structures available of a full Na_V_ channel in a complex with CaM. Additionally, modeling CaM presents a challenge due to its Ca^2+^-regulated conformational diversity. We observed conformational changes across the recycles, with a tendency toward different CaM states in the two cases tested (Fig. 8 I; Fig. S9 AB, i). Specifically, for hNa_V_1.2, the Ca^2+^-free state was most frequently sampled, whereas for hNa_V_1.5, the Ca^2+^-bound state was more common, in agreement with experimental functional data and structures (Feldkamp et al., 2011; Hovey et al., 2017; Mahling et al., 2021; Wu and Hong, 2021). Despite the challenges in extracting firm physiologically relevant conclusions from these results, the AlphaFold Multimer models of Na_V_ – CaM complexes might be useful for structural hypothesis formulation and guiding experimental studies aimed at deciphering the molecular mechanisms of Na_V_ channel CaM-dependent modulation. The next step would involve incorporating Ca^2+^ into the modeling, which is possible with AlphaFold3 (Abramson et al., 2024) and RoseTTAFold-3 (Corley et al., 2025).

Extracting firm mechanistic or energetic insights from AI-generated ensembles remains challenging, and most studies in this direction have been conducted in simplified systems with two functional states (Sala et al., 2023; Monteiro da Silva et al., 2024), while Na_V_ channel conformational landscape is far more complex. When modeling the Na_V_ α-subunit alone, we observed robust state–state couplings (Fig. 4), most notably coordinated activation among the VSDs and positive VSDII/VSDIII–AG relationships, largely consistent with known gating logic, alongside patterns (e.g., VSDIV–IFM) that were more complex to interpret. Introducing protein partners reshaped these couplings, with β-subunits and CaM strengthening VSD–VSD correlations and sensor–gate coupling (Fig. S10), though such changes may partly reflect modeling context rather than actual causal mechanisms. Unsupervised clustering of all models yielded a small set of recurrent, interpretable ensembles with subtype-dependent occupancy (Fig. 5). We do not claim that the observed state frequencies reflect *in vivo* occupancies, as sampling limitations, especially for deeply deactivated VSD configurations, and known biases (templates, training data, absence of ions) must be kept in view. Definitive assignments will require more advanced deep learning methods, well-thought experiments and/or physics-based simulations to connect the modelled ensembles to actual molecular mechanisms, energetic parameters and kinetics.

At the same time, our study illustrates why ensemble modeling is the next frontier for voltage-gated ion channels, now that most ion channels have experimental structures. To our knowledge, this is the first attempt to generate full-channel conformational ensembles across all human Na_V_ subtypes and accurately assemble multimer complexes with native partners, offering practical guidance for researchers building models for their own purposes. A recent study similarly reported AlphaFold2’s ability to sample multiple states across the broader voltage-gated ion channel family (Tao and Corry, 2025), underscoring the community’s shift from static structures to conformational ensembles. We also see value in making this rich and well-characterized ensemble repository publicly available for downstream use (e.g., state-specific docking/virtual screening, and allosteric hypothesis testing) by other researchers in the field. Newer deep-learning frameworks (Abramson et al., 2024; Krishna et al., 2024; Corley et al., 2025; Passaro et al., 2025; Wang et al., 2025) that handle heteroatoms/ions and improved sampling schemes should help with the current limitations described here, while continued cross-validation against experiment will keep the modeling grounded and reliable for functional inference.

## Acknowledgments

We dedicate this article to the memory of Dr. William A. Catterall, a creative scientist, supportive mentor, and inspiring collaborator. We thank Drs. Heike Wulff and Jon Sack and members of the VY-Y laboratory for valuable discussions. The work in VY-Y lab was supported by NIH grants R61NS127285, R01HL128537, and R01HL174001. DLM was supported by Fulbright Fellowship and the University of California Davis Center for Precision Medicine and Data Science.

## Data Availability

All the models generated in this study for all test cases are openly available in the Dryad repository: https://doi.org/10.5061/dryad.rn8pk0pn3. (Active link for Reviewers: http://datadryad.org/stash/share/XjQoAjQ7urq1GsHCu1falyK7Rwe_lqobGJEsQPTZhSI).

All the data and analysis scripts can be found in a dedicated Github repository (https://github.com/VYYlab/nav_conf_modeling) with a step-by-step guide to replicate all the results from this study.

## Supplemental Material

### Supplemental Figure Legends

**Figure S1.**
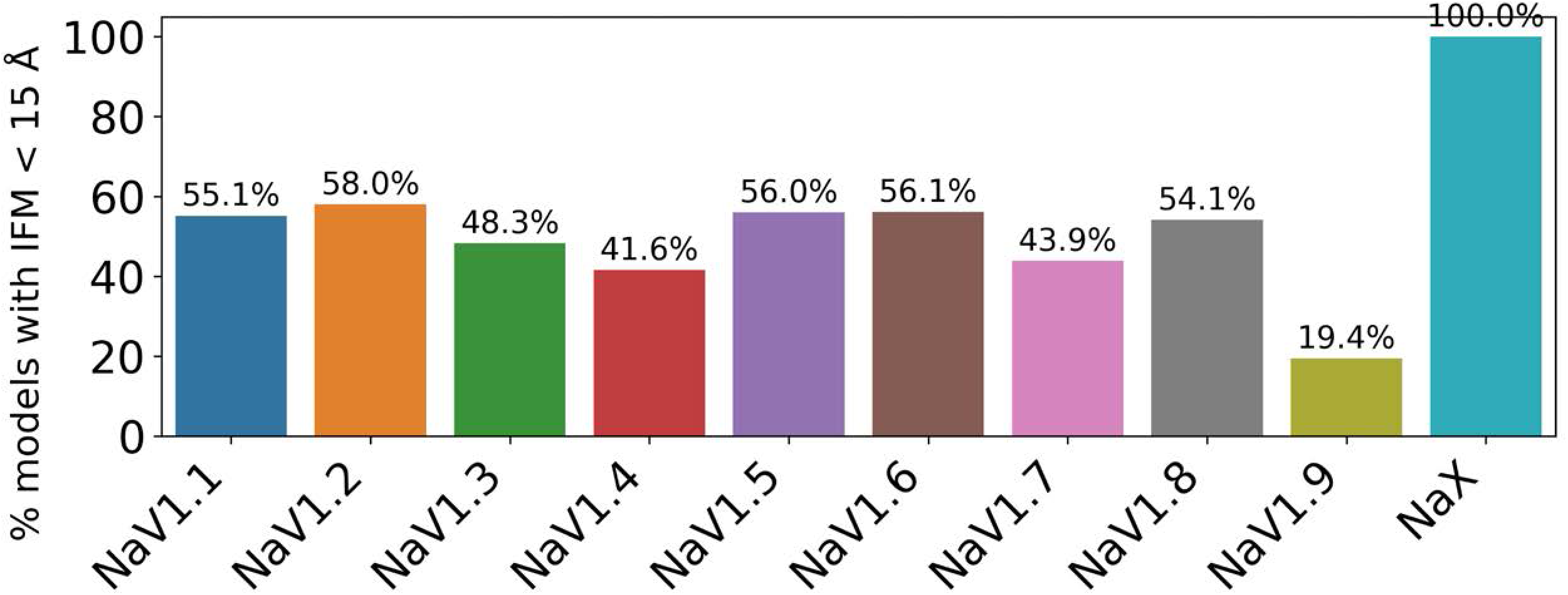
Distribution of IFM motif bound states across hNa_V_ channel subtypes. Percentage of AlphaFold2-generated models per channel subtype in which the IFM motif is positioned within 15 Å of its receptor pocket (IFM–PD distance < 15 Å), corresponding to a “bound” or fast inactivated configuration.

**Figure S2.**
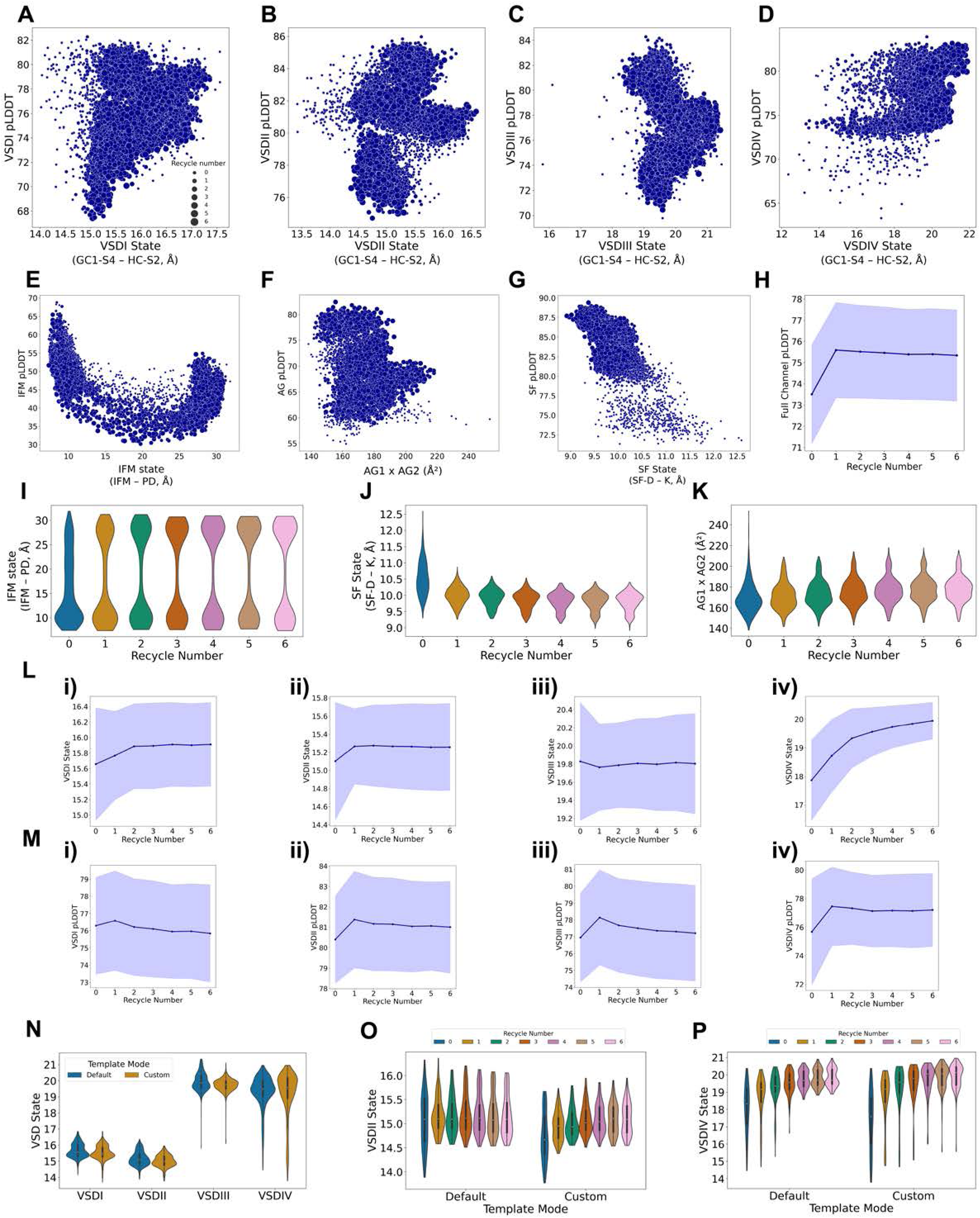
Relationships between state distributions, model pLDDT, number of recycles and use of custom templates. (A-G) Relationship between the state distributions of the seven investigated regions and the corresponding pLDDT values of these regions; dot sizes represent the recycle number. (H) Evolution of the average adjusted-global pLDDT values across recycles; shaded outlines represent standard deviation. (I-L) Evolution of state distributions across recycles for the studied regions. (M) Evolution of average pLDDT values of the four VSDs across recycles; shaded areas indicate the standard deviation. (N) Comparison of the distribution of hNa_V_1.7 VSD states with the default and custom template modes. (O,P) Comparison of the distribution of hNa_V_1.7 VSDII (O) and VSDIV (P) states across recycles with the default and custom template modes.

**Figure S3.**
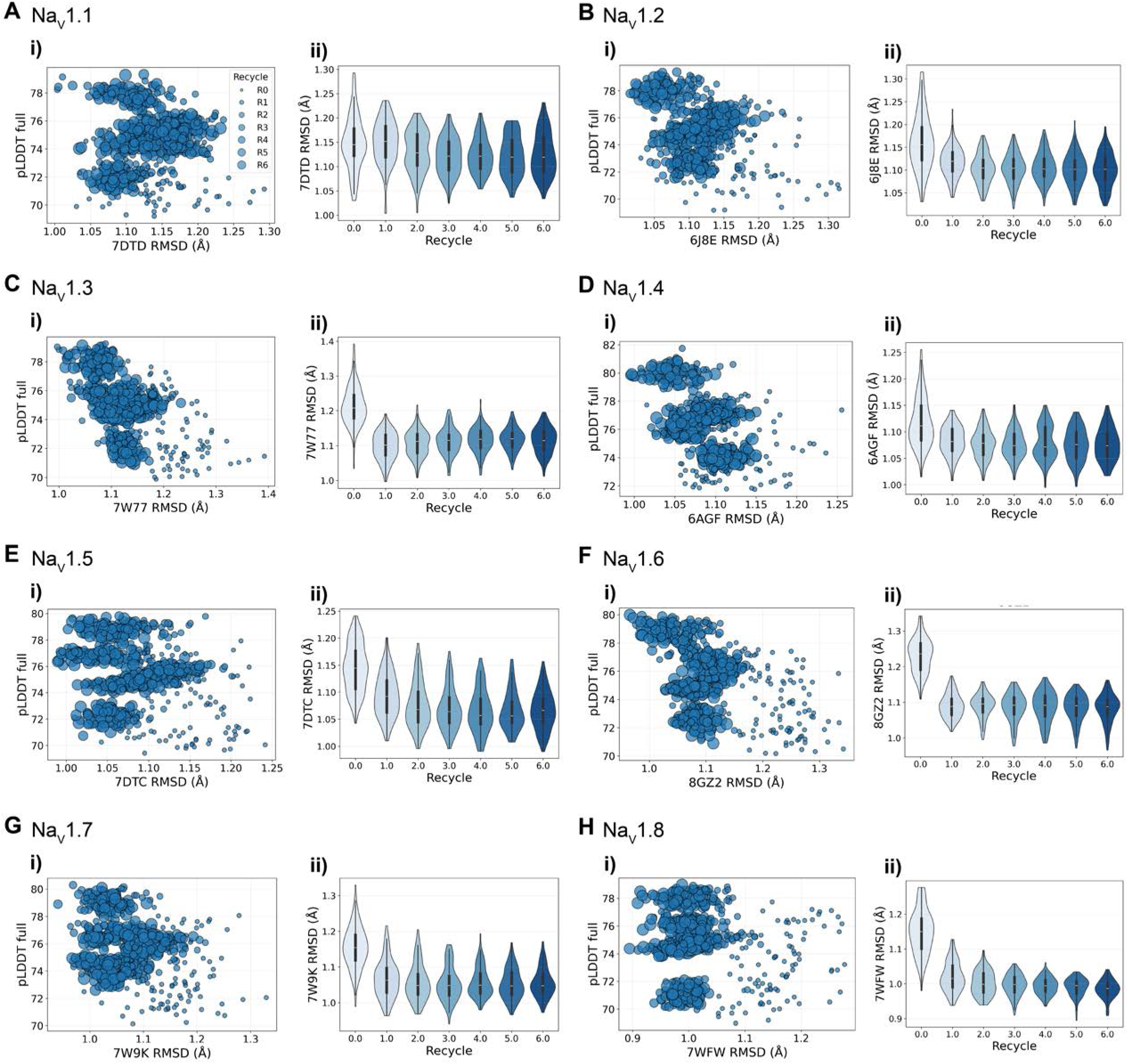
Structural agreement and model confidence across hNa_V_ subtypes. For each human Na_V_ subtype with available experimental structure (hNa_V_1.1–hNa_V_1.8), generated models were compared to corresponding experimental structures. (A–H) Panels show (i) scatter plots of global pLDDT versus backbone RMSD (Å) relative to the subtype-specific experimental structure of reference (indicated in each plot x-axis label) and (ii) distributions of RMSD values as a function of recycle number. Dot sizes in the scatter plots represent recycle number.

**Figure S4.**
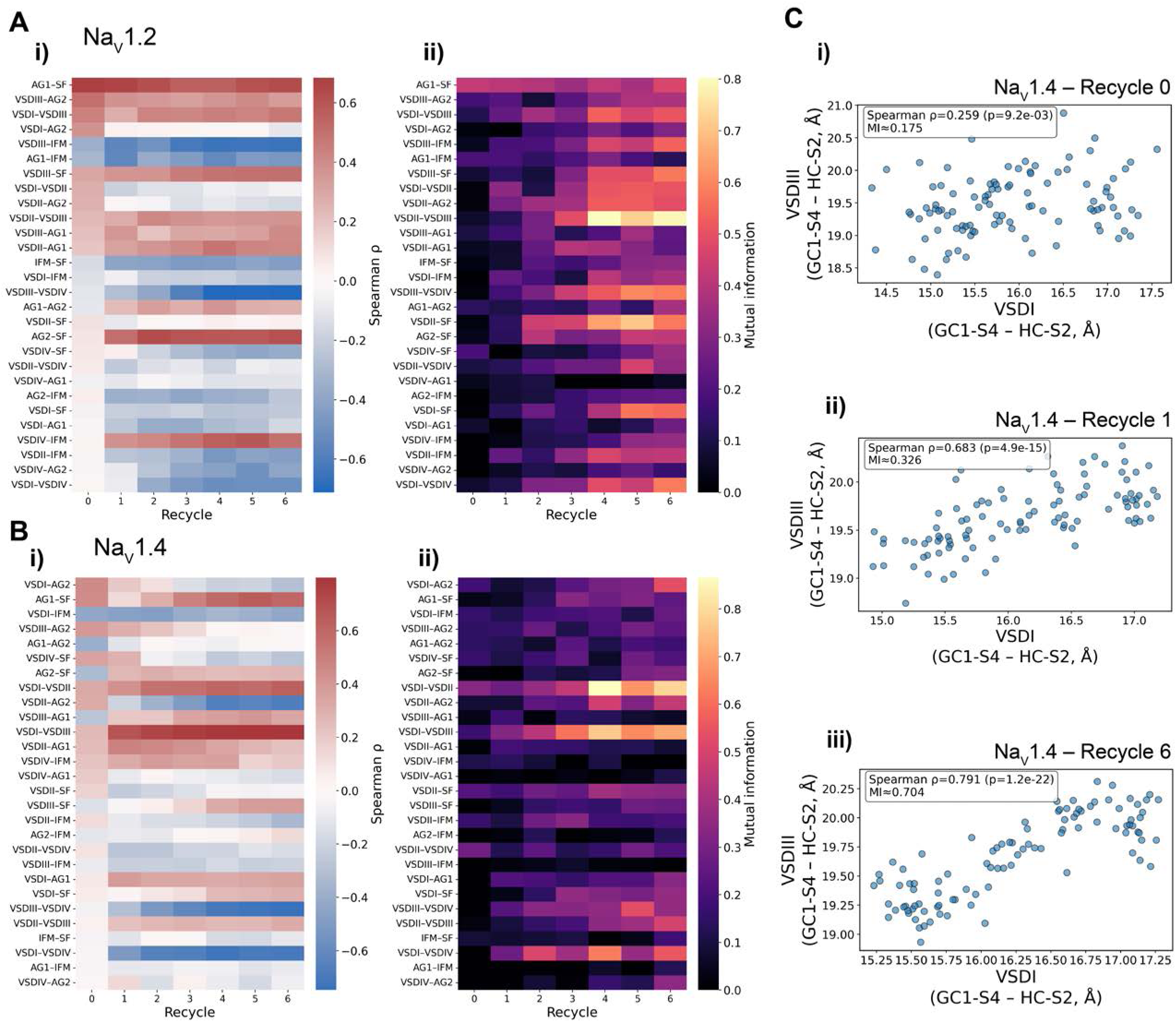
Effect of recycle number on state–state correlations. (A–B) For hNa_V_1.2 and hNa_V_1.4, heatmaps show pairwise relationships among state coordinates across recycles, quantified by (i) Spearman’s ρ (monotonic dependence) and (ii) Mutual Information (non-monotonic dependence). (C) Representative examples from hNa_V_1.4 illustrating how increasing recycles (i–iii) strengthens VSDI–VSDIII coupling.

**Figure S5.**
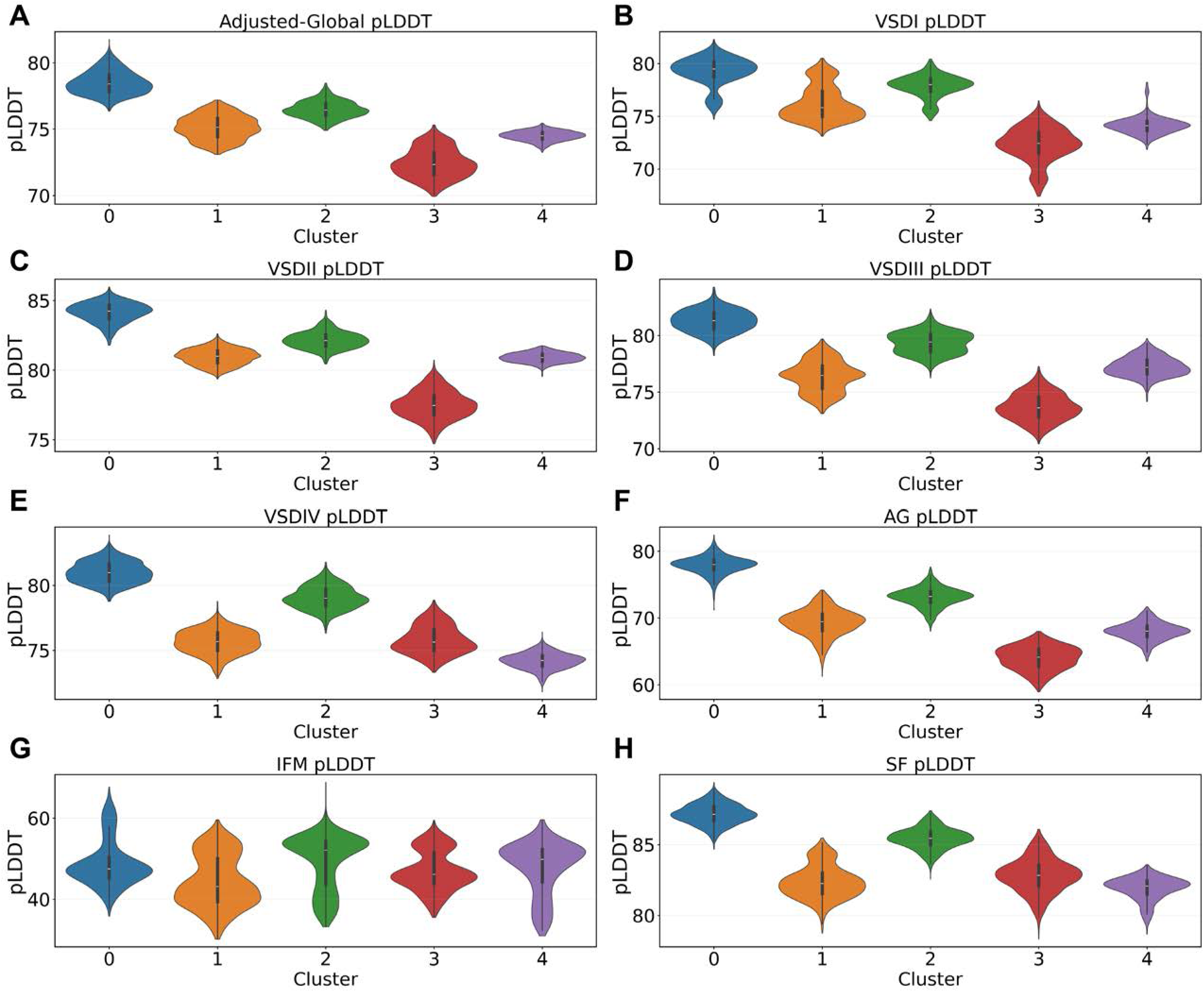
Distribution of confidence values (pLDDT) across conformational clusters. Violin plots show the distributions of pLDDT scores for models grouped by cluster (0–4) as defined in Figure 5. Panels display (A) adjusted global pLDDT and region-specific pLDDT for (B) VSDI, (C) VSDII, (D) VSDIII, (E) VSDIV, (F) activation gate (AG), (G) IFM motif, and (H) selectivity filter (SF).

**Figure S6.**
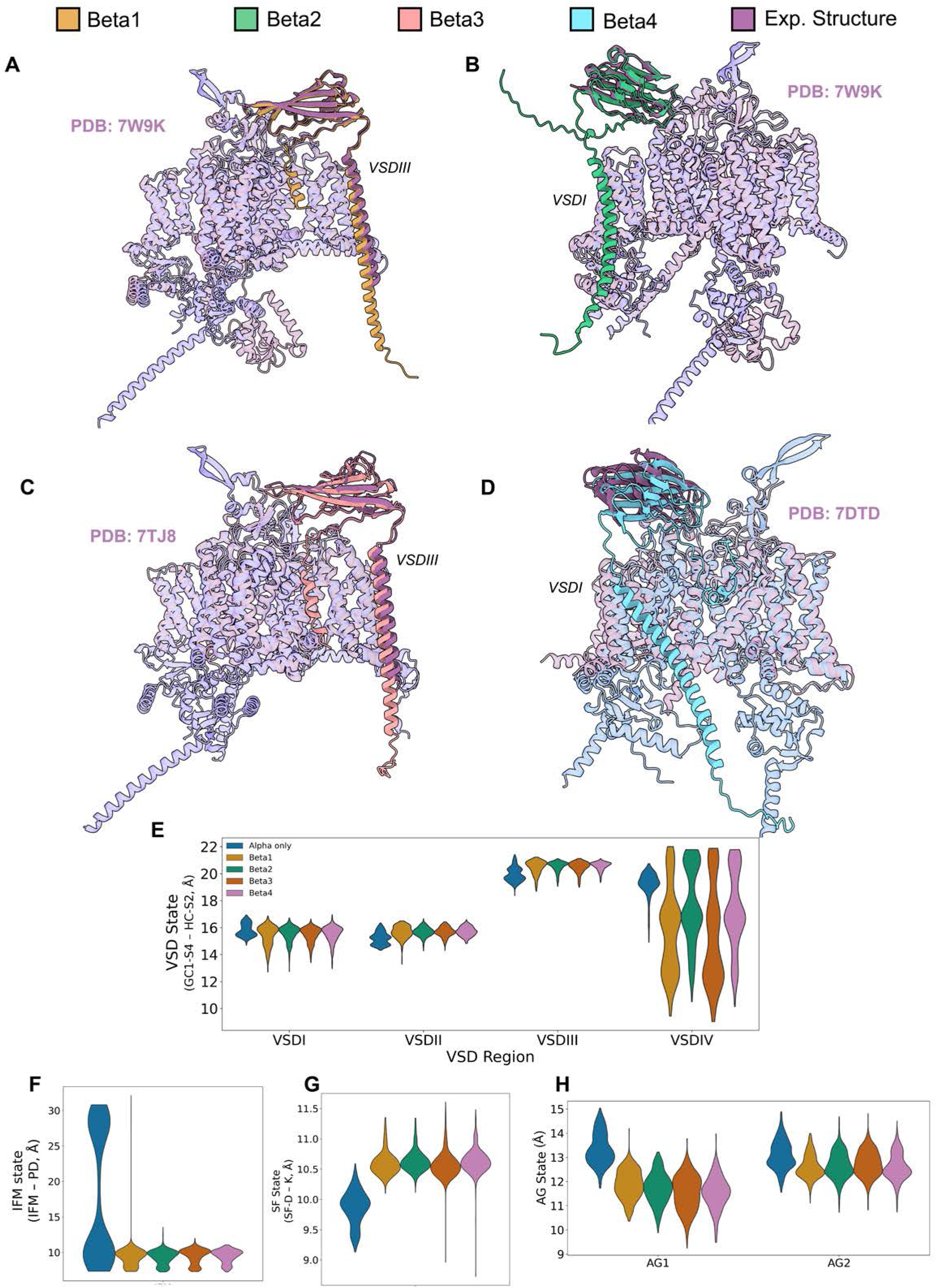
Modeling of hNa_V_1.1 with β-auxiliary subunits. (A-D) Superimposition of top ranked AlphaFold mo_d_els of hNa_V_1.1 in complex with the four β-auxiliary subunits with the corresponding experimental structure o_f_ reference: (A,B) hNa_V_1.7-β1-β2 complex, PDB: 7W9K (Huang, et al., 2022a); (C) hNaX-β3 complex, PDB: 7T_J_8 (Noland, et al., 2022); (D) hNa_V_1.1-β4 complex, PDB: 7DTD (Pan et al., 2021). (E-H) Comparison of the distribution of states of the VSDs (E), IFM motif (F), AG (G), and SF (H) in the presence of the four different β-subunits and when the α-subunit is modeled alone.

**Figure S7.**
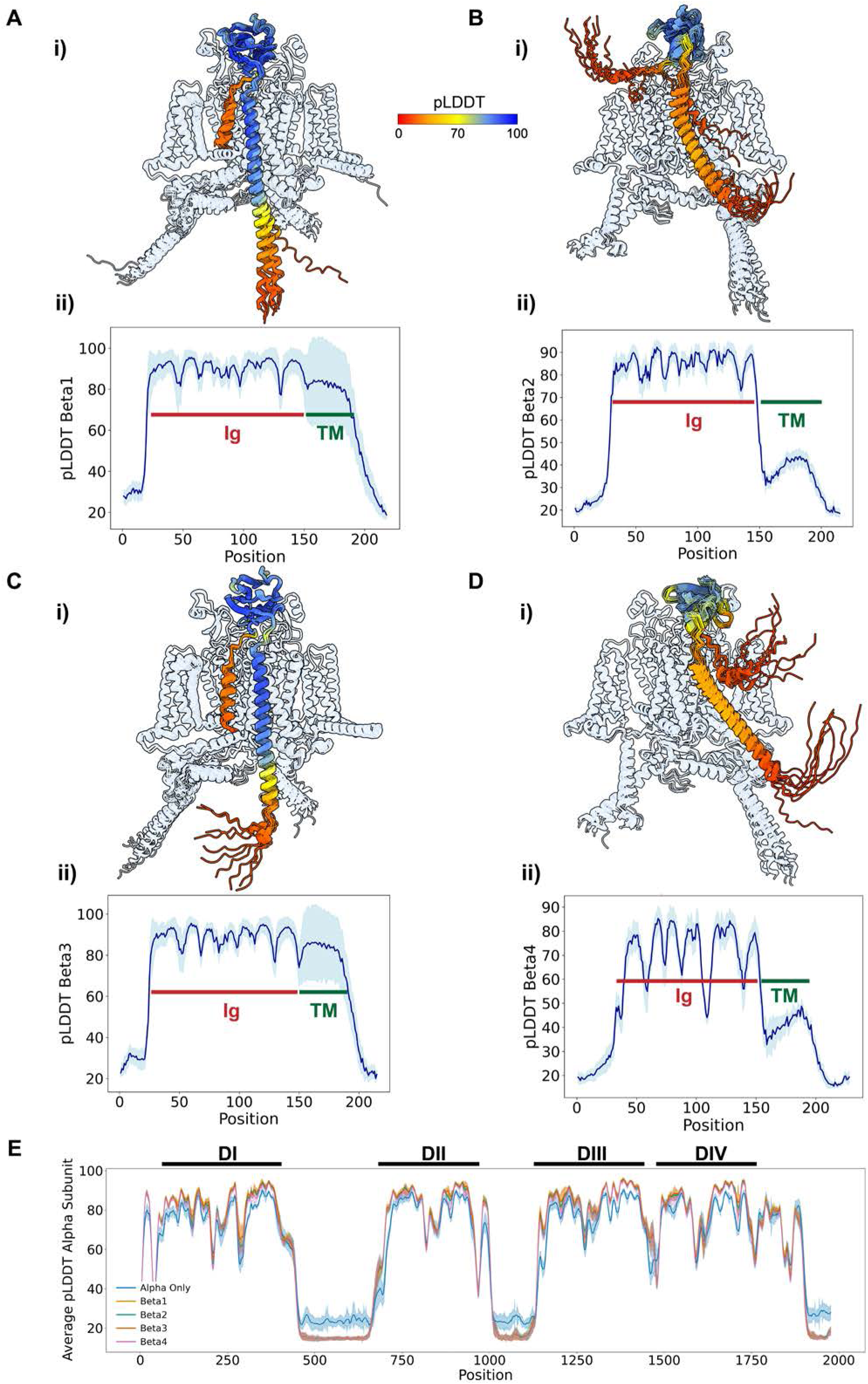
Distribution of pLDDT values across auxiliary β-subunits in complex with hNa_V_1.7. (A-D, i) Lateral view of top 10 models of hNa_V_1.7 in complex with the four auxiliary β-subunits; the α-subunit ribbon is shown with transparency, and the auxiliary β-subunit ribbons are colored by pLDDT values. (A-D, ii) Average pLDDT values per position across the four auxiliary β-subunits in all generated models; shaded areas indicate the standard deviation. (E) Average pLDDT values per position across the α-subunit in all generated models with the α-subunit alone, or in complex with the four auxiliary β-subunits; shaded areas indicate the standard deviation.

**Figure S8.**
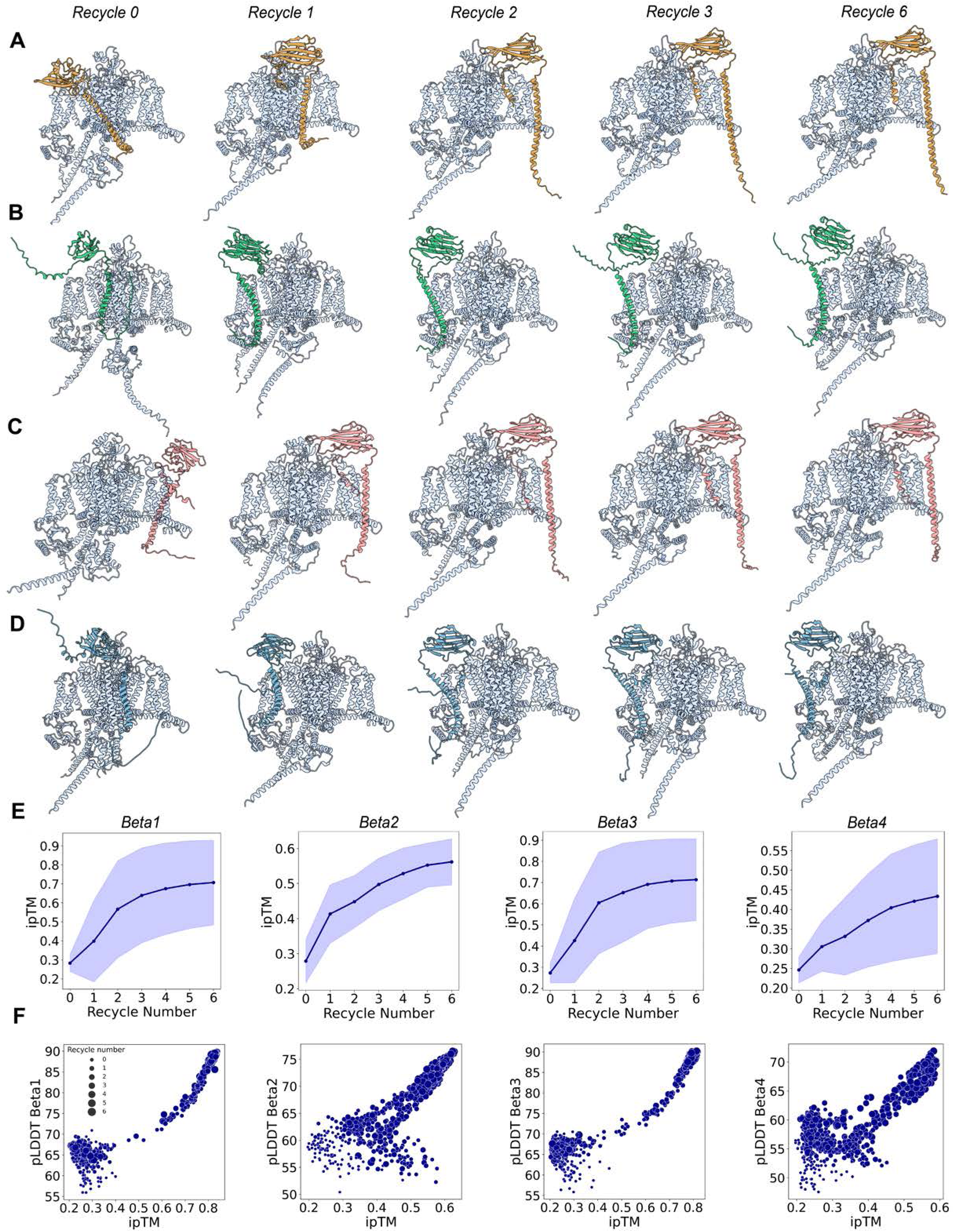
Analysis of the effect of recycles and ipTM values on Na_V_ α-β multimer complex modeling. (A-D) Top model of hNa_V_1.7 in complex with each of the four auxiliary β-subunits across recycles. (E) Evolution across recycles of the average iPTM values of all generated models for each of the four β-subunit cases; shaded areas indicate the standard deviation. (F) Relationship between model iPTM and β-subunit pLDDT values.

**Figure S9.**
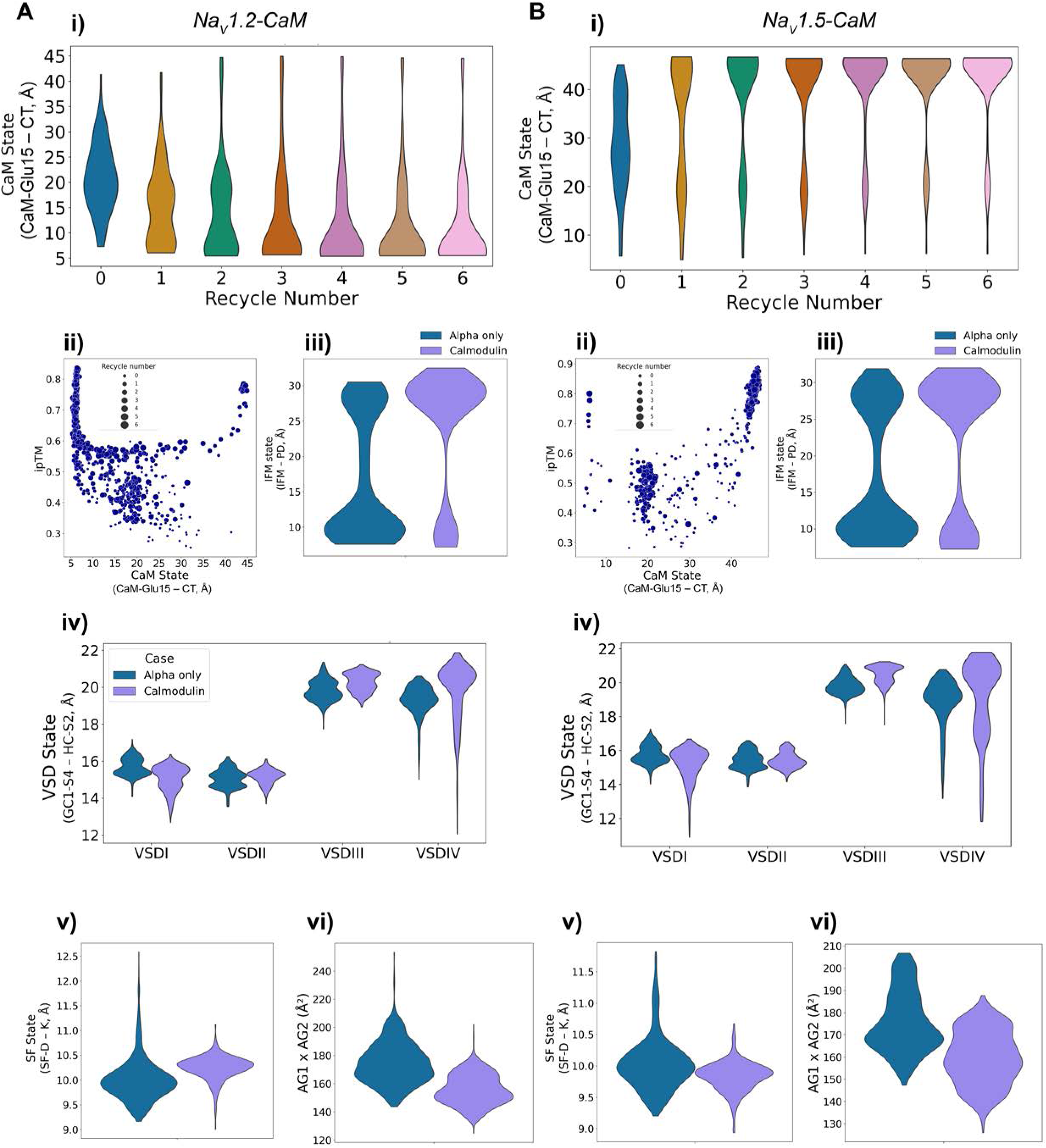
Calmodulin (CaM) state distributions and effects on Na_V_ α-subunit conformations. (A–B) Analyses of CaM-bound models for hNa_V_1.2 (A) and hNa_V_1.5 (B). (i) Violin plots show the distribution of CaM states, defined by the CaM-Glu15 – CT distance, across recycle numbers. (ii) Scatter plots illustrate the relationship between CaM state and model iPTM score. (iii) Comparison of IFM state between α-only and CaM-bound models. (iv) Distribution of VSD state coordinates for each domain (VSD I–IV) in α-only versus CaM-bound models. (v–vi) Comparison of selectivity filter (SF) and activation gate (AG) states between α-only and CaM-bound models.

**Figure S10.**
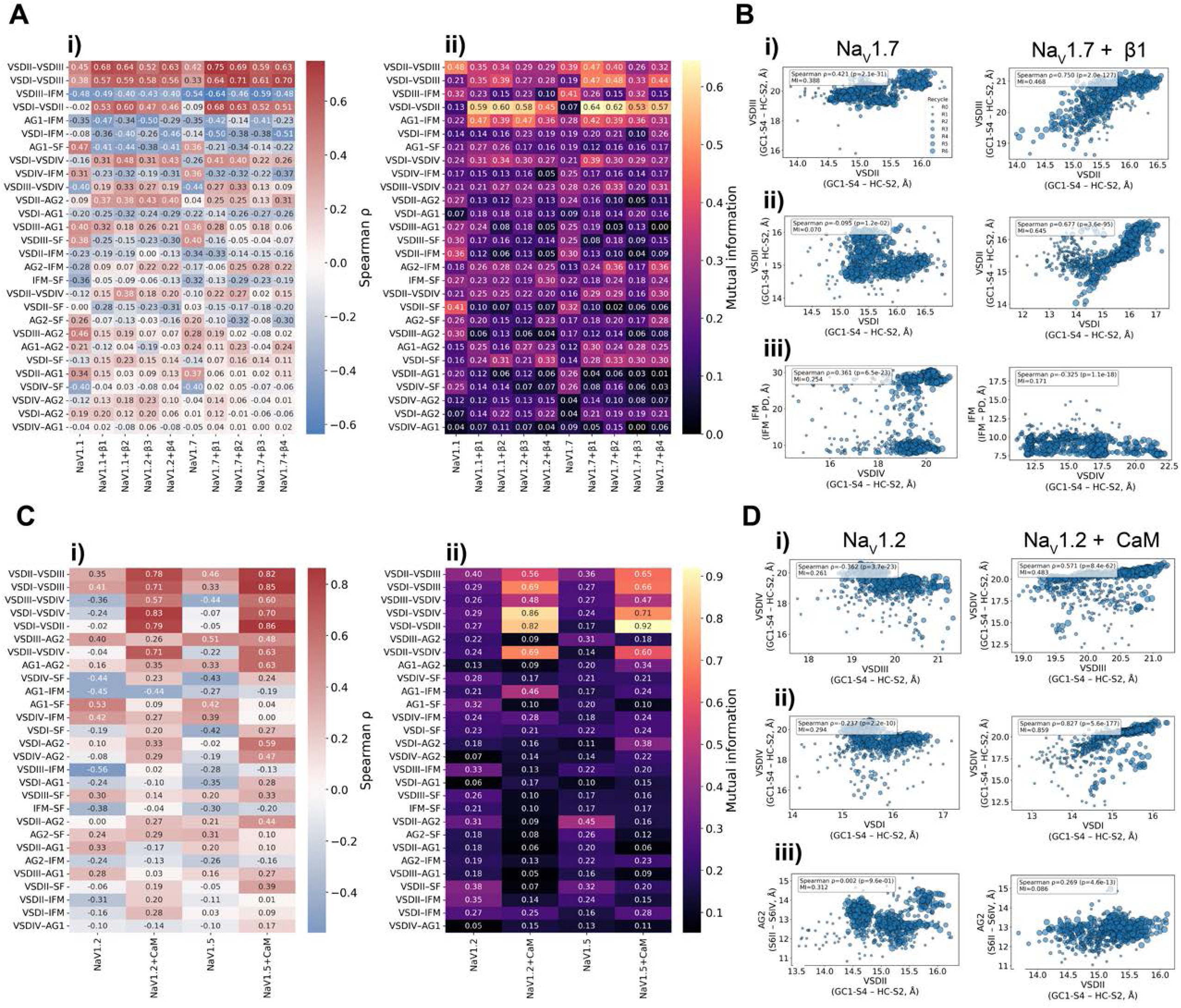
Effect of protein partners on correlations among Na_V_ α-subunit state coordinates. (A–B) Analysis of β-subunit eff_e_cts on hNa_V_1.7. (A) Pairwise relationships among state coordinates for α-only and α+β1–β4 models quantified by (i) Spearman’s ρ (monotonic) and (ii) Mutual Information (non-monotonic). (B) Representative scatter plots illustrate examples of strengthened correlations upon β-subunit inclusion: (i) VSDII–VSDIII, (ii) VSDI–VSDII, and (iii) VSDIV–IFM. (C–D) Analysis of CaM effe_c_ts on hNa_V_1.2 and hNa_V_1.5. (B) Pairwise state relationships computed for α-only and CaM-bound models. (D) Representative scatter plots show selected examples of enhanced VSD–VSD and VSD–AG correlations in CaM-bound complexes: (i) VSDIII–VSDIV, (ii) VSDI–VSDIV, and (iii) VSDII–AG2. Dot sizes represent recycle number.

### Supplemental Table Legends

**Table S1.** List of amino acid sequences of all the proteins used in this study.

**Table S2.** Definition of atom pairs to calculate the distances to use as state coordinates for each region of interest for each of the nine hNa_V_ channels and hNa_X_.

**Table S3.** Reference values of distance coordinates for each region in experimental structures of human Na_V_ channels. Other non-human structures are included to use as reference for particular states. All distances are in Å.

### Supplemental Video Legends

**Video S1.** Morphing between the most deactivated and most activated models for each of the VSDs.

**Video S2.** Morphing between the most contracted and most dilated SF conformations obtained in our models.

**Video S3.** Morphing between different states of the AG: (i) a closed state, (ii) the most open state obtained, (iii) an inactivated state, and (iv) returning to the closed state.

**Video S4.** Morphing across the six recycles of the top models of hNa_V_1.7 bound to the four β-auxiliary subunits.

**Video S5.** Morphing between the most deactivated VSDIV state, obtained in the presence of β-3, and the most activated state.

**Video S6.** Morphing between the three identified states of the SF, being the same as Video S2 plus the new contracted state found in the presence of β-4.

**Video S7.** Morphing across the 6 recycles of the top model of CaM bound to hNa_V_1.2. Only CaM and the C-T of the α-subunit are shown for clarity.

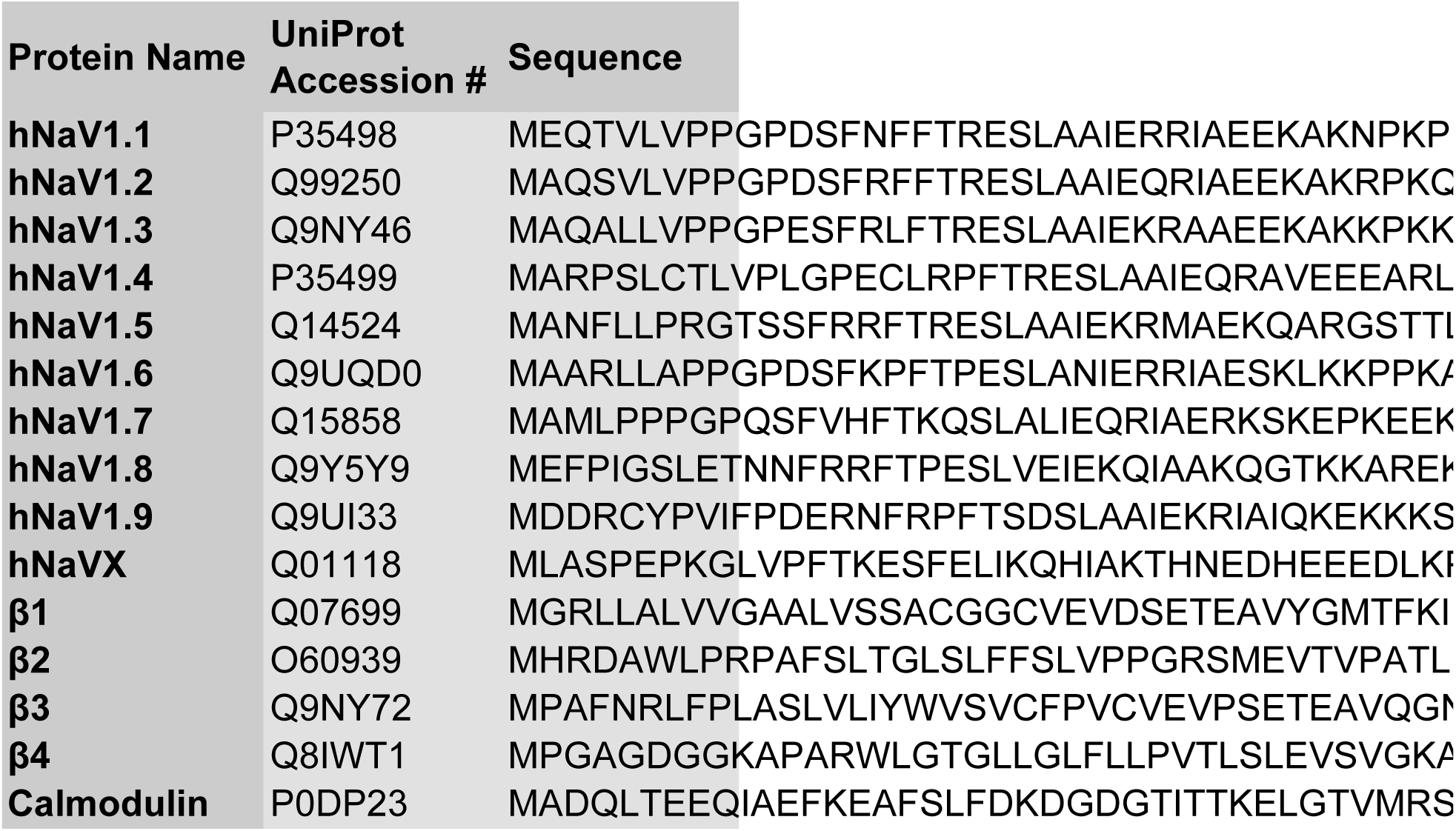
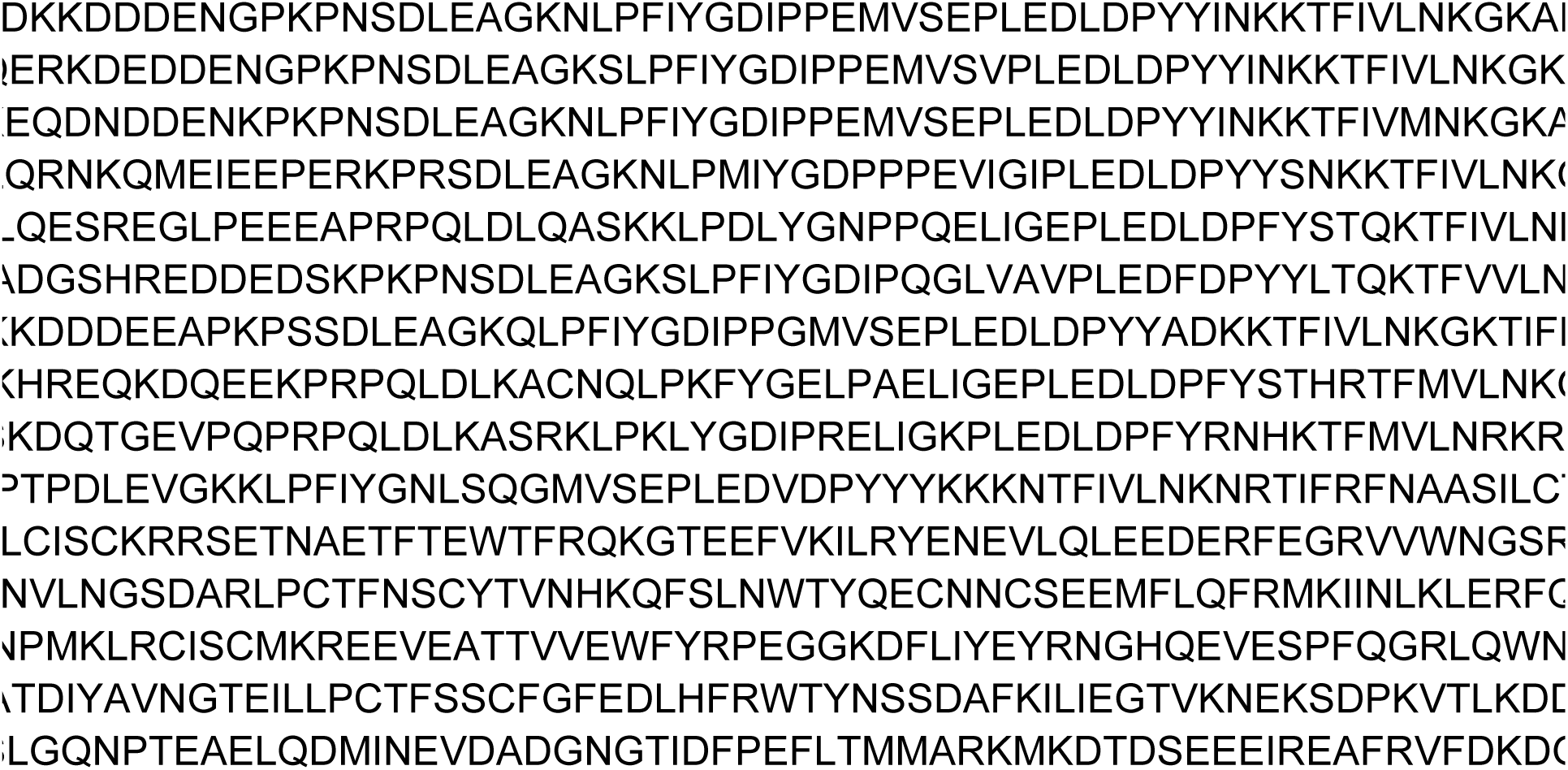
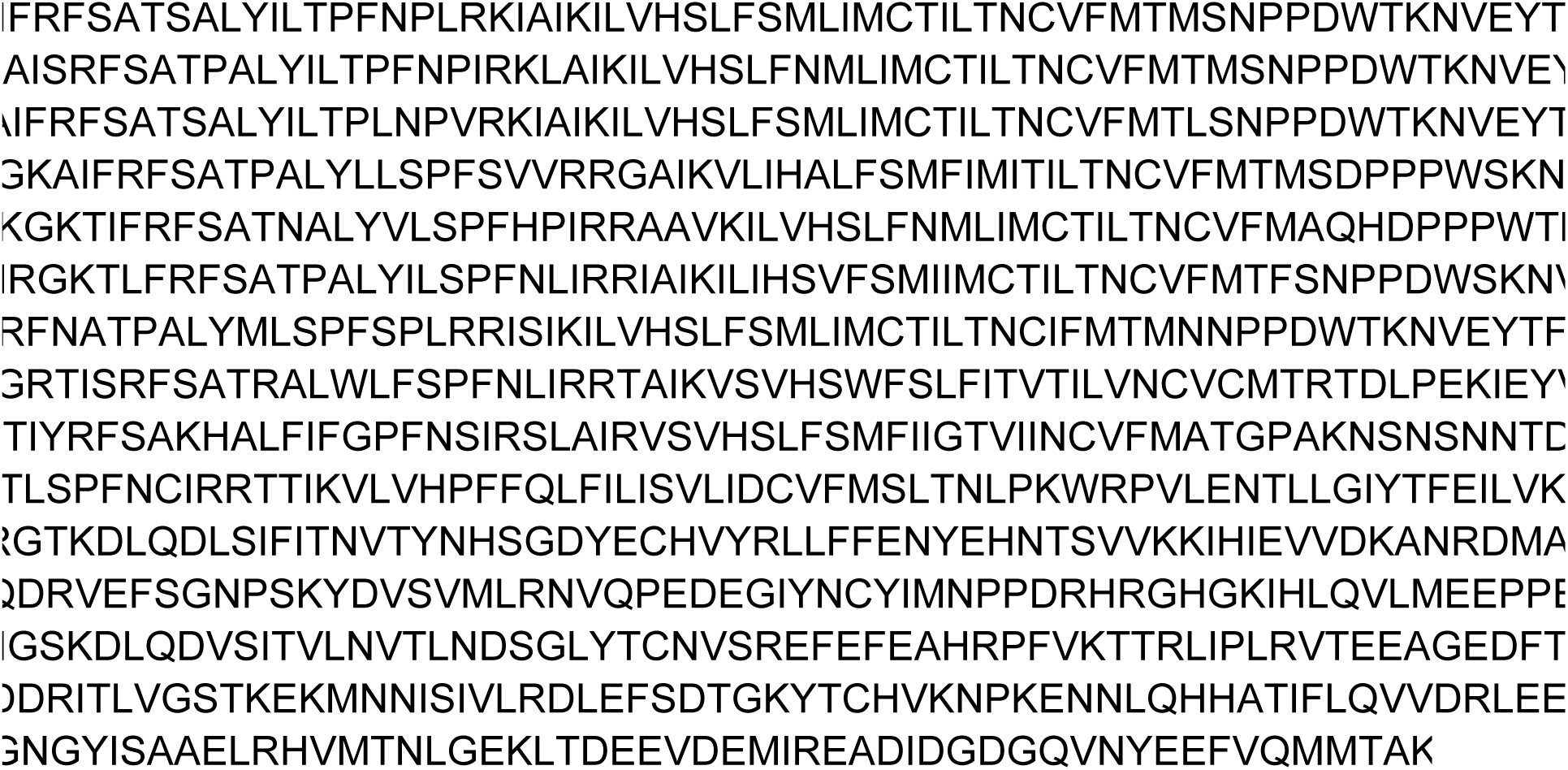
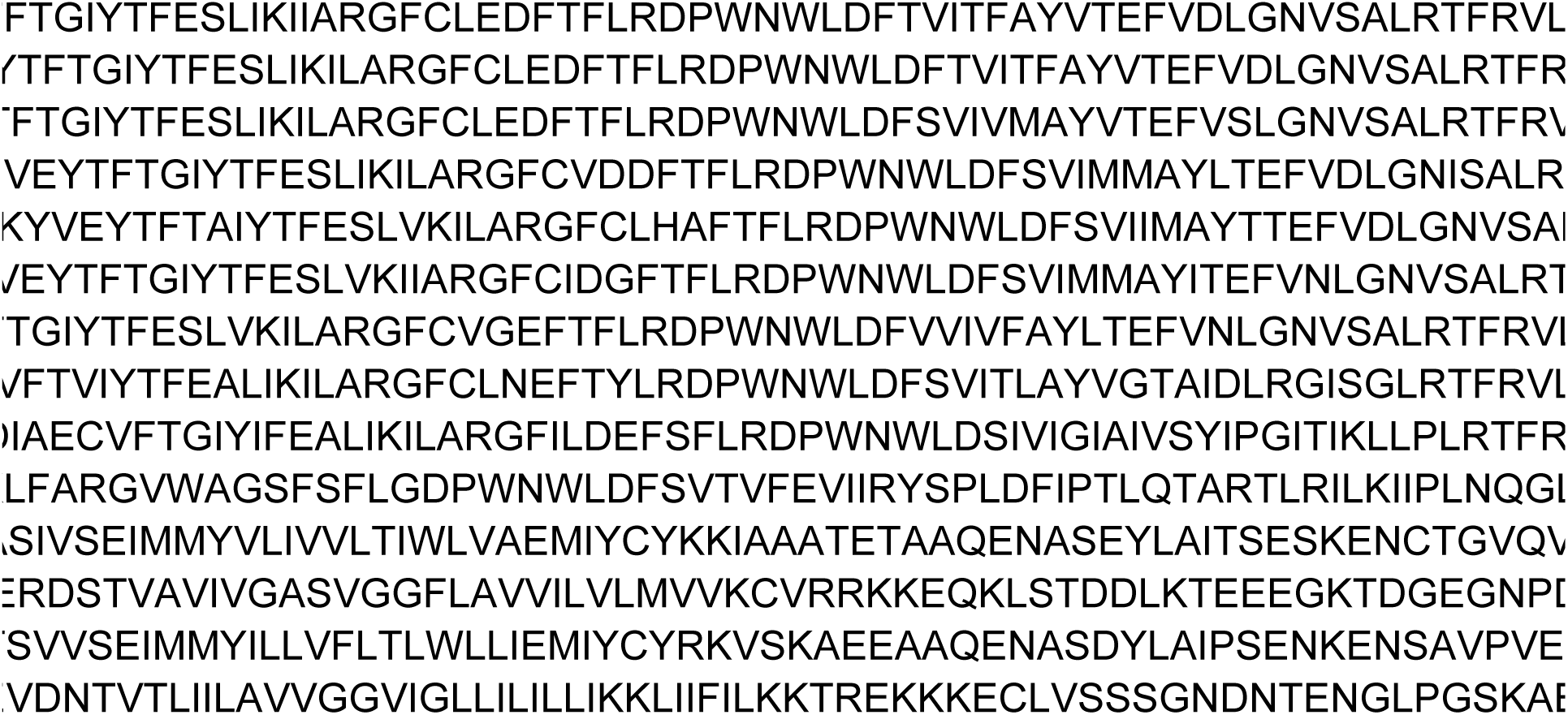
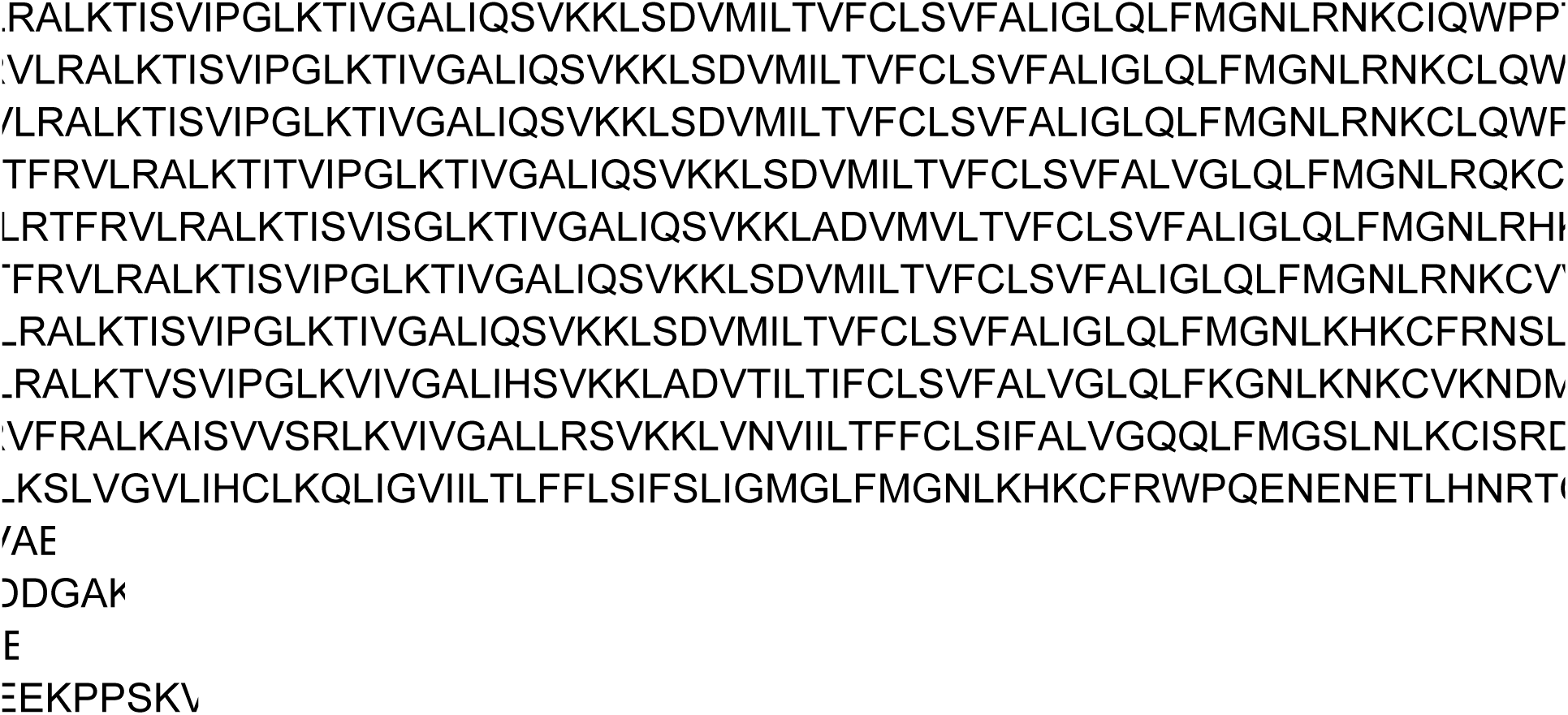
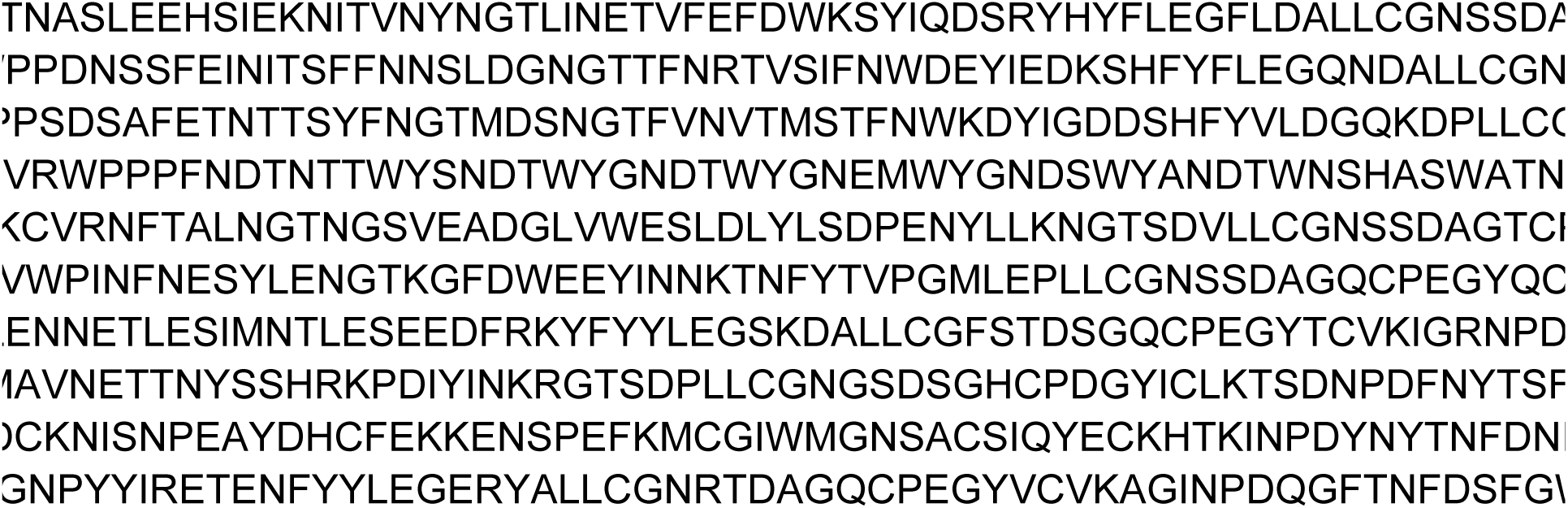
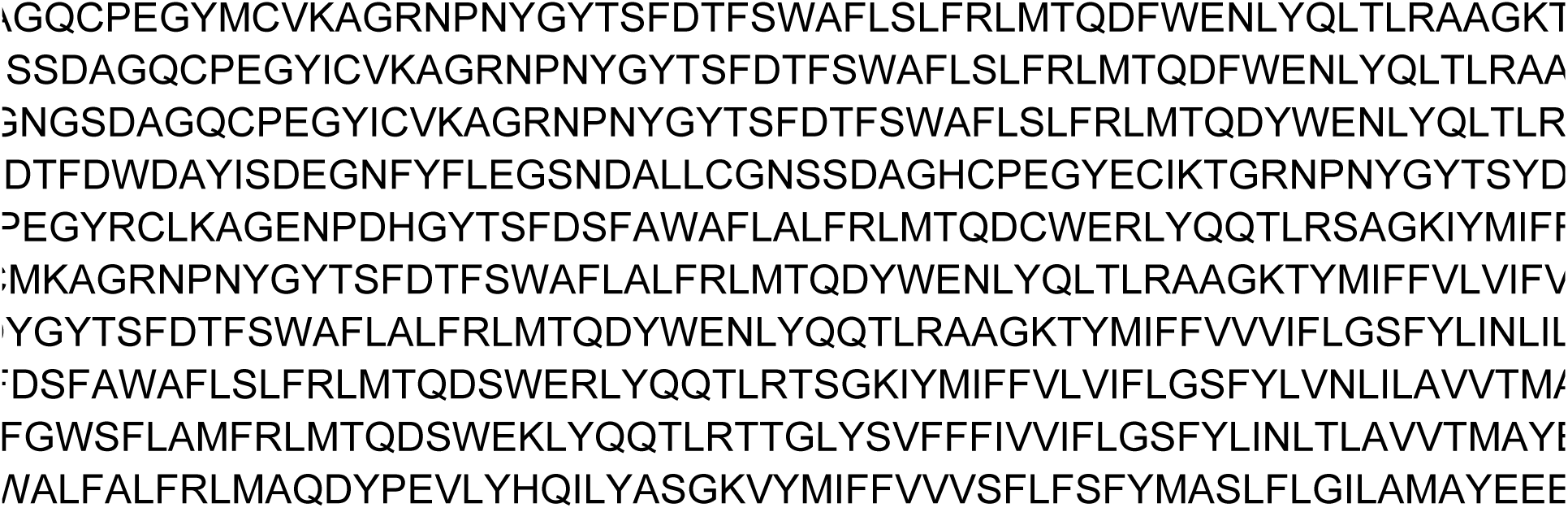
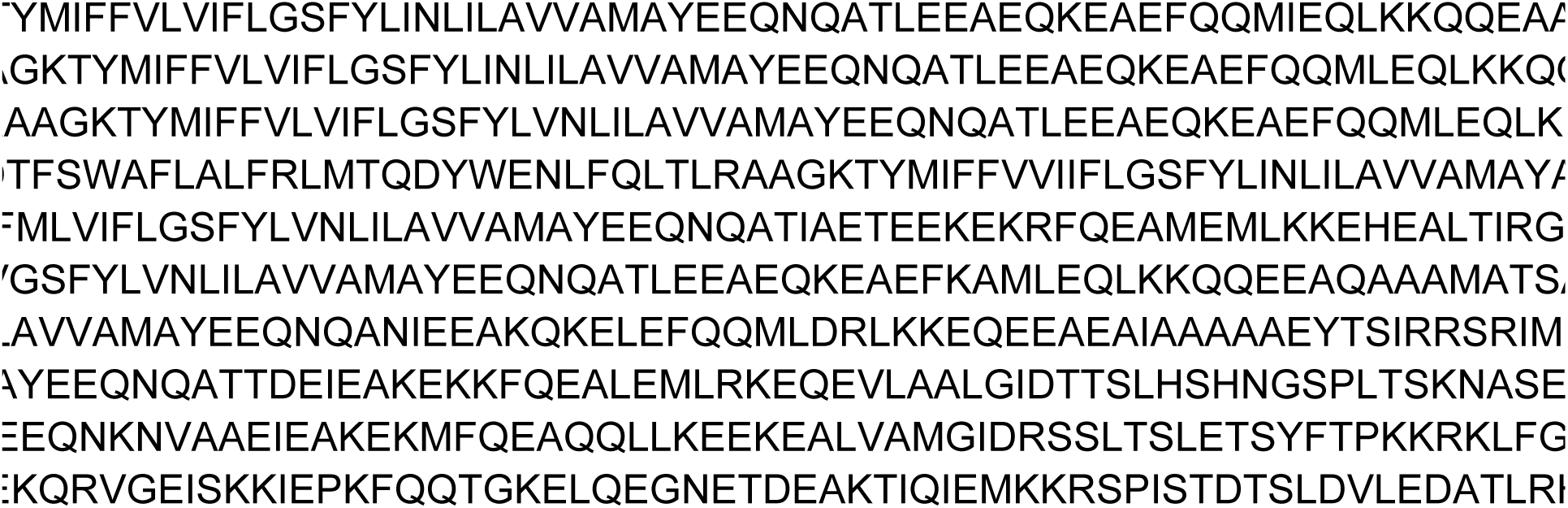
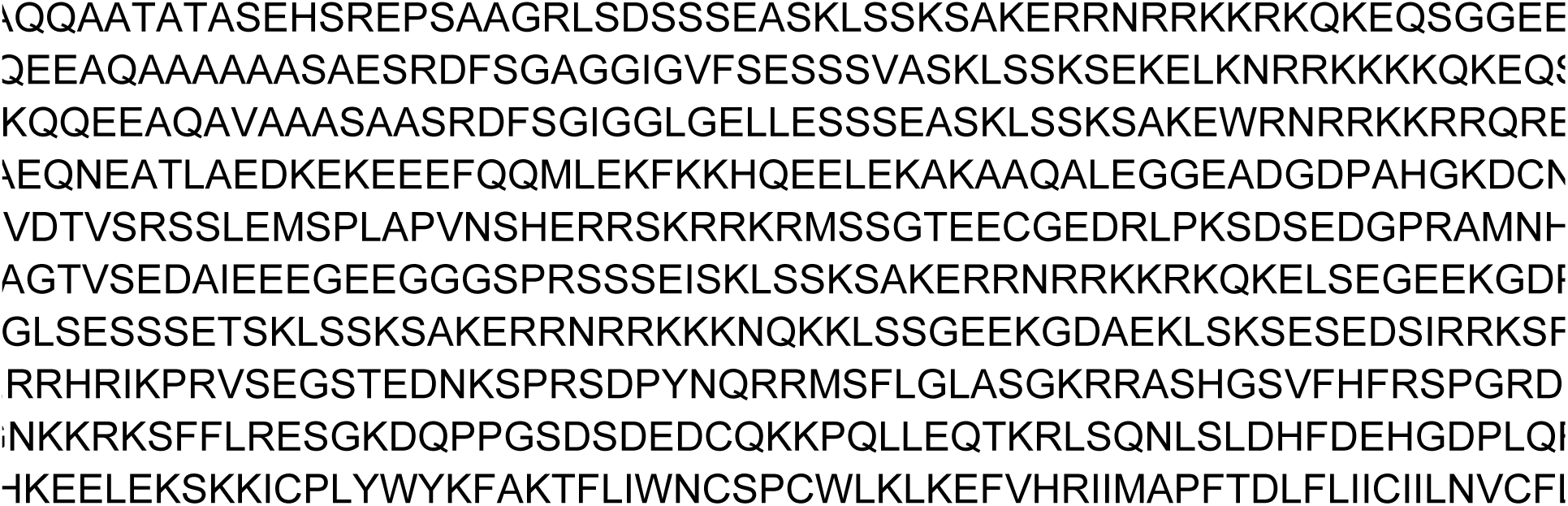
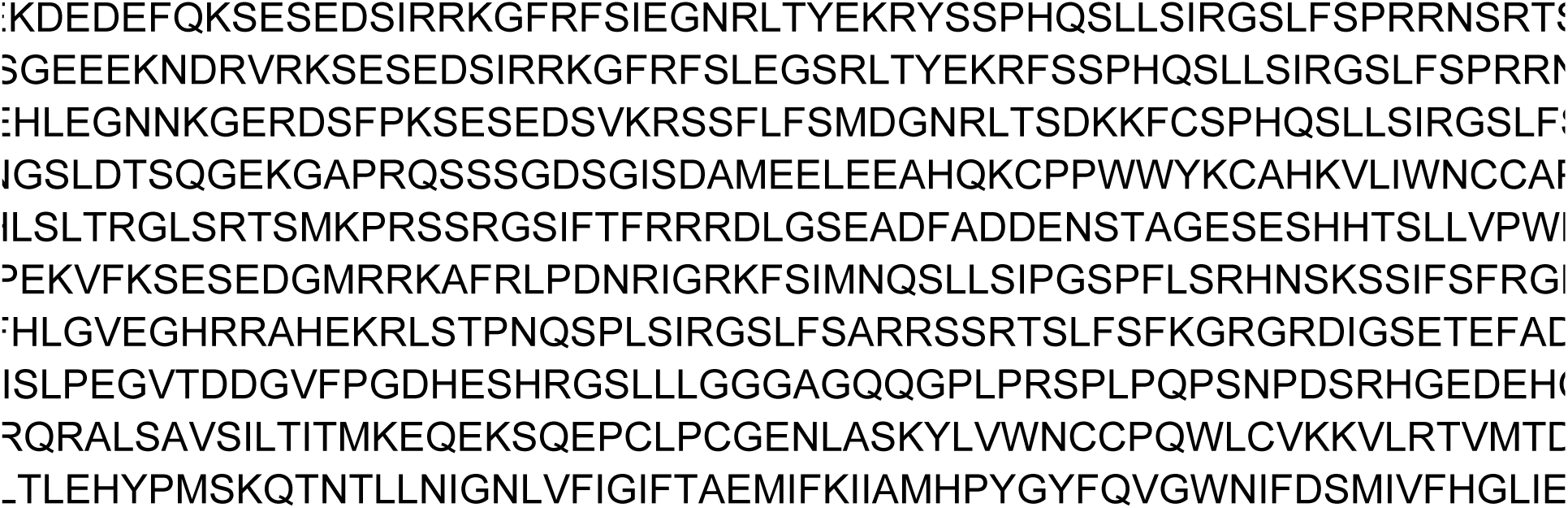
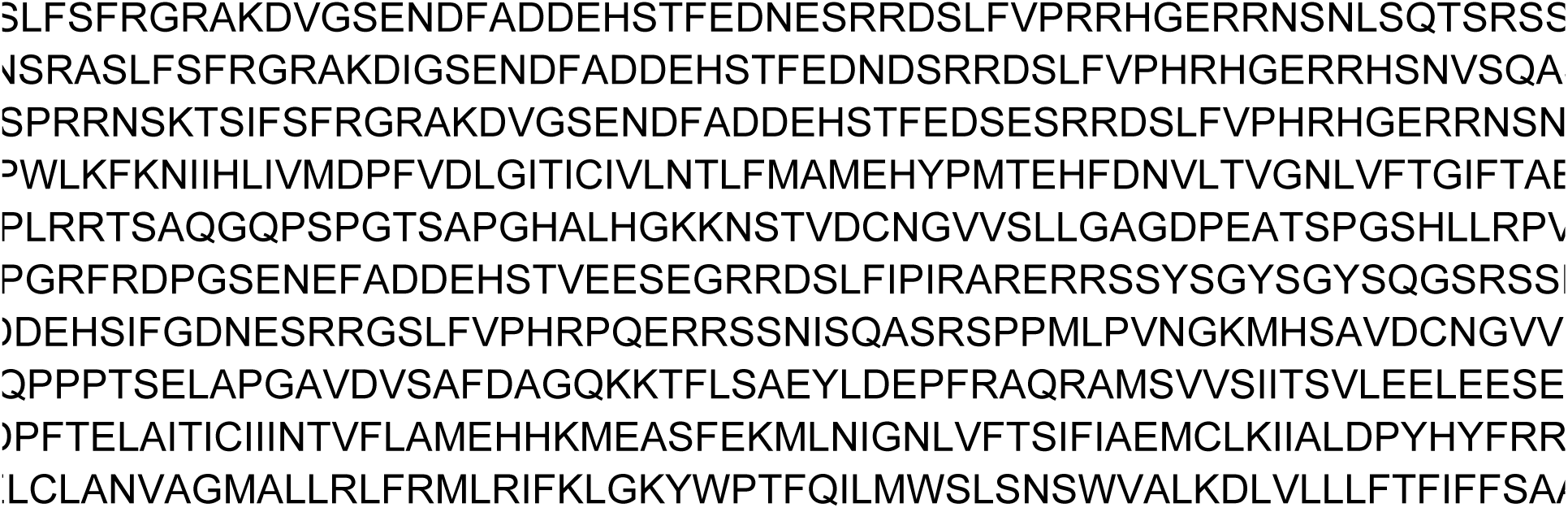
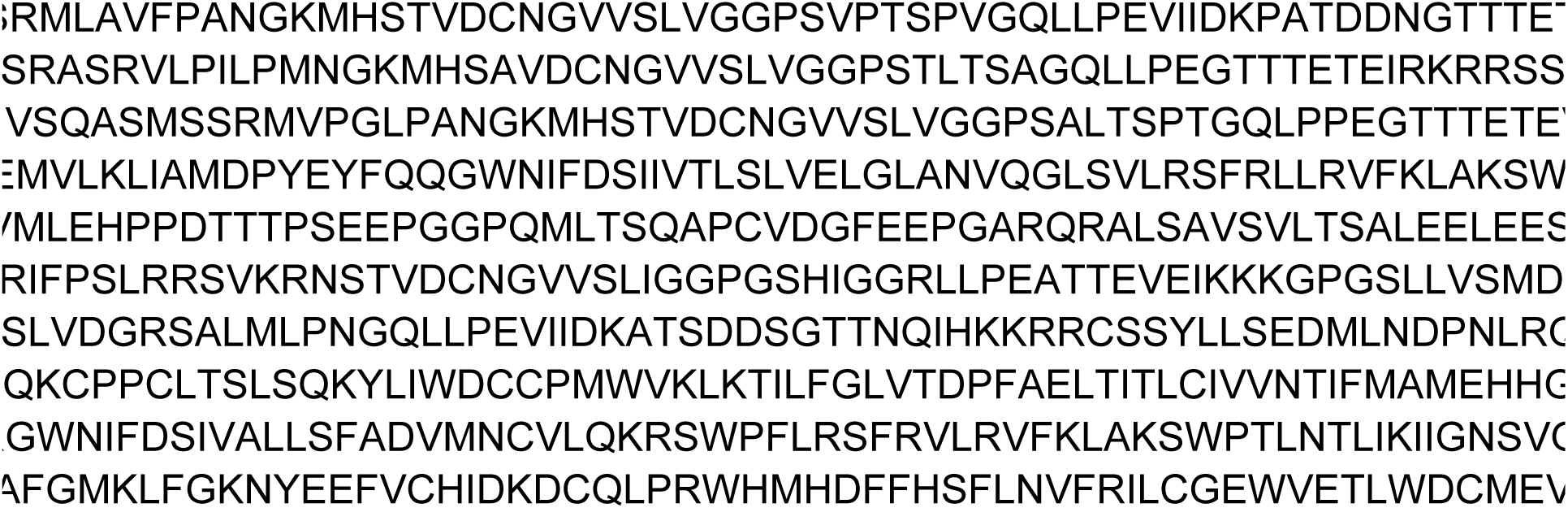
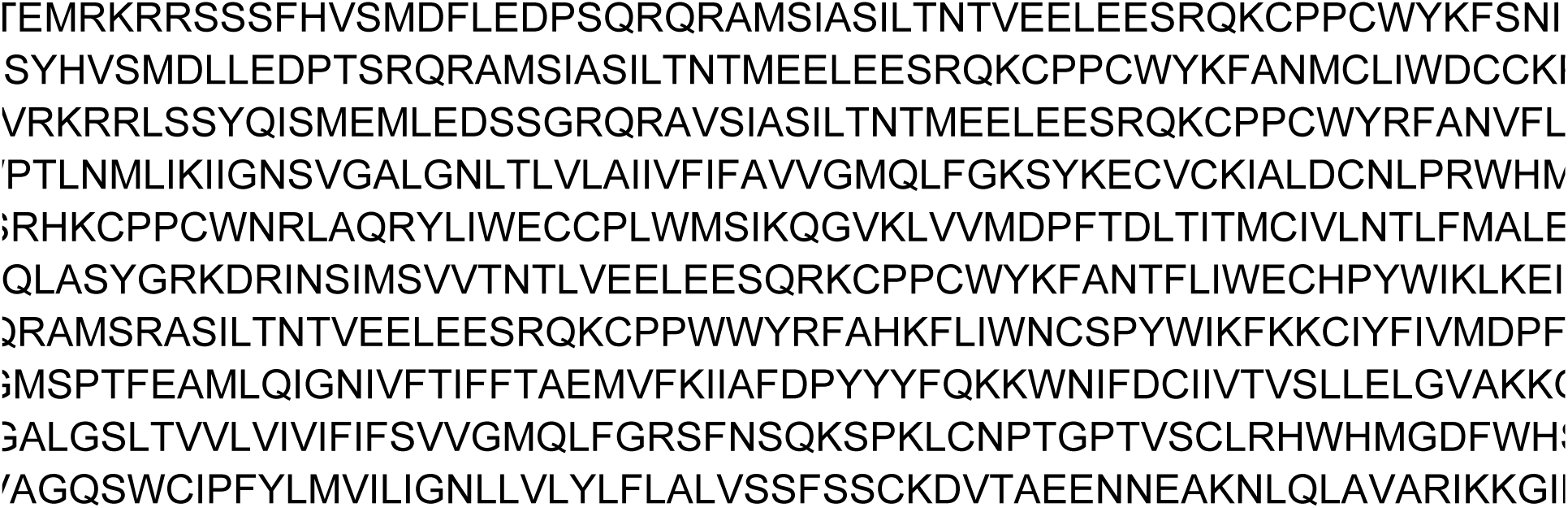
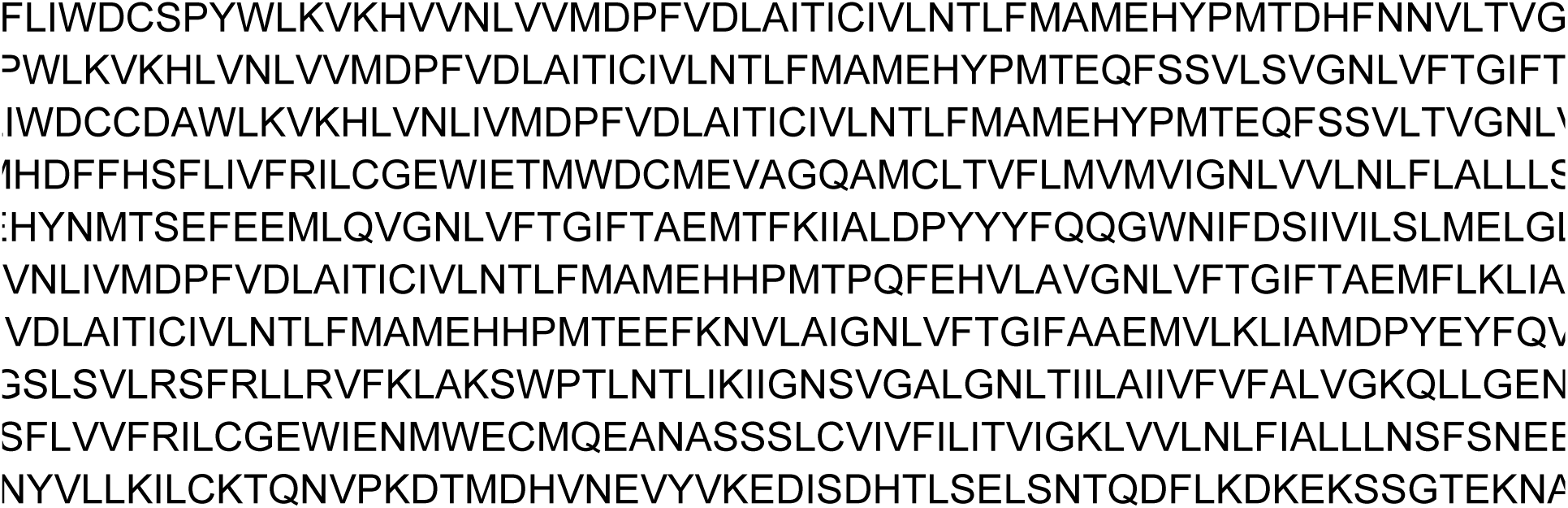
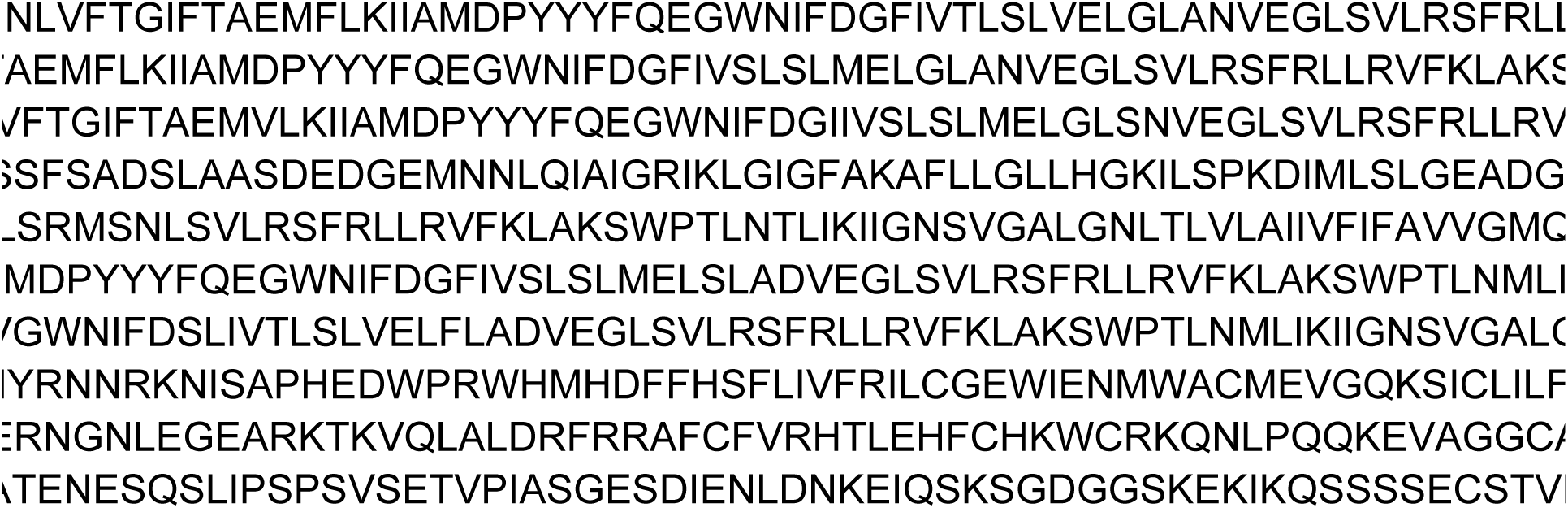
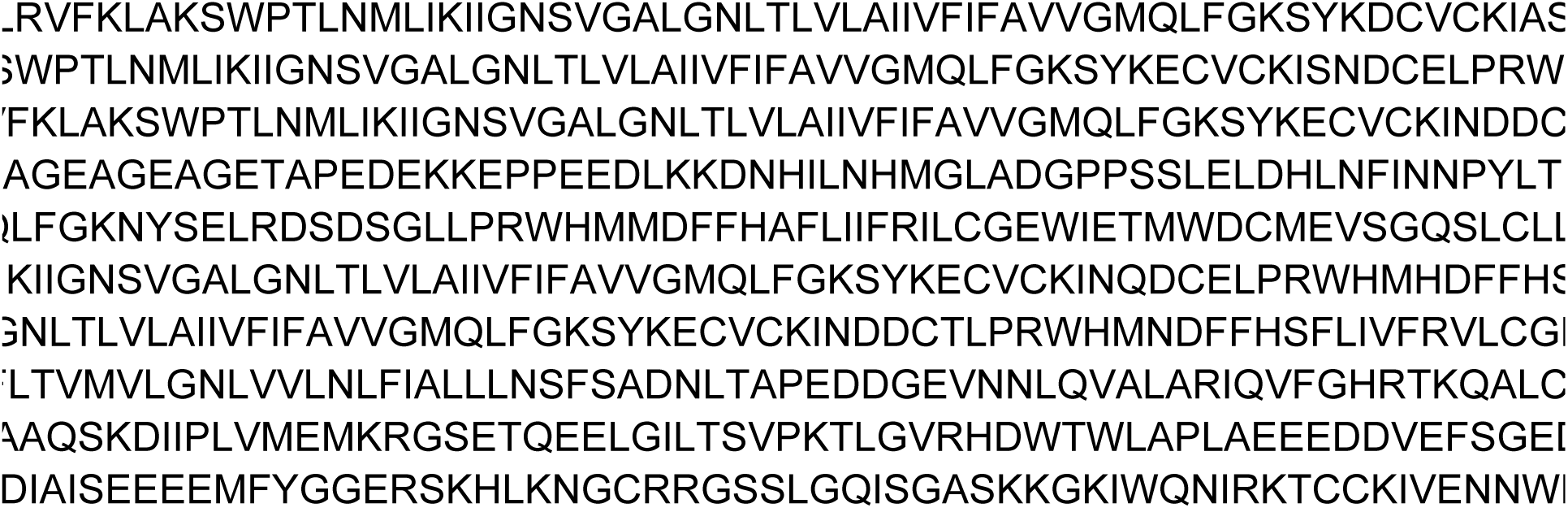
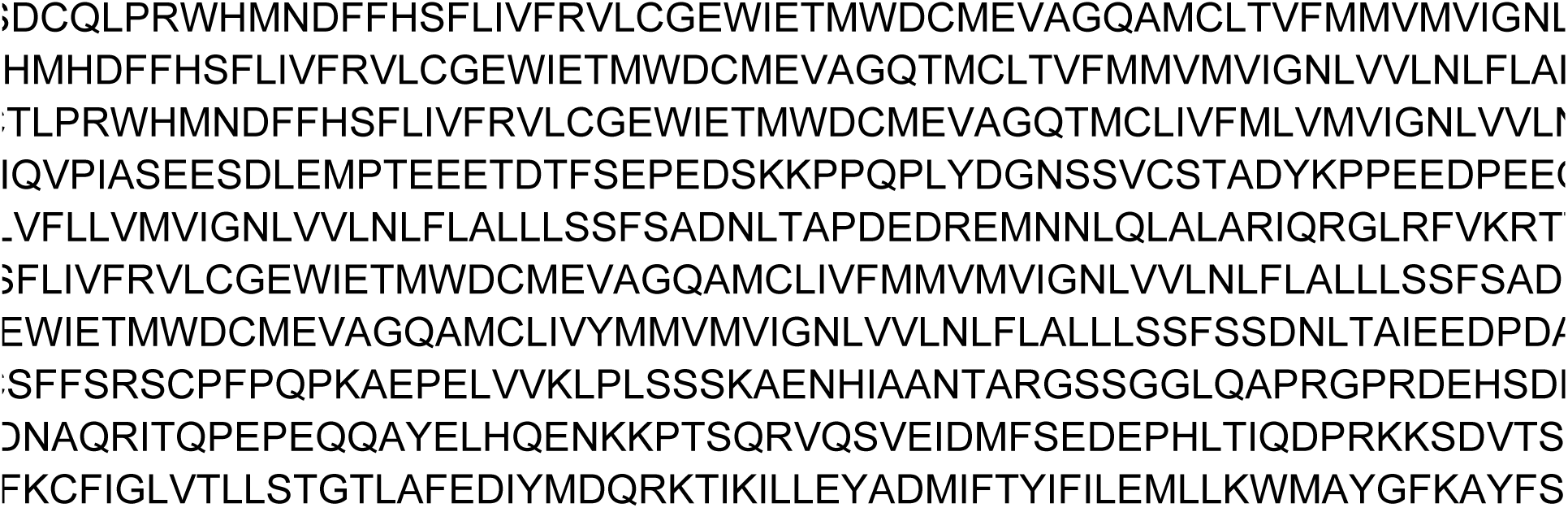
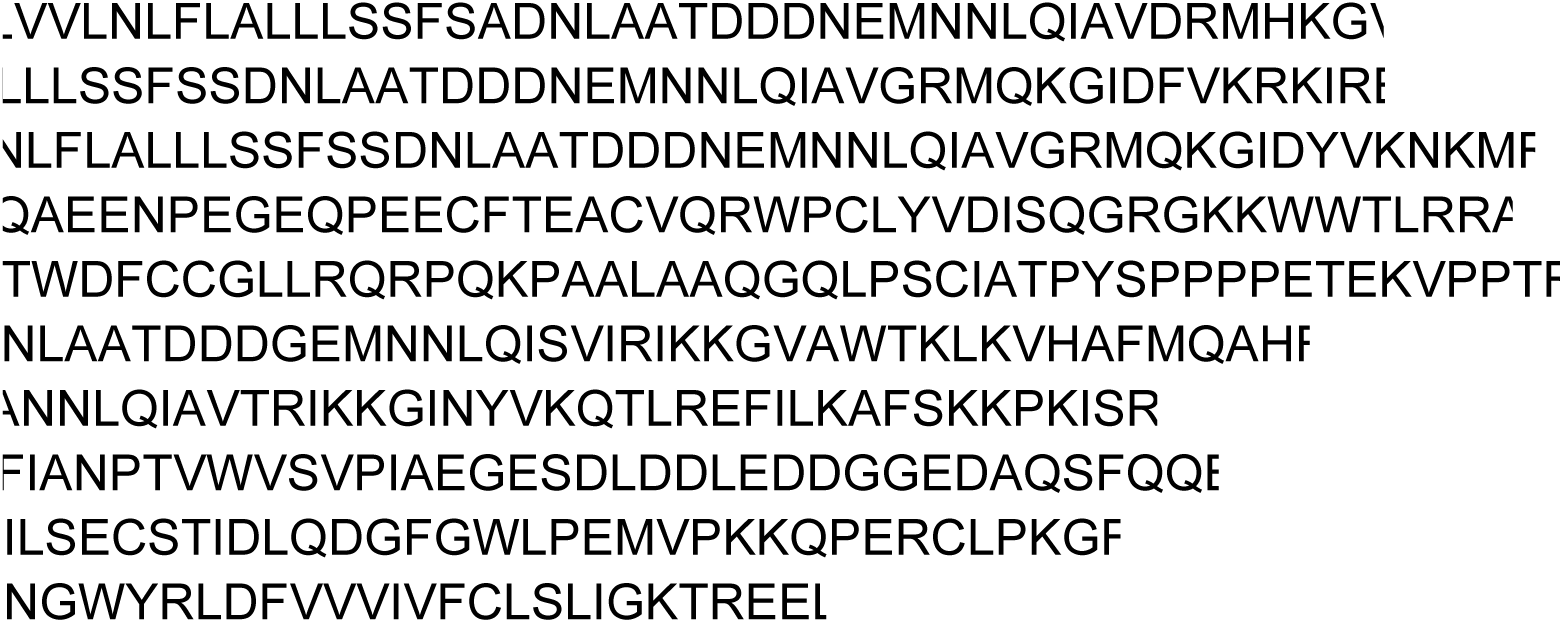
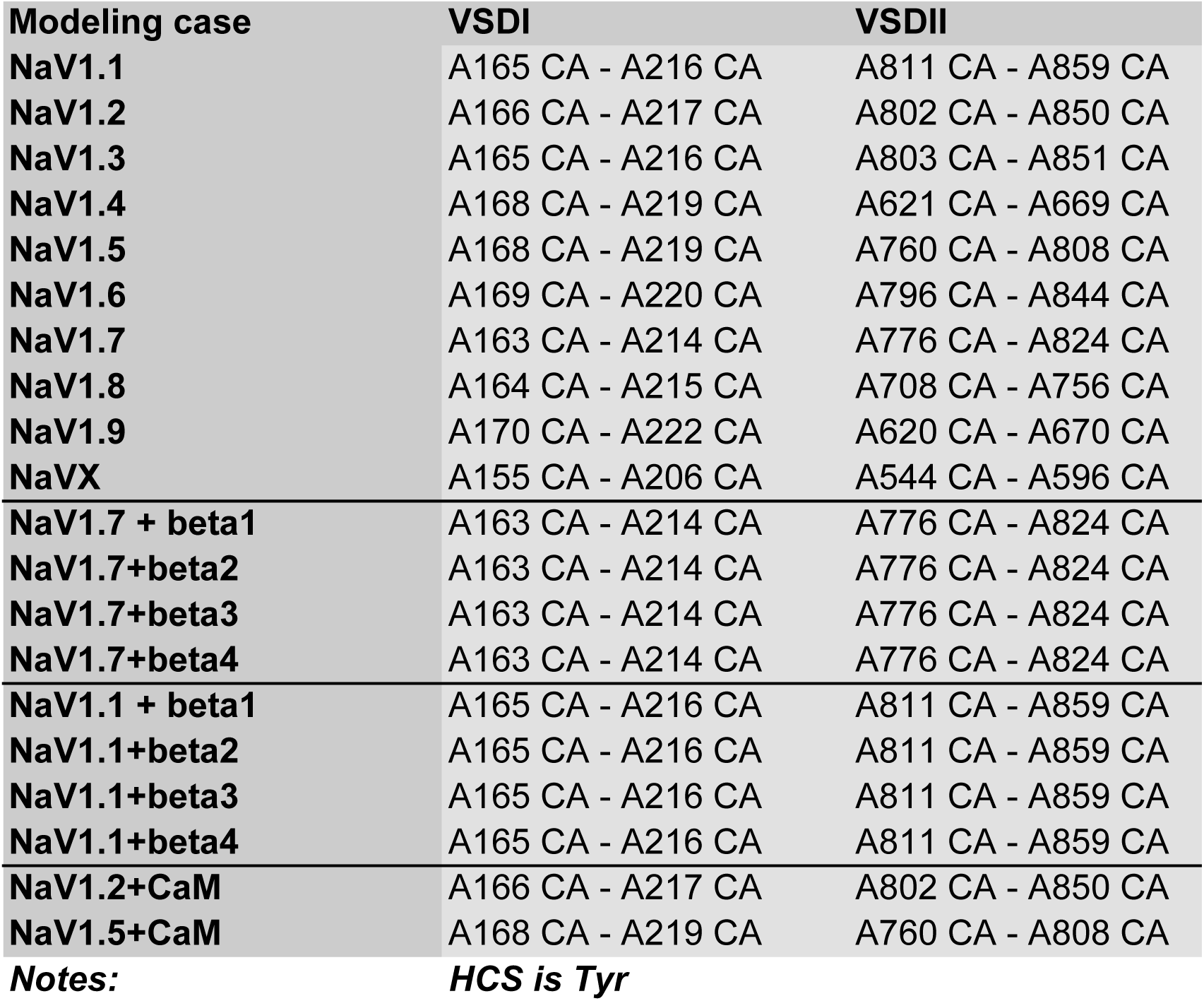
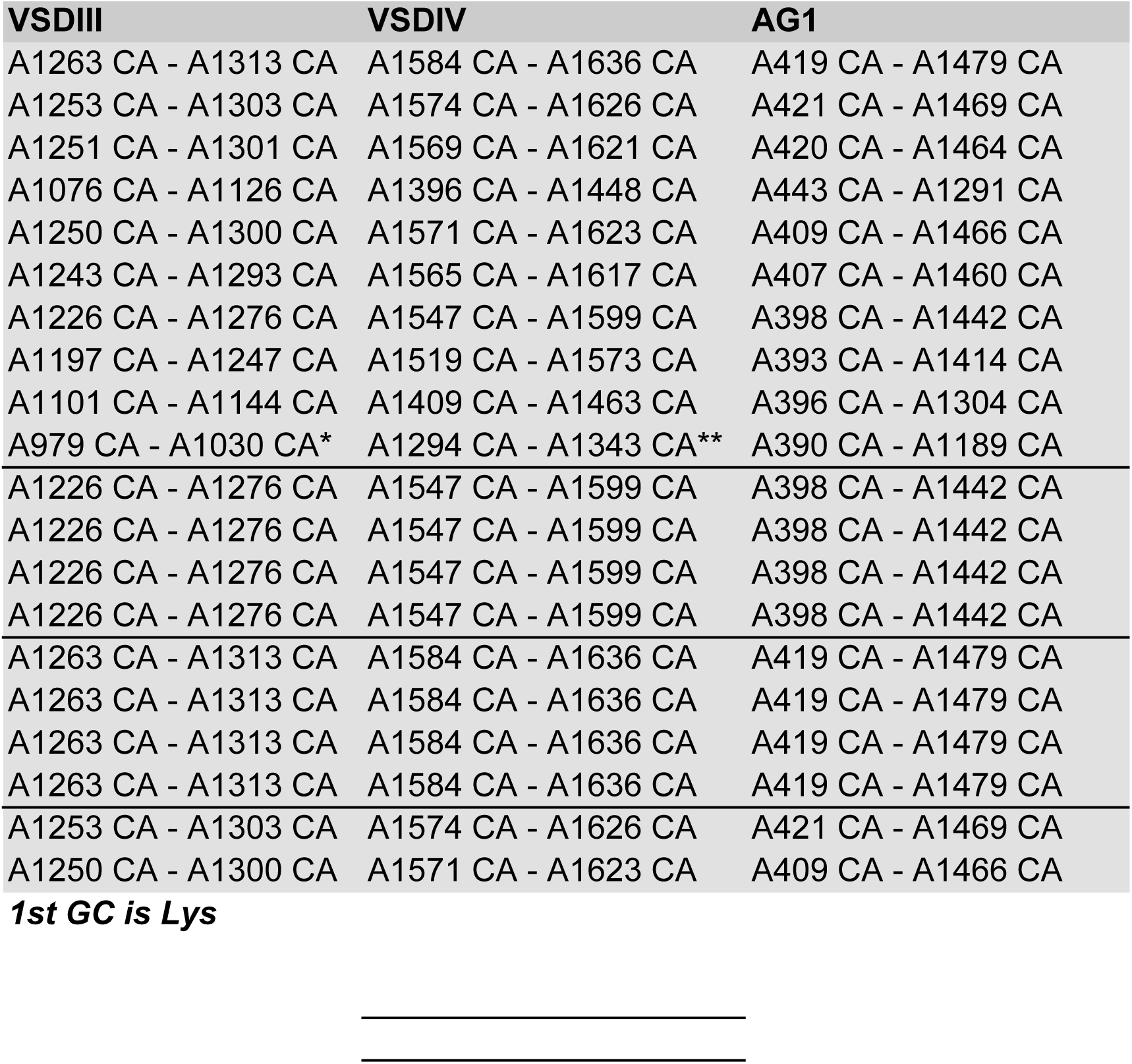
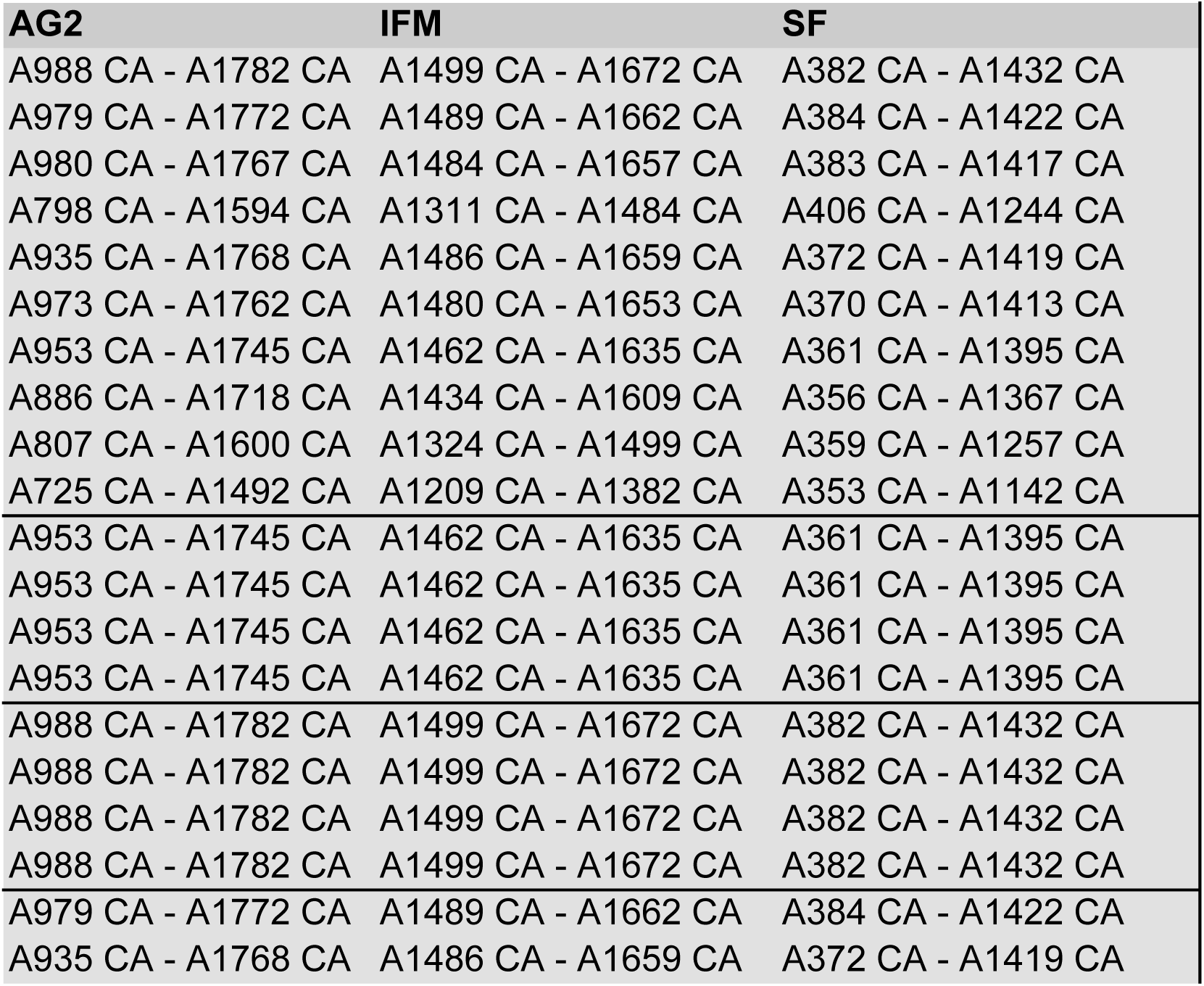
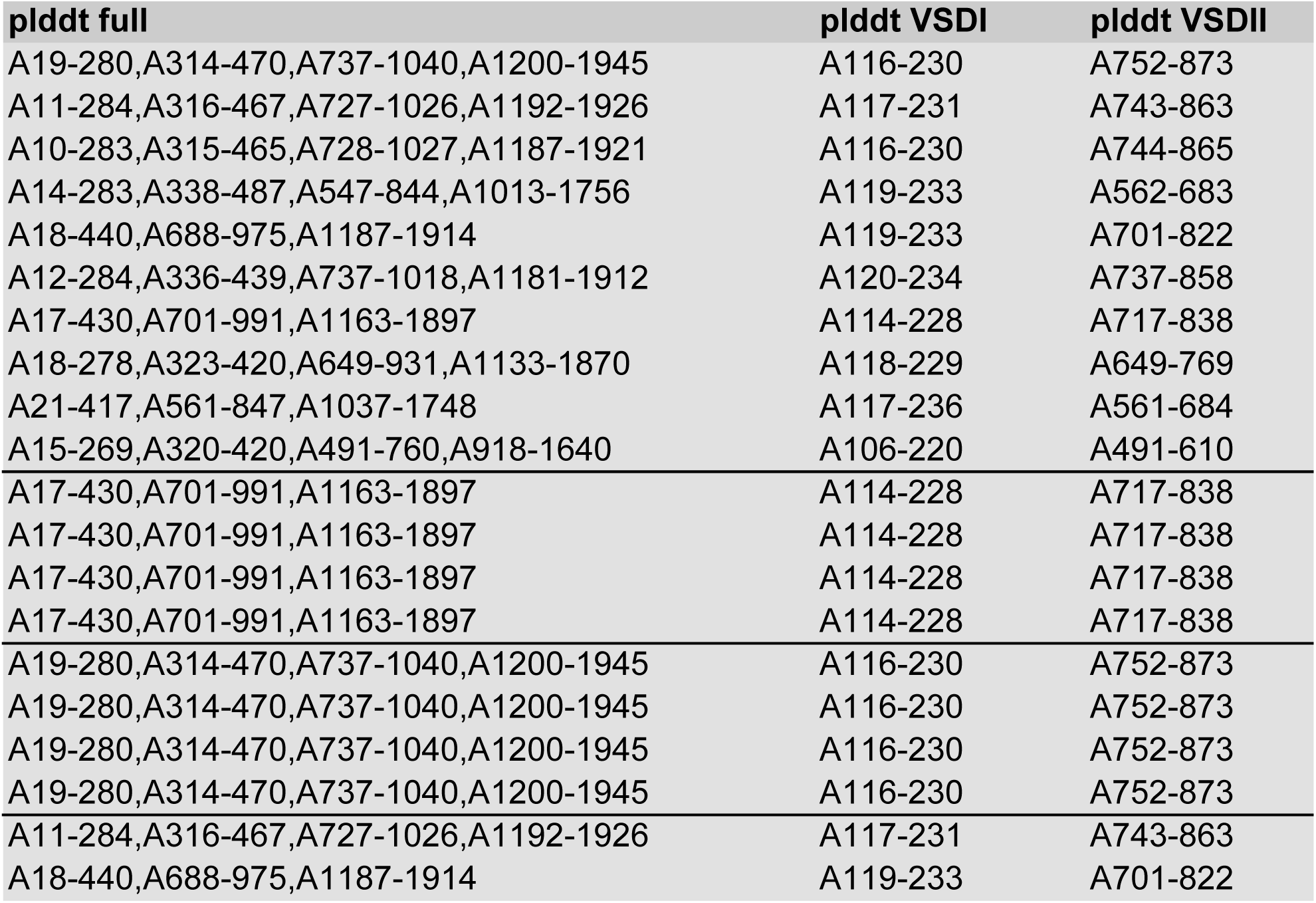
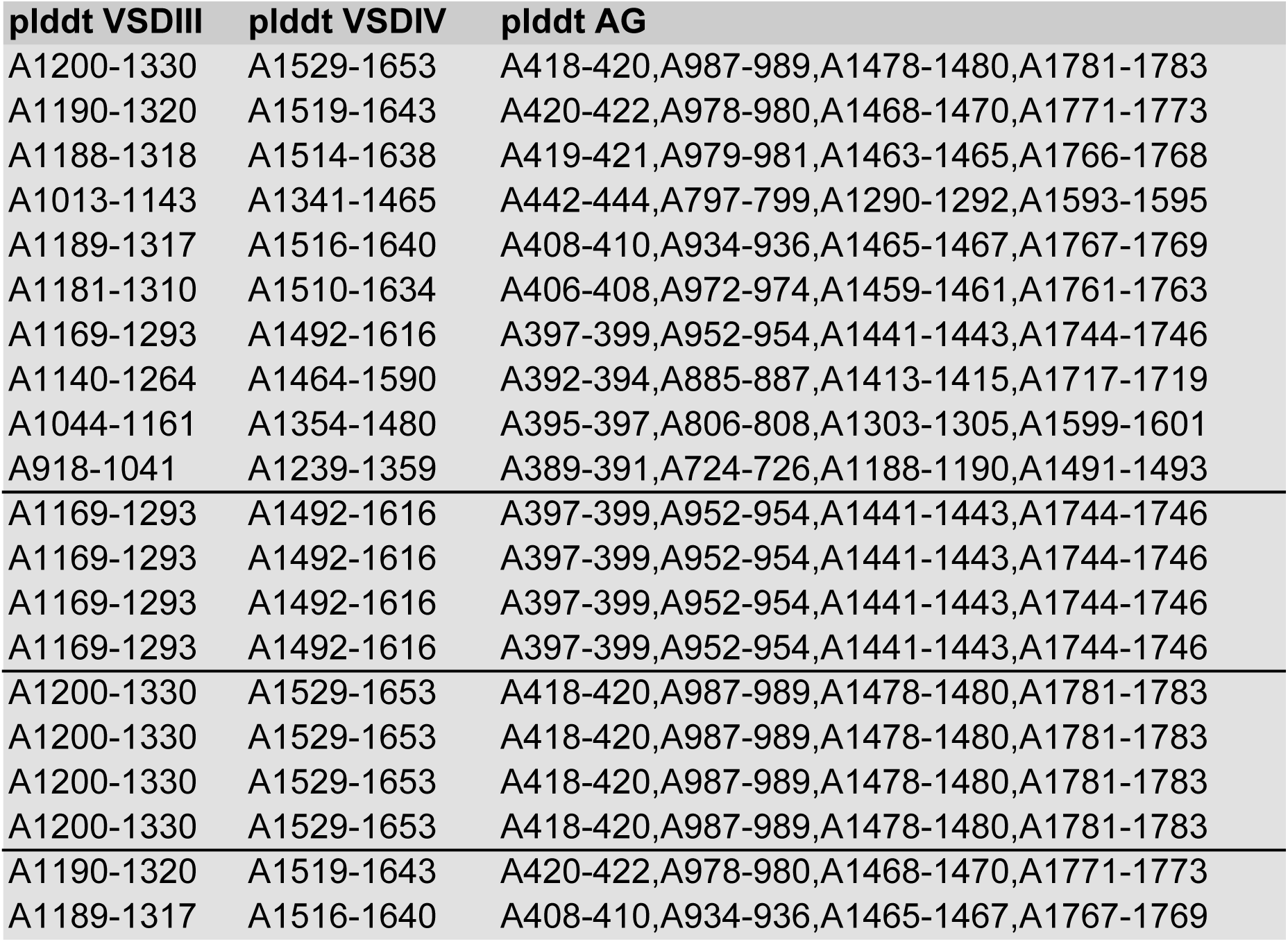
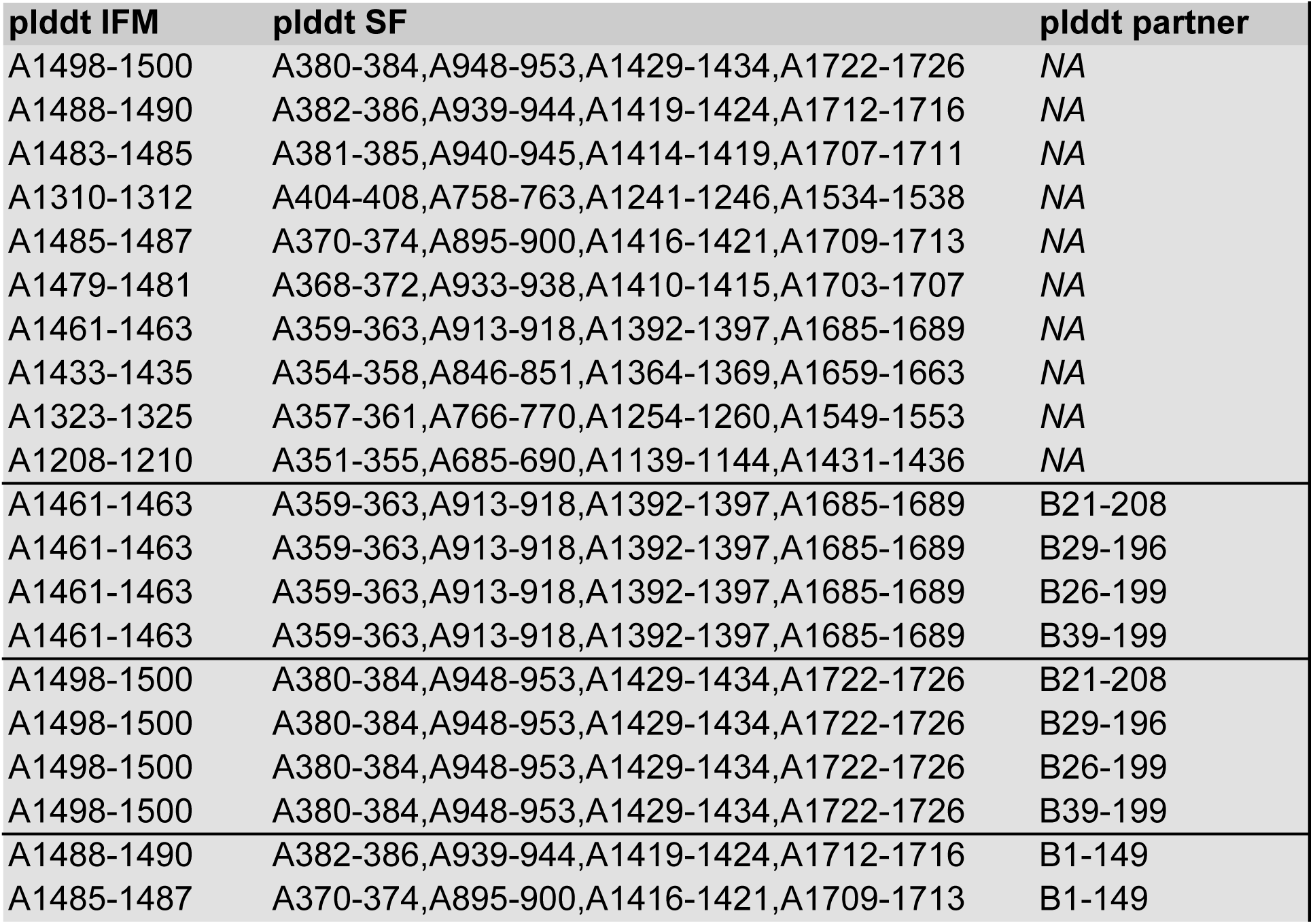
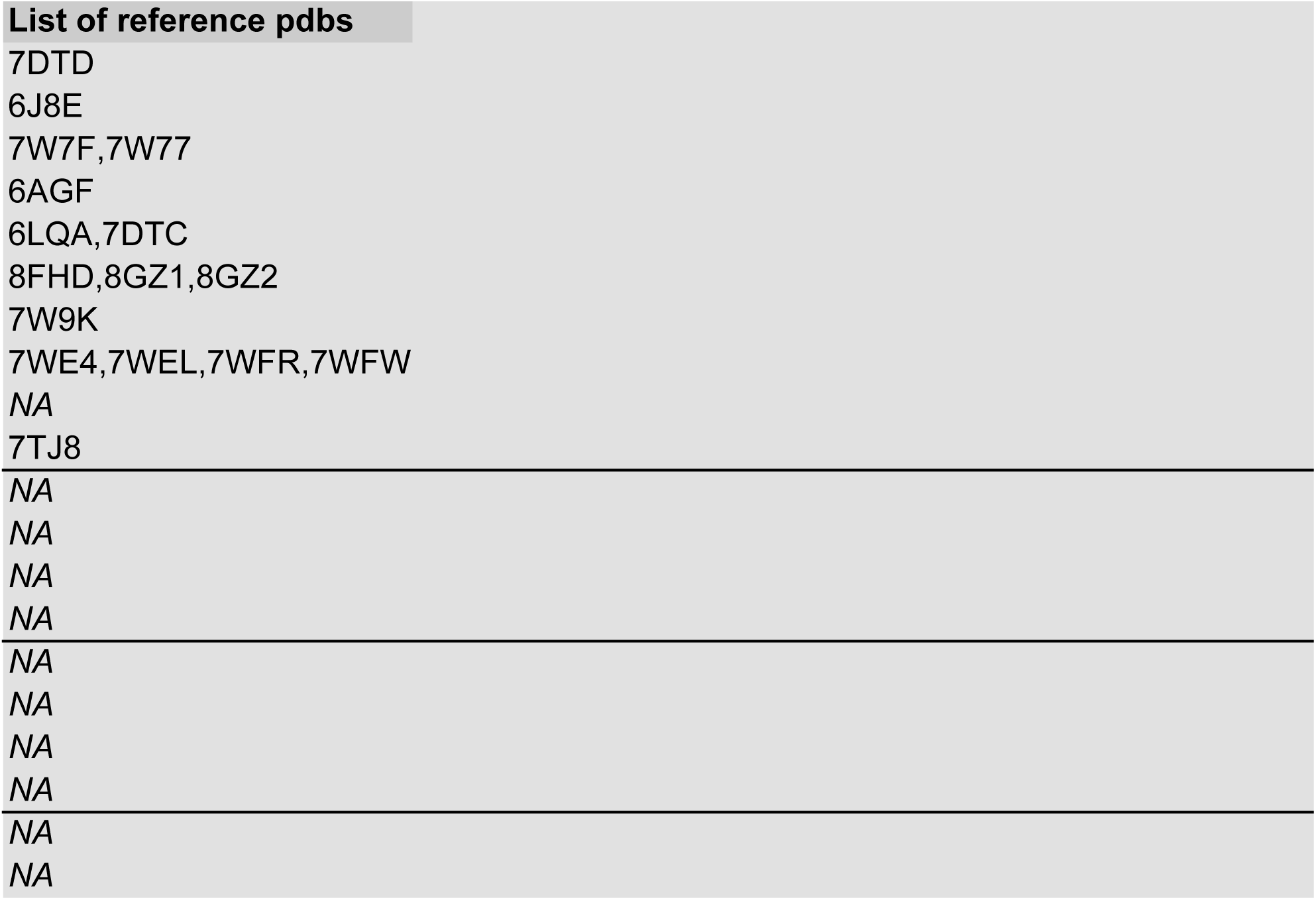
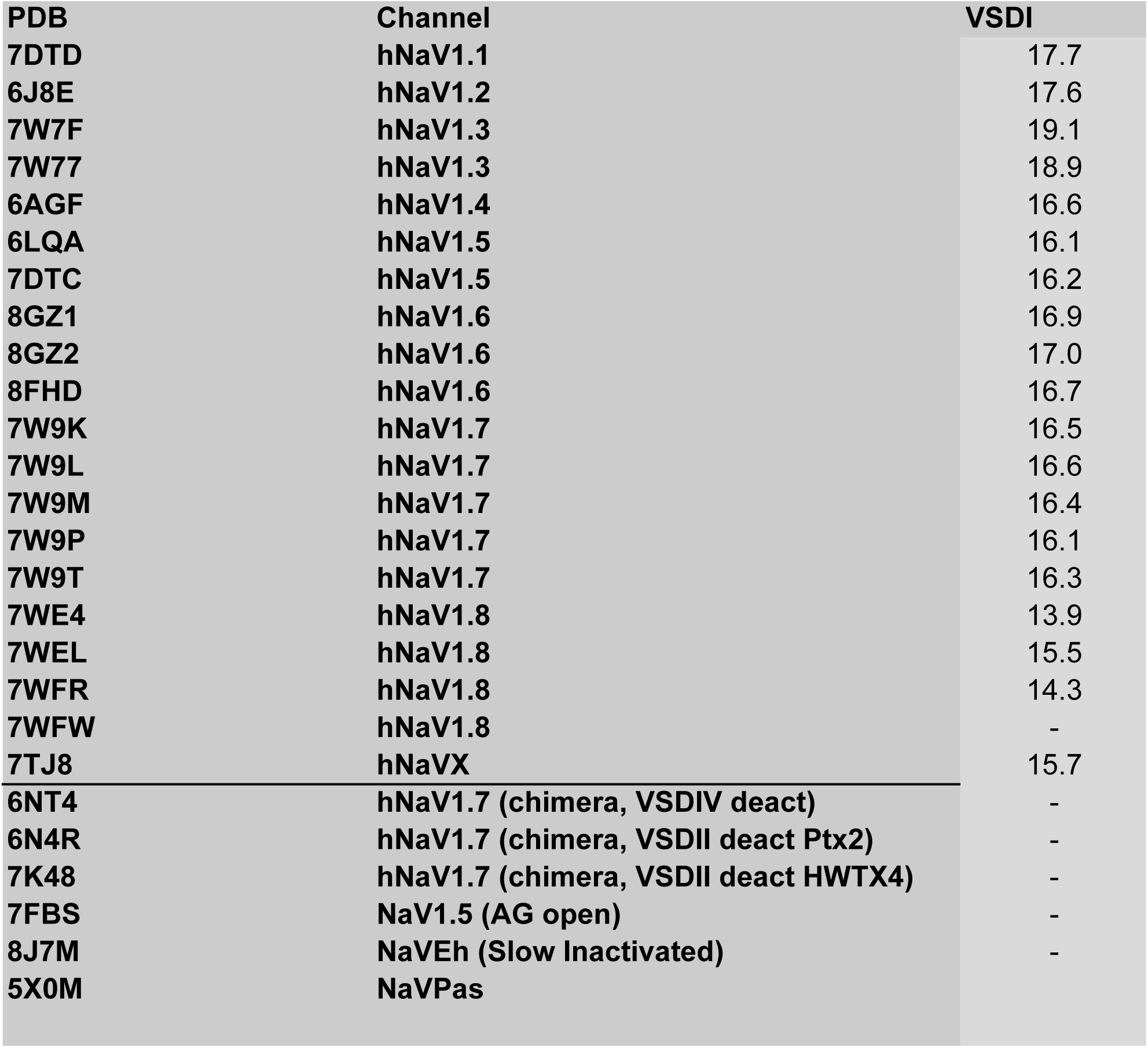
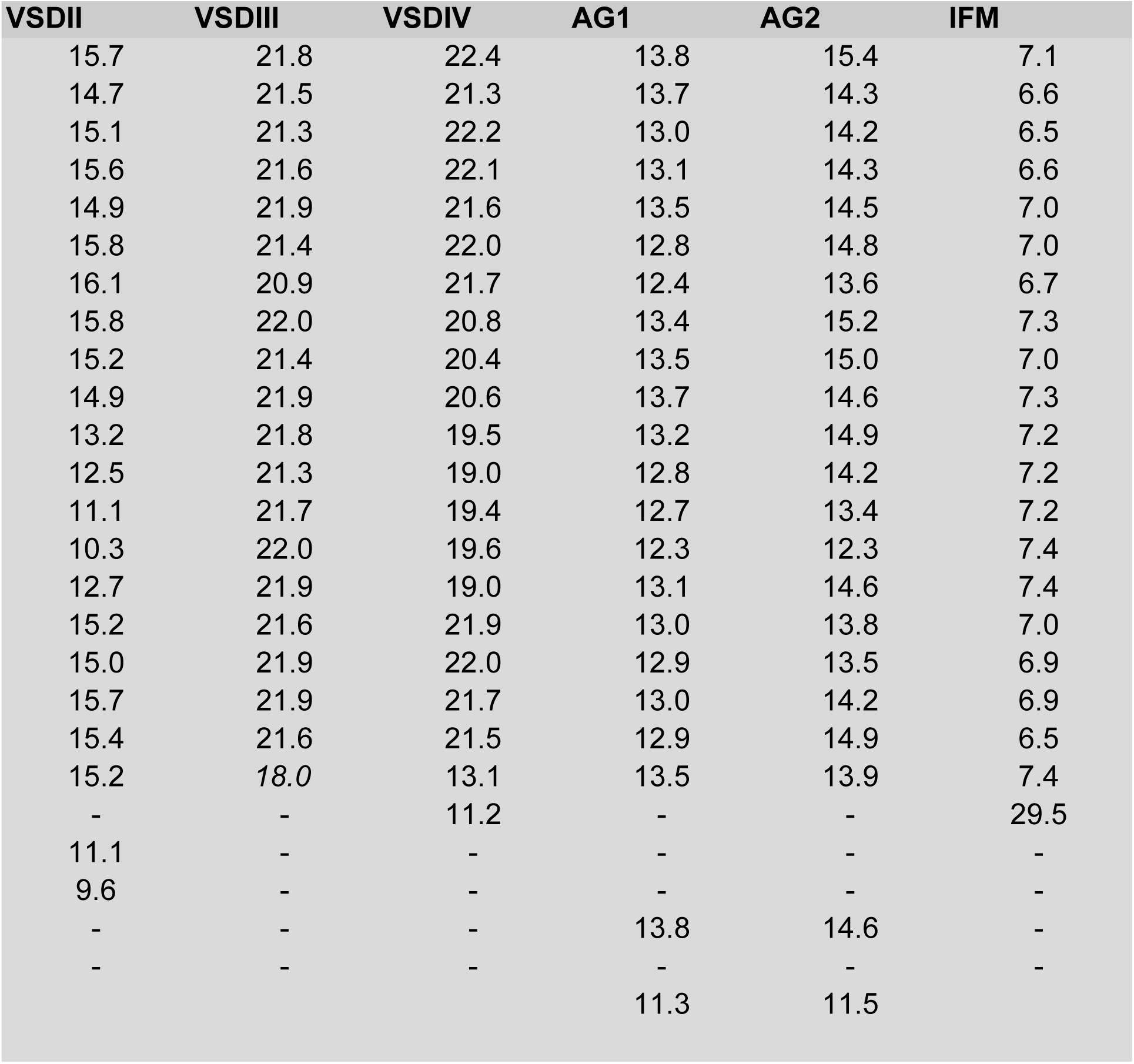
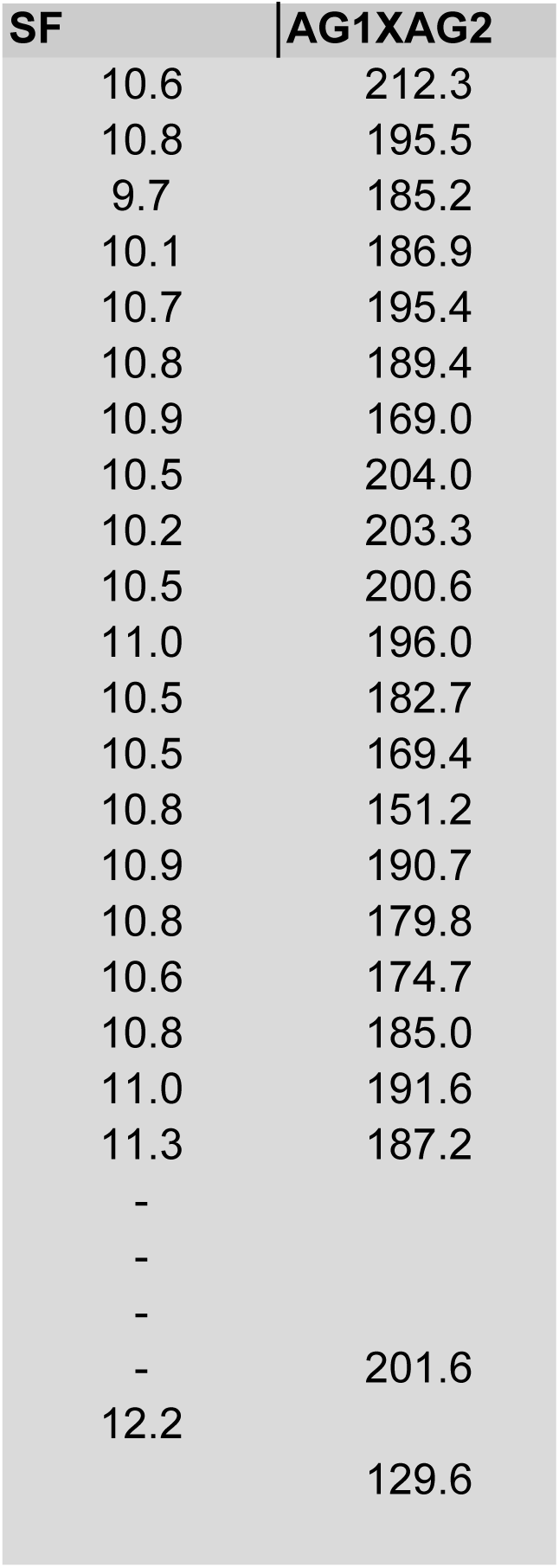

